# Structure and specialization of mycorrhizal networks in phylogenetically diverse tropical communities

**DOI:** 10.1101/2022.05.10.491376

**Authors:** Benoît Perez-Lamarque, Rémi Petrolli, Christine Strullu-Derrien, Dominique Strasberg, Hélène Morlon, Marc-André Selosse, Florent Martos

**Affiliations:** Institut de Systématique, Évolution, Biodiversité (ISYEB), Muséum national d’histoire naturelle, CNRS, Sorbonne Université, EPHE, UA, CP39, 57 rue Cuvier 75 005 Paris, France; Institut de biologie de l’École normale supérieure (IBENS), École normale supérieure, CNRS, INSERM, Université PSL, 46 rue d’Ulm, 75 005 Paris, France; Science Group, The Natural History Museum, Cromwell Road, SW7 5BD London, UK; Peuplements Végétaux et Bioagresseurs en Milieu Tropical, Université de La Réunion, UMR PVBMT, 97 400 Saint-Denis, La Réunion, France; Department of Plant Taxonomy and Nature Conservation, University of Gdansk, Wita Stwosza 59, 80-308 Gdansk, Poland

**Keywords:** common mycorrhizal networks, endomycorrhiza, root endophytism, Mucoromycotina fine root endophytes, early-diverging plants, fungal metabarcoding.

## Abstract

**Background:** The root mycobiome plays a fundamental role in plant nutrition and protection against biotic and abiotic stresses. In temperate forests or meadows dominated by angiosperms, the numerous fungi involved in root symbioses are often shared between neighboring plants, thus forming complex plant-fungus interaction networks of weak specialization. Whether this weak specialization also holds in rich tropical communities with more phylogenetically diverse sets of plant lineages remains unknown.

We collected roots of 30 plant species in semi-natural tropical communities including angiosperms, ferns, and lycophytes, in three different habitat types on La Réunion island: a recent lava flow, a wet thicket, and an ericoid shrubland. We identified root-inhabiting fungi by sequencing both the 18S rRNA and the ITS2 variable regions. We assessed the diversity of mycorrhizal fungal taxa according to plant species and lineages, as well as the structure and specialization of the resulting plant-fungus networks.

**Results:** The 18S and ITS2 datasets are highly complementary at revealing the root mycobiota. According to 18S, *Glomeromycotina* colonize all plant groups in all habitats forming the least specialized interactions, resulting in nested network structures, while *Mucoromycotina* (*Endogonales*) are more abundant in the wetland and show higher specialization and modularity compared to the former. According to ITS2, mycorrhizal fungi of *Ericaceae* and *Orchidaceae*, *namely Helotiales*, *Sebacinales*, and *Cantharellales*, also colonize the roots of most plant lineages, confirming that they are frequent endophytes. While *Helotiales* and *Sebacinales* present intermediate levels of specialization, *Cantharellales* are more specialized and more sporadic in their interactions with plants, resulting in highly modular networks.

**Conclusions:** This study of the root mycobiome in tropical environments reinforces the idea that mycorrhizal fungal taxa are locally shared between co-occurring plants, including phylogenetically distant plants (*e.g.* lycophytes and angiosperms), where they may form functional mycorrhizae or establish endophytic colonization. Yet, we demonstrate that, irrespectively of the environmental variations, the level of specialization significantly varies according to the fungal lineages, probably reflecting the different evolutionary origins of these plant-fungus symbioses. Frequent fungal sharing between plants questions the roles of the different fungi in community functioning and highlights the importance of considering networks of interactions rather than isolated hosts.

## Background

The fungal root microbiome, or root mycobiome, plays a fundamental role in the functioning of plant organisms: it contributes to their nutrition, improves their protection, and fosters their development [1–5]. Among plant-associated fungi, mycorrhizal fungi colonizing the roots of most plant species on Earth supply the plants with minerals gathered from the soil in exchange for plant-assimilated carbon [5–7]. Several categories of mycorrhizae have been proposed based on the morphological structure they form inside plant roots and on the identity of the fungal and plant lineages involved [3, 5]. Currently colonizing >70% of land plants, the *Glomeromycotina* and *Mucoromycotina* (*Endogonales*) subphyla form arbuscular mycorrhizae and coil endomycorrhizae respectively [5,7–9]. Other types of mycorrhizae involve fungal lineages among the *Basidiomycota* division, like the orders *Sebacinales* and *Cantharellales*, or the *Ascomycota*, like the orders *Helotiales* and *Pezizales*. In particular, ectomycorrhizae are present in >13 lineages of land plants [10, 11] and *Ericaceae* and *Orchidaceae* develop specific mycorrhizae [5,7,12]. Although new associations continue to be discovered [12–14], mycorrhizal associations of angiosperms have been thoroughly described in the past decades [7]. In contrast, little is known about the fungal associations of early-diverging plants [5, 15]. For instance, the *Lycopodiaceae* (or clubmosses), a vascular plant lineage that emerged *ca.* 350 million years ago [16], were mainly thought to interact with *Glomeromycotina* fungi [17], until recent studies demonstrated that some lycopod species may also interact with *Mucoromycotina* or *Basidiomycota* [8,18,19].

Besides ‘true’ mycorrhizal associations, advances in DNA sequencing technology have also revealed that many fungi often colonize plant tissues in an ‘endophytic niche’ [20, 21], *i.e.* as biotrophic organisms loosely colonizing plant tissues without forming any visible mycorrhiza [22]. There is therefore an endophytic continuum between fully functional mycorrhizal interactions and saprophytic colonizations [12]. For instance, herbaceous plants that typically interact with *Glomeromycotina* fungi have been found to be also colonized by ectomycorrhizal fungal lineages [23]. These endophytic fungi can importantly contribute to plant nutrition and protection [24, 25]. However, without proper experimental evidence [26], such colonizations by mycorrhizal fungal lineages are not informative on the functionality of these interactions (are there any nutritional exchanges or any protective effect? [8]). Therefore, the impact of an endophyte on its host plant remains generally unknown.

In local communities, *i.e.* at small spatial scales, interactions between plants and mycorrhizal fungi are often not only one-to-one. Plant-fungus interactions rather form a complex and dense network linking different plant taxa and fungi of various lineages [27, 28]. The mycorrhizal fungi of a given plant are indeed often shared with some surrounding plants, where they can either form active mycorrhizae or simply colonize as root endophytes [29, 30]. These interconnected mycorrhizal networks can allow the movement of carbohydrates between plants [31–33] or even inter-individual communication [34]. Many studies have investigated these plant-fungus mycorrhizal networks and have revealed that plant-fungus interactions are non-random [35–38]. Additionally, their specificity ranges from moderately specific interactions, resulting in a lot of fungal sharing between plant species [35,39,40], to very specific interactions [41, 42]. Some studies have also suggested that the structure of the plant-fungus networks varies according to the environmental conditions [3, 43] and the fungal lineages. For instance, plant-*Helotiales* interactions seem to be less specialized than plant-*Cantharellales* interactions [40, 44].

Studies looking at plant-fungus interaction networks have mostly focused on temperate communities dominated by angiosperms, with a few, if any, representing taxa of “early-diverging” plant lineages, such as bryophytes, ferns, or lycopods. Whether plant lineages that diverged hundreds of million years ago are homogenously sharing similar fungi, or whether plant-associated fungi tend to segregate because of different nutrition strategies, spatial heterogeneity, ecological preferences, and/or evolutionary constraints have often been debated [15]. Yet, the extent of fungal sharing between distantly related plant lineages has rarely been measured in local communities and remains unclear. On a global scale, in a recent meta-analysis of plant-*Glomeromycotina* interactions [41], some lycopod species appeared to specifically interact with a distinct clade of *Glomeromycotina*, forming a separate module of interaction, which is very unusual in arbuscular mycorrhizal symbioses. However, whether this specialization in lycopod-fungus interactions also exists in local communities is unclear.

Here, we studied plant-fungus interactions in three local communities including various plant taxonomic groups (bryophytes, lycopods, ferns, and angiosperms) across the tropical island of La Réunion. We investigated (i) what are the main mycorrhizal fungal lineages colonizing phylogenetically distant plant taxa in these tropical communities characterized by contrasted environmental conditions and (ii) how these fungi are shared between surrounding plants in each local community. We sampled roots of the main plant species in the three communities and identified the plant-associated fungi by using metabarcoding technics targeting the 18S rRNA and ITS2 fungal markers. We selected all root-associated fungi that may form mycorrhizal interactions with at least one plant species in the communities, reconstructed the networks of interactions between plants and fungi at the local scale, and evaluated the degree of specialization of these plant-fungus interactions across the main plant taxonomic groups and across the different environmental conditions. We particularly focused on the distinctiveness in terms of specialization and network structure of the main fungal lineages. Finally, we specifically assessed whether strong lycopod-fungus specialization exists at the level of each sampled community. Based on previous findings on a global scale [15, 41], we expected to find distinct fungal colonizations across plant species from different taxonomic groups and predicted to observe less fungal sharing between phylogenetically distant plant lineages. This would result in (i) plant lineage-specific fungi, *e.g.* lycopod-specific fungi, (ii) an evolutionary conservatism of plant-fungus interactions, *i.e.* distantly related plant species would interact with less similar sets of partners than closely related ones, and (iii) modular network structures where some subsets of species form separated modules of interactions.

## Material & Methods

### Study sites and sampling

The study was conducted on the tropical volcanic island of La Réunion in July 2019. To maximize the phylogenetic breadth of vascular plant species co-occurring in a sampling site, we chose three plant communities containing lycopods (clubmosses), ferns and/or bryophytes, and angiosperms across contrasted habitats described in [45]. The first community, in Grand brûlé (S21°16’39’’, E55°47’29’’), is derived from a recent lava flow that occurred in the 19^th^ century. The soil in Grand brûlé is therefore shallow, poorly differentiated, and colonized by many non-indigenous invasive plant species (Supplementary Table 1). This community is typical of the earliest stages of ecological succession following a disturbance. Near the sea and at a low elevation (100 meters), Grand brûlé has frequent precipitations (annual rainfall of 4,000-5,000 mm). The second community, in Plaine-des-Palmistes, is a wet area very dense in shrubs, called a ‘thicket’, in the central valley (S21°07’08’’, E55°38’36’’, elevation 900 meters). The soil in Plaine-des-Palmistes is derived from old lava flows that happened between 80,000 and 20,000 years ago. The very frequent precipitations in Plaine-des-Palmistes (annual rainfall of 5,000-7,000 mm) generate many ponds in these thickets and greatly leach nutrients out of the soil. The third community, in Dimitile, is an ericoid shrubland formed on the high-elevation ridge of Cilaos Circus, dominated by the endemic species *Erica reunionensis* (S21°16’39’’, E55°47’29’’, elevation 2,000 meters). Compared with the two previous communities, Dimitile experiences less precipitation (annual rainfall of 1,000-2,000 mm). The soil in Dimitile, derived from >400,000-year-old lava flows, is particularly enriched in acidic humus. Both plant communities in Plaines-des-Palmistes and Dimitile are mostly composed of native species (Supplementary Table 1). We therefore covered contrasted habitats, especially in terms of soil composition, elevation, disturbance, and humidity (Fig. 1).

**Figure 1:**
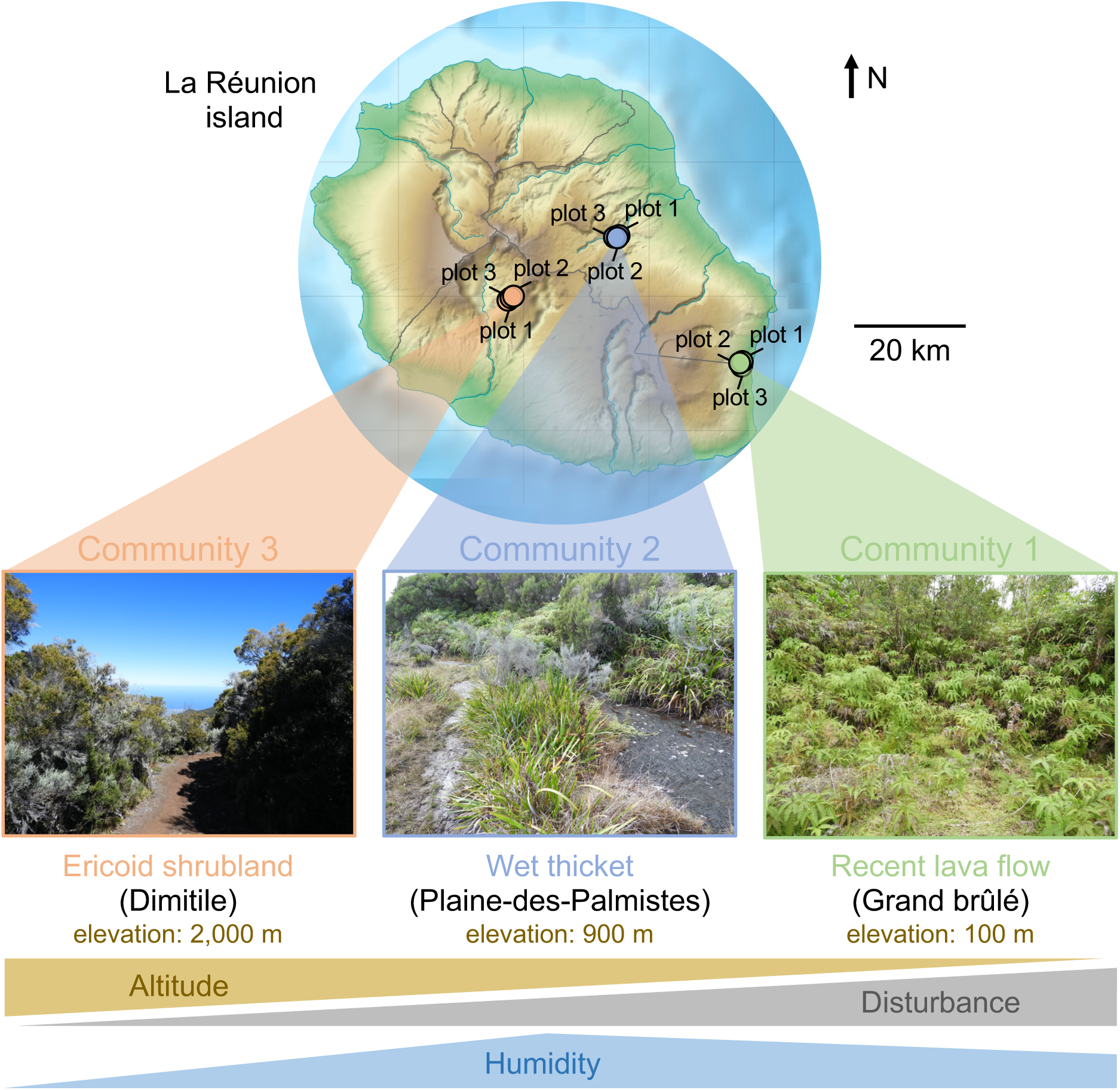
The three sampled communities correspond to habitats with contrasted environmental conditions. A map of La Réunion island indicating the three sampled communities in this study. The sampling sites were characterized by different vegetations and abiotic conditions with different elevations, levels of disturbance, humidity, and soil conditions: (right) Grand brûlé (recent lava flow close to the ocean on the wet East coast), (middle) Plaine-des-Palmistes (wet thicket on old lava flows in the central valley, elevation 900 meters), and (left) Dimitile (ericoid shrubland on old lava flows in the dry crests of Cilaos circus on the West side dominated by ericoid vegetation, elevation 2,000 meters). In each community, we replicated the sampling in three plots distant from 50 meters to 250 meters. The photos illustrate the overall vegetation in each sampled community and the gradients at the bottom resume the main variations in the environments. The raw map in the background was generated by Eric Gaba (Wikimedia Commons user: Sting).

In each community, we sampled 3 plots distant from 50 to 250 meters having similar sets of plant species. As lycopods tend to have patchy distributions in these communities, we specifically targeted zones where they were present. For each plot, we harvested the roots of all the plant species that were present: within a radius of 1.5 meters, we sampled up to 3 individuals per plot per plant species if available. For small non-woody plants, the entire root systems were carefully removed from the soil, cleaned with sterile water, and immediately dried in silica gel. In Dimitile, as the thin colonized roots could not be sampled under large shrubs, *e.g. Erica reunionensis, Phylica nitida,* and *Stoebe passerinoides*, several soil cores were then collected and up to 23 individual roots per plot were amassed without a direct species identification on the field (they were identified using plant DNA sequencing - see next section).

### Molecular analyses and bioinformatics

Dried roots were crushed using sterile tungsten beads in the TissueLyser II (Qiagen). Plant and fungal DNA was extracted from 30 mg of root powder using the NucleoSpin 96 Plant II kit (Macherey-Nagel) following the manufacturer’s instructions. Negative controls were carefully included during DNA extraction [46]. To better characterize the whole fungal diversity that may colonize these different plant species, we amplified two nuclear regions of the rDNA operon: the 18S rRNA gene and the ITS2 region. We used two sets of tagged primers: AMADf-AMDGr [47] and ITS86F-ITS4 [48, 49], that respectively amplify a fragment of 380 and 280 bp on average. The former marker gene rather detects *Glomeromycotina* and *Mucoromycotina*, whereas the latter marker is more specific to *Ascomycota* and *Basidiomycota*. Amplicons pools were carried out as in Taberlet *et al.* (2018) and Petrolli *et al.* (2021) and the resulting DNA amplicons were sequenced using Illumina 2x250 bp MiSeq technology (see Supplementary Methods 1).

The sequencing reads were processed using VSEARCH [52] following the pipeline of [53] (Supplementary Methods 2). In short, paired-end reads were assembled, quality checked, demultiplexed using cutadapt [54], and clustered into operational taxonomic units (OTUs) using two different methods. We first used Swarm [55], a clustering approach that does not rely on a global threshold of similarity but instead uses local thresholds and amplicon abundances. Secondly, we performed a classical 97% OTU clustering using VSEARCH. We removed the chimera in both sets of OTUs and assigned taxonomy to each OTU using Silva and UNITE databases [56, 57] (Supplementary Methods 2). Given that both Swarm and 97% OTU clustering gave qualitatively similar results, we only reported the results obtained with Swarm. We used the decontam pipeline to filter out the contaminants of our OTU tables [58] and evaluated the amount of index hopping and cross-contaminations (Supplementary Methods 3). Finally, the 18S and ITS2 OTUs assigned to plant species were used to identify the roots directly collected in the soil in Dimitile. Non-fungal OTUs were then discarded for subsequent analyses.

We used FUNGuild [59], a program that automatically assigns the possible niches of a fungal OTU based on its taxonomic assignation, and manual filtering, to only retain the putative mycorrhizal OTUs for the rest of the analyses. The few samples having less than 20 reads of these selected OTUs were discarded. We also drew rarefaction curves to see how the fungal diversity within each plant species increases as a function of the number of sampled plant individuals.

### Assessing the similarity in root mycobiome composition across samples

We first assessed the effect of the three sampled communities (Grand brûlé, Plaine-des-Palmistes, and Dimitile) on root mycobiome composition. For this purpose, we used a permutational analysis of variance (PERMANOVA; *adonis* function from the R-package vegan [60] with 10,000 permutations) based on Bray-Curtis dissimilarities to test whether the root compositions were significantly different across the three sampled communities when comparing (i) all the plant species or (ii) only the plant species simultaneously present in several sampled communities. We also visualized the similarities of the root mycobiota compositions by performing hierarchical clustering: We built the dendrogram between samples using neighbor-joining (function *nj* in the R-package ape [61]) based on the Bray-Curtis diversities. All the diversity analyses were replicated using generalized UniFrac distances, a phylogenetically-informed diversity index [62]. Given that this did not qualitatively change our results, we only reported the results obtained with Bray-Curtis dissimilarities in the main text.

Second, we investigated whether, in each local community, the three plots were sufficiently similar to be merged. We performed principal coordinate analyses (PCoA) and PERMANOVA to assess whether samples from the same species but from different plots tend to host similar fungi.

### Measuring the influence of the plant taxonomic groups on plant-fungus interactions

We first measured the level of fungal sharing between plant species by reporting the proportion of fungal OTUs simultaneously present in different plants species and in different plant taxonomic groups. We split plant species into 7 main plant taxonomic groups: the bryophytes, the lycopods, the ferns, the monocots excluding *Orchidaceae*, the eudicots excluding *Ericaceae*, the *Orchidaceae*, and the *Ericaceae*.

We next investigated whether plant mycobiome compositions were evolutionarily conserved using a PERMANOVA that tested whether root samples belonging to the same plant taxonomic group were colonized by similar fungal OTUs. Given the phylogenetic breadth of the sampled communities, we evaluated the evolutionary conservatism of the plant mycobiome compositions by testing for discrete compositional shifts between the main plant taxonomic groups, *i.e.* using a PERMANOVA, instead of testing for continuous changes along evolutionary time [63]. In each sampled community, we also used hierarchical clustering to visually examine whether the mycobiome compositions tend to be clustered by plant taxonomic group. Given that the root mycobiota were dominated by 5 mycorrhizal fungal lineages (see Results), we also specifically replicated our PERMANOVA analyses on these lineages: the *Glomeromycotina*, *Mucoromycotina* (*Endogonales*), *Sebacinales*, *Helotiales*, and *Cantharellales*. The two former lineages were characterized using the 18S rRNA marker, whereas the three latter were characterized using the ITS2 marker.

### Reconstructing plant-fungus interaction networks

We used bipartite networks to study plant-fungus interactions. To reconstruct such networks, following [64], we considered that an interaction occurs between a plant and a fungus if the fungal OTU is represented by at least 1% of the reads of the root sample [64]. By converting read abundances into relative abundances, we thus corrected for the heterogeneous number of reads per sample and avoided counting spurious interactions occurring in samples with high coverage [65]. In addition, based on our estimates of cross contaminations (Supplementary Fig. 1), we considered that having less than 5 reads of an OTU within a sample was likely contamination. Using other cutoffs (*e.g.* 10 reads and 0.1%) did not qualitatively affect our results (not shown).

In each sampled community and each of the five main fungal lineages (*Glomeromycotina*, *Mucoromycotina*, *Sebacinales*, *Helotiales*, or *Cantharellales*), we considered three types of species-level plant-fungus networks: binary networks that do not consider interaction strengths and two types of weighted networks that differently account for interaction strengths. The first type - **binary networks** - indicates if an interaction between one plant species and one fungal OTU has been found in at least one root sample. The second type - **abundance networks** - is based on OTU read abundances within a root sample: for a given plant-OTU interaction, we reported its relative abundance as the number of reads belonging to this OTU per thousand of reads colonizing the corresponding plant species. However, as relative read abundances can be a bad proxy for the true fungal abundances colonizing the roots [64], we considered a third type of network, the **incidence networks**. For each plant-fungus interaction, the incidence network indicates the number of root samples in which the interaction has been found. To check that abundance and incidence networks gave similar quantifications of plant-fungus interaction strengths, we measured the relationship between them using linear models.

Using these reconstructed networks, we studied the fungal sharing in the different plant communities by (i) analyzing the structure of the plant-fungus networks and by (ii) evaluating the specialization among the plant-fungus interactions.

### Analyzing the structure of the plant-fungus networks

We analyzed and compared networks structures according to the fungal lineage and the sampled community. We first computed the connectance of each network, *i.e.* the percentage of realized interactions. A high connectance indicates that plant species are largely sharing fungi and a low connectance indicates infrequent fungal sharing. We also computed the checkerboard score (C-score), which measures the mean partner avoidance of pairs of species in a binary network. A high C-score indicates that some plant species tend to avoid sharing the same fungal OTUs. Next, we investigated whether plant-fungus networks were nested, *i.e.* whether specialist species tend to share partners with generalist ones. We computed nestedness using the NODF2 index for binary networks and the weighted NODF index for weighted networks (*nested* function in the bipartite R-package [66]). Lastly, we performed modularity analyses to evaluate whether some subsets of species, called modules, interact more with each other than with the rest of the species. We used Newman’s algorithm for binary networks and Beckett’s algorithm for weighted networks (*computeModules* function in the bipartite R-package). Modularity algorithms search for the most modular structure in the network and output the modularity value (M) and the ratio of interactions within modules (Q).

The significance of these structural properties was evaluated using two null models [67]. The first null model, generated using the quasiswap algorithm (*permatswap* function, R-package vegan [60]) keeps a constant connectance and constant marginal sums, *i.e.* the total number of interactions per plant species or fungal OTUs. Thus, the **quasiswap null model** investigates whether the structural properties of the network are conserved when plant-fungus interactions are randomly attributed based on the total availabilities of each interactor, with the additional constraint of keeping a similar connectance. The second null model shuffles the sample names and therefore randomly attributes the fungi associated with each root sample to a plant species. Thus, the **shuffle-sample null model** tests whether the emerging patterns in a species-level network come from the plant species properties and not from the sample properties [64].

By computing each index (*e.g.* nestedness, modularity…) for each null model and comparing their values to the ones of the original network, we get p-values indicating whether the original network has significant structural properties [68]. For instance, for the nestedness, if at most 2.5% of the null models have a higher or equal (resp. lower or equal) NODF value than the original one, the network is significantly nested (resp. anti-nested). We computed 10,000 null models using either the quasiswap or the shuffle-sample algorithms from our different original networks (binary, abundance, and incidence networks), which were used to evaluate the significance of the NODF values, checkerboard scores, and modularity values. In addition, we used the shuffle-sample null models to evaluate the significance of the connectance.

### Evaluating the degree of specialization among plant-fungus interactions

We evaluated the specialization of each plant species toward its fungal partners in each plant-fungus network. We only used the abundance network as a proxy for weighted interactions, as the incidence networks contained generally too little weighted information. We first computed the normalized degree of each plant species, as its number of fungal partners divided by the total number of available partners (*ND* function in the bipartite R-package). It indicates whether a species tends to be specialist (degree close to 0) or generalist (degree close to 1). Second, we computed *d’* which measures the specificity of a plant species toward its fungal partners (*dfun* function in the bipartite R-package [69]): a *d’* value close to 0 indicates that the plant species interacts with the most abundant fungal partners available with little specificity, whereas a *d’* value close to 1 indicates that the plant species specifically interacts with its fungal partners irrespectively of the abundance of the other fungi. From the *d’* values, we computed *H2’*, which is a network-level measure of specialization [69]. Low *H2’* indicates low specialization and therefore frequent fungal sharing among different plant species (and *vice versa*).

For each network, the significance of the indices of specialization was evaluated by generating 10,000 null models using the Patefield algorithm (*r2table* function, [69–71]). This null model algorithm keeps constant marginal sums but allows the connectance to vary. It thus tests whether the observed patterns of specialization are similar when interactions are only constrained by species abundances; we referred to them as the **marginal null models**. We also used the shuffle-sample null models (see the previous section).

To further compare the different plant-fungus networks in terms of specialization, we performed motif analyses, where a motif corresponds to a particular subnetwork architecture, a “building block”, between a given number of species [72]. For each plant-fungus network, we computed the frequencies of the motifs containing between 2 and 5 species using the *mcount* function from the BMOTIF R-package [72] and compared the motif frequencies in the different networks using PCoA. Finally, we specifically tested whether strong specialization in lycopod-fungus interactions exists in each sampled community (see Background): we compared their motif frequencies with those of the surrounding plant species to assess if lycopod species tend to be more associated with lycopod-specific fungi.

## Results

### The identification of the root-associated fungi shows a diversity of interactions

We collected a total of 233 root samples belonging to 30 plant species. We identified

29 of them at the species level (Fig. 2) and one species of *Ipomoea* remains taxonomically unidentified in Plaines-des-Palmistes (Fig. 2). We managed to collect 3 samples per plot per community for only 38% of the plant species because of the patchy distribution of some species. Yet, we collected at least 6 samples per community for 57% of the plant species, and only 11% of the plant species were represented by less than 3 samples.

**Figure 2:**
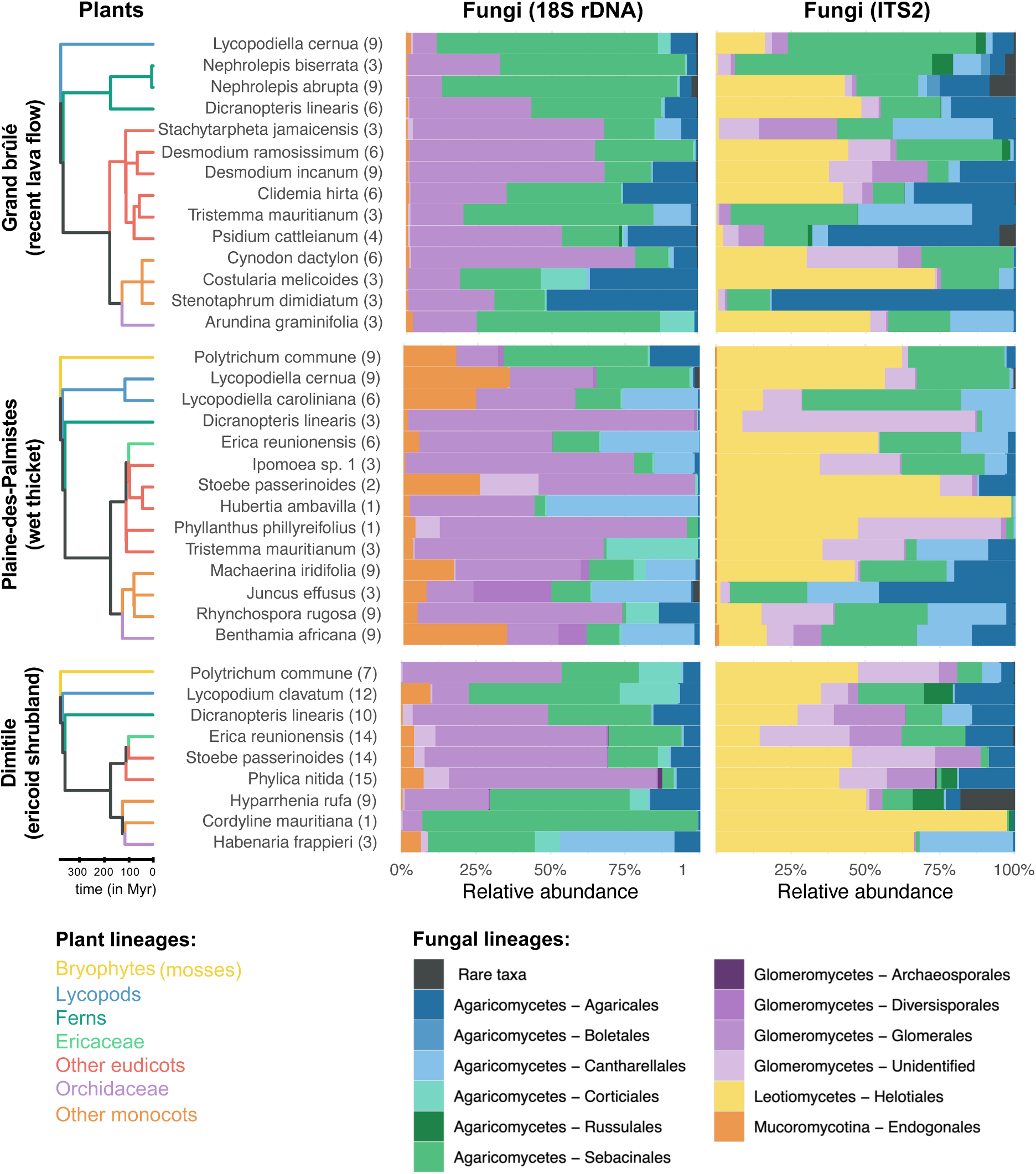
The composition of the root mycobiome varies according to the plant species and the habitats. For each plant species, the different root samples were merged and the relative abundances of the root fungi are indicated according to the 18S rRNA (left) or ITS2 (right) markers. We only retained the fungal lineages that may form mycorrhizal interactions with at least one plant species. Plant species are separated according to the sampled community (Grand brûlé, Plaine-des-Palmistes, or Dimitile) covering contrasted habitats. For each species, the number of individual root systems sampled is indicated in brackets. One species of *Ipomoea* remains taxonomically unidentified at the species level in Plaines-des-Palmistes. In each sampled community, a phylogenetic tree of the plants is represented on the left, with branch colors indicating the main plant taxonomic groups we considered in our study. The bar plots represent in colors the class and the order of each fungus. Rare taxa, representing less than 0.5% of the data, are plotted in dark grey.

We successfully amplified fungi in a total of 231 root samples, *i.e.* 99% of the sampled roots. We obtained a total of 25,250,698 and 26,914,809 reads for the ITS2 and 18S rRNA regions respectively, with an average coverage of 60,537 fungal reads per sample (+/- 29,044) for ITS2 and 19,414 fungal reads per sample (+/- 14,974) for 18S (Supplementary Table 2). We clustered these reads into 5,236 Swarm OTUs for ITS2 and 4,371 Swarms OTUs for 18S. When filtering the fungal OTUs assigned to putative mycorrhizal lineages, we obtained 622 OTUs for ITS2 and 1,177 OTUs for 18S, with a coverage larger than 1,000 reads in most of the root samples (Supplementary Fig. 2).

The two markers characterize different components of the root mycobiota (Fig. 2; Supplementary Fig. 3). Indeed, the 18S rRNA marker successfully detects colonization by *Glomeromycotina* and *Mucoromycotina* (*Endogonales*) fungi but fails at precisely characterizing *Basidiomycota*, which are at best identified at the order level, and even at detecting any *Helotiales* (*Ascomycota*). Conversely, the ITS2 marker fails at detecting *Mucoromycotina* fungi, but successfully amplifies and identifies *Basidiomycota* and *Ascomycota*, including the abundant *Sebacinales*, *Helotiales*, *Cantharellales*, and *Agaricales* orders. Altogether the five most abundant fungal lineages that may form mycorrhizal interactions, *i.e.* the *Glomeromycotina*, *Mucoromycotina*, *Sebacinales*, *Helotiales*, and *Cantharellales*, represent > 78% of the root mycobiota reads (Fig. 2).

The colonization frequency of *Glomeromycotina* is generally very consistent across root samples of the same plant species (Supplementary Table 1). To a lesser extent, the colonization frequencies of *Sebacinales* and *Helotiales* are also quite regular for some plant species (*e.g. Lycopodiella cernua*), but appear more facultative for others (*e.g. Dicranopteris linearis*; Supplementary Table 1). On the opposite, colonizations by *Mucoromycotina* and *Cantharellales* are recurrent in only a few plant species, such as *Lycopodiella caroliniana* or *Benthamia africana* respectively, but are very sporadic in others (Supplementary Table 1). In addition, we notice that both measures of interaction strength, using relative read abundance or interaction incidence, are significantly correlated (Supplementary Fig. 4), indicating that abundant interactions also tend to correspond to frequent ones.

Rarefaction curves representing OTU richness by plant species as a function of the number of root samples often do not reach a plateau, meaning that we may not document the entire diversity of fungi associated with plant species (Supplementary Fig. 5). Yet, sample-specific OTUs represent only ∼7% of the total root mycobiota reads.

### Root mycobiome compositions vary significantly across communities

The composition of the root mycobiomes varies significantly across communities, as revealed by both the hierarchical clustering and the PCoA of all the samples. The two methods show a clear clustering of the samples across the three sampled communities (Fig. 3; Supplementary Fig. 6a-b; PERMANOVA: p<0.05). For instance, *Mucoromycotina* are much more abundant in Plaine-des-Palmistes and rarer in other sampled communities (Fig. 2). This compositional variability is also found when comparing the mycobiota associated with the same plant species concurrently sampled in several communities (Supplementary Fig. 6c-d): When comparing the fungi present in the root samples of *Lycopodiella cernua* present in both Grand brûlé and Plaine-des-Palmistes, we find a significant shift in their composition, with enrichment in both *Mucoromycotina* and *Helotiales* in Plaine-des-Palmistes (Supplementary Fig. 7).

**Figure 3:**
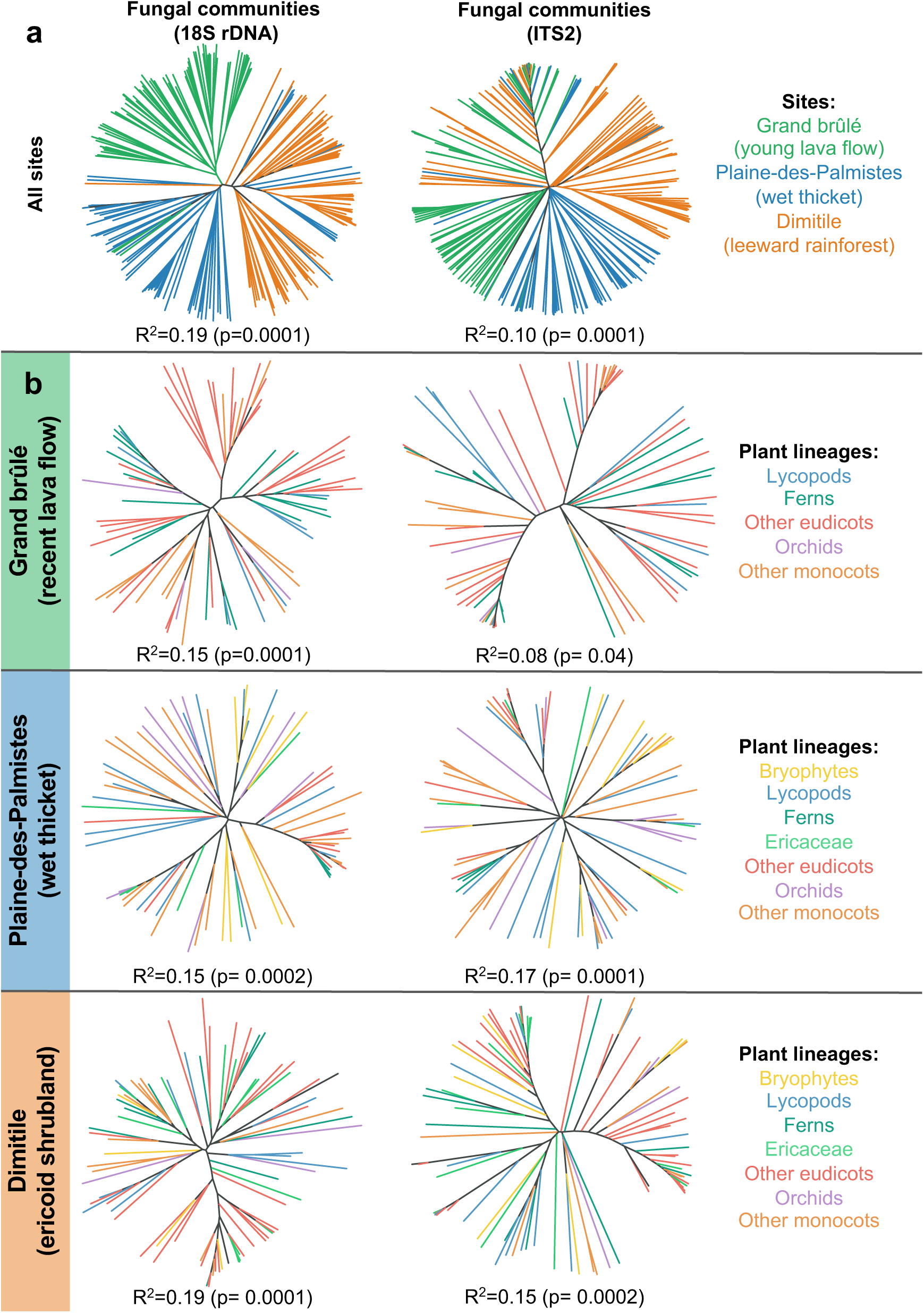
Both the habitat and the plant taxonomic group influence mycobiome composition despite frequent fungal sharing. Dendrogram representations of the different root mycobiota across all the sampled communities (**a**) or within each sampled community (**b**; Grand brûlé, Plaine-des-Palmistes, or Dimitile) based on the 18S rRNA (left) or ITS2 (right) markers. For each community, we only retained the fungal lineages that may form mycorrhizal interactions, computed the dissimilarity between pairs of samples (using Bray-Curtis distances), and reconstructed the dendrogram using neighbor-joining: two plant root samples that are close in the dendrogram tend to have similar fungal compositions. Branches are colored according to the sampled community (a) or to the plant taxonomic group (b). For each dendrogram, we also indicated the results of the PERMANOVA (R^2^ and p-value based on 10,000 permutations) testing the effect of the sampled community (top row) or the plant taxonomic groups (bottom rows) on the Bray-Curtis diversity between root samples.

To a lesser extent, we also find that samples from the same plots tend to cluster together (Supplementary Fig. 8a; PERMANOVA: p<0.05 in each community). However, this clustering per sampling plot is moderate (R^2^<0.10), and we thus merged the different plots to perform the following analyses at the community level.

### Weak but significant influence of the plant taxonomic groups on root mycobiome composition

Mycobiota compositions also tend to vary according to the different plant taxonomic groups (Fig. 2; Supplementary Fig. 3; Table 1). For instance, ferns are mainly colonized by *Glomeromycotina*, *Helotiales*, and *Sebacinales,* but not by *Mucoromycotina* (Table 1). In contrast, *Mucoromycotina* are regularly found across many plant species of other taxonomic groups, including the two early diverging lineages, bryophytes and lycopods (Table 1). Apart from *Mucoromycotina*, lycopods are recurrently colonized by *Glomeromycotina*, resulting in frequent *Mucoromycotina*-*Glomeromycotina* dual symbioses, but also by *Sebacinales*, *Helotiales*, and even by some *Cantharellales* (Supplementary Fig. 3). In angiosperms, most samples are mainly associated with *Glomeromycotina*, and to a lesser extent with *Helotiales*, *Sebacinales*, and *Cantharellales* (Fig. 2; Table 1). Besides their typical *Cantharellales* and *Sebacinales* partners, we also observe in *Orchidaceae* some unexpected colonization by *Mucoromycotina* and *Glomeromycotina*. Such changes in the plant-fungus associations according to the plant taxonomic groups are confirmed using PERMANOVA (0.08<R^2^<0.19, p<0.05; Fig. 3; Supplementary Table 3) and visually detectable when using hierarchical clustering (Fig. 3) or PCoA (Supplementary Fig. 8b), that both show a trend of a clustering per plant taxonomic group. We similarly observe that plant species from the same taxonomic group tend to interact with similar fungal OTUs when considering each fungal lineage separately (PERMANOVA: 0.09<R^2^<0.26; Supplementary Table 3).

**Table 1:** Plant taxonomic groups tend to be colonized by different fungal lineages of putative mycorrhizal fungi. This table recapitulates the fungal colonizations reported in the different plant taxonomic groups (Fig. 2, Supplementary Fig. 3). “++” indicates frequent and abundant colonizations, “+” indicates more sporadic and less abundant colonizations, and “-” indicates that the fungal colonization is never (or rarely) observed. Many of these colonizations are likely to be endophytic colonizations, rather than functional mycorrhizal associations.

| Plant taxonomic group | Fungal lineage |  |  |  |  |
| --- | --- | --- | --- | --- | --- |
|  | *Glomeromycotina* | *Mucoromycotina* | *Sebacinales* | *Helotiales* | *Cantharellales* |
| *Bryophytes (mosses)* | + | + | + | + | - |
| *Lycopods* | ++ | ++ | ++ | + | + |
| *Ferns* | ++ | - | ++ | + | + |
| *Monocots (excluding Orchidaceae)* | ++ | + | + | + | + |
| *Orchidaceae* | + | + | ++ | + | ++ |
| *Eudicots (excluding Ericaceae)* | ++ | + | + | ++ | + |
| *Ericaceae* | ++ | + | + | + | + |

Although plant taxonomic groups influence root mycobiome composition, we concomitantly detect frequent sharing of the fungal OTUs between plant species, even across different plant taxonomic groups. Indeed, in the three sampled communities, we find that ∼47% of the fungal OTUs are present in the roots of at least two different plant species, and ∼44% are even shared between different plant taxonomic groups (Supplementary Table 4). Such frequent fungal sharing across plant taxonomic groups tends to be even higher in some fungal lineages like the *Sebacinales*, the *Helotiales,* and especially the *Glomeromycotina* (Supplementary Table 4). The fact that many *Glomeromycotina* OTUs are shared across diverse plants explains the lower contribution of plant taxonomic groups in plant-*Glomeromycotina* interactions (PERMANOVA: R^2^<0.15) compared with other fungal lineages (Supplementary Table 3). Altogether, these shared fungal OTUs represent >90% of the total root mycobiota reads, meaning that fungal sharing is dominant in the sampled communities.

### The main fungal lineages harbor distinct network structures

The reconstructed species-level networks for the five main fungal lineages result in networks of different sizes and structures, reflecting important differences in terms of plant-fungus interactions (Fig. 4; Supplementary Fig. 9). Plant- *Glomeromycotina* networks have species-rich, well-connected, typical nested structures with a core of abundant generalists surrounded by rare specialists, whereas plant- *Mucoromycotina* and plant-*Cantharellales* networks appear to be species-poor, less connected, and much more modular, and plant-*Sebacinales* and plant-*Helotiales* networks have intermediate topologies (Fig. 4). When looking at the position of the different plant species in the networks, we notice that plants from the same taxonomic group mostly tend to be closer (Fig. 4), as previously indicated by the hierarchical clustering and PERMANOVA. However, this clustering is not exclusive: lycopods, ferns, and bryophytes appear to be generally well connected to angiosperms (monocots and eudicots) by shared fungi (Fig. 4; Supplementary Fig. 9). On the opposite, *Orchidaceae* and *Ericaceae* species often form different modules, mainly separated from the other species forming the interaction core (see for instance the plant-*Sebacinales* network in Grand brûlé or the plant-*Mucoromycotina* network in Plaine-des-Palmistes). These differences in terms of network structure are robust when excluding the rare plant species represented by less than 3 samples (Supplementary Fig. 10).

**Figure 4:**
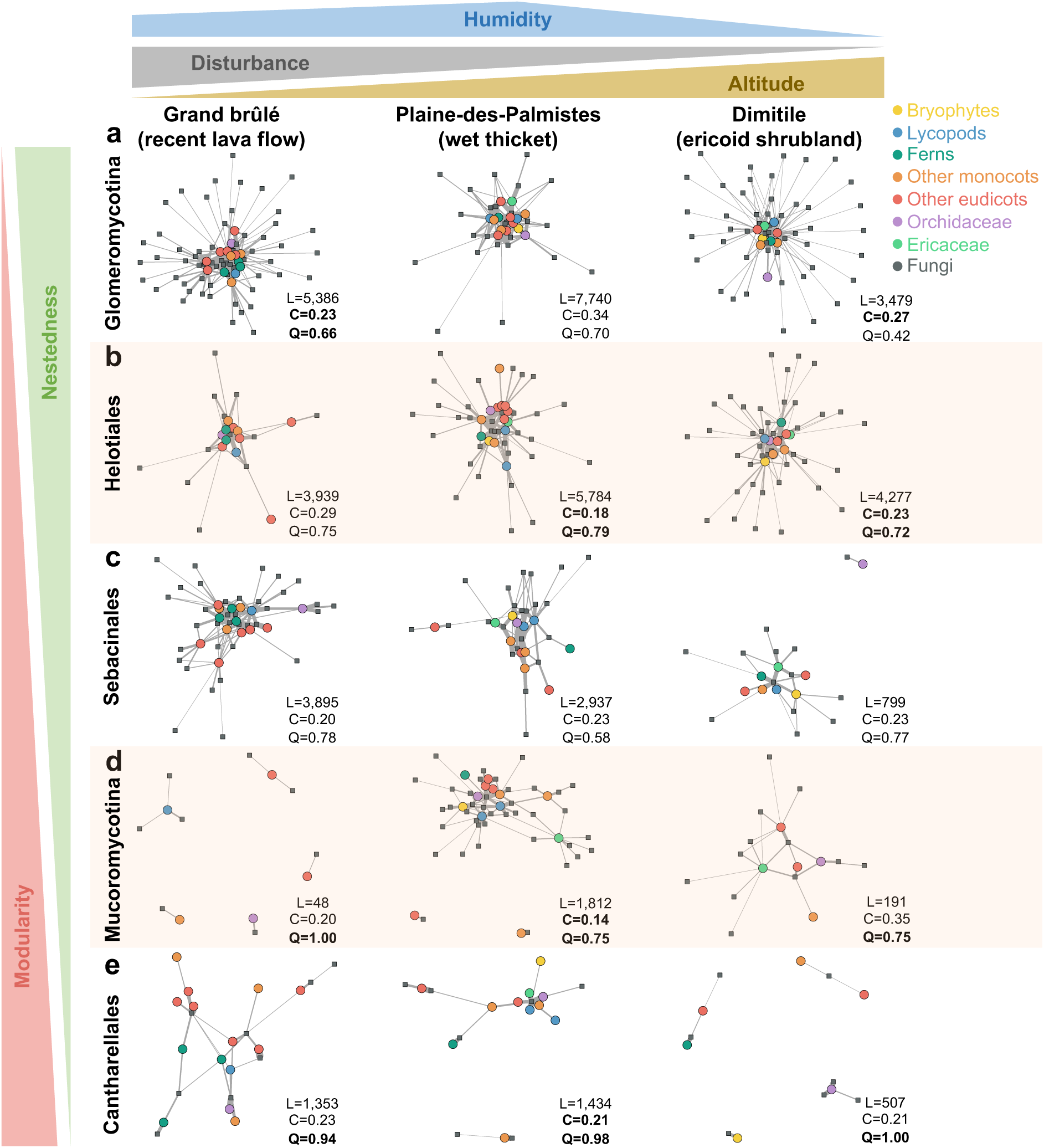
Plant-fungus network structures vary between fungal lineages, irrespectively of the environmental variations. Species-level network representation in each sampled community (Grand brûlé, Plaine-des-Palmistes, or Dimitile) for the different fungal groups: *Glomeromycotina* (a), *Helotiales* (b), *Sebacinales* (c), *Mucoromycotina* (d), or *Cantharellales* (e). Colored round nodes represent plant species (colors indicate the main plant taxonomic groups) and grey squared nodes correspond to fungal OTUs. Grey links represent plant-fungus interactions and their widths are proportional to interaction abundances. The position of the nodes reflects the similarity in species interactions using the Fruchterman-Reingold layout algorithm [99] from the *igraph* R-package. Fungal lineages (in rows) are ordered according to their network structures: networks that tend to be nested are at the top, whereas networks that tend to be modular are at the bottom (Supplementary Tables 6-8). The sampled communities are in columns and we indicated the environmental gradients on the top. For each network, the total read abundances (L), the connectance (C), and the ratio of interactions within modules (Q) are indicated. Q is computed from the most modular structure according to Beckett’s algorithm for abundance networks, Q close to 0 indicates that most interactions are between modules (*i.e.* low modularity), while Q close to 1 indicates that most interactions are within modules (*i.e.* high modularity). Significant connectance and modularity values, evaluated using shuffle-sample null models, are highlighted in bold. We did not report nestedness values as they are not meaningful when compared across networks of different sizes. Details about the fungal taxonomy can be seen in Supplementary Figure 18.

Our quantitative investigation of network structures reveals an important range of connectance values (*i.e.* the percentage of realized interactions) from 0.14 to 0.34, confirming that fungal sharing between plant species is important. In addition, plant-fungus networks tend to be less connected than the shuffle-sample null models (Fig. 4, Supplementary Table 5), indicating that plant-fungus interactions within plant species are more alike in samples than between plant species. In terms of nestedness, compared to the quasiswap null models, and when considering weighted interactions, large networks, like plant-*Glomeromycotina* and plant-*Helotiales* networks, are often significantly nested (Supplementary Table 6a). On the opposite, the smaller plant-*Sebacinales*, plant-*Cantharellales*, and plant-*Mucoromycotina* networks are often non-significantly nested. When considering shuffle-sample null models, all networks except plant-*Glomeromycotina* incidence networks appear non-significantly nested (Supplementary Table 6b). We also observe similar trends when comparing the C-score of the networks to null models (Supplementary Table 7): Nested/anti-checkerboard structures, reflecting strong asymmetrical specialization and important partner sharing, are only significant in large networks and when considering interaction strength. Finally, there is limited evidence for significant modular structures in the networks. Indeed, most of the networks are not significantly modular (Supplementary Table 8). Yet, for those that are significantly modular, in particular plant-*Mucoromycotina*, plant-*Cantharellales*, and plant-*Helotiales* networks, we observe Q values (the proportion of within-modules interactions) above 0.75 (Fig. 4), suggesting that these inferred modules explain more than 75% of the interactions. Many of these non-significances might arise from the relatively small sizes of the network, which reduce the statistical power of the null model comparisons.

### The main fungal lineages show distinct levels of specialization with plants

The *H2’* values, characterizing the average network-level specialization, are lower in plant-*Glomeromycotina* networks than in the other networks (Fig. 5a; Supplementary Fig. 11), indicating that plant-*Glomeromycotina* interactions tend to be less specialized than other plant-fungus interactions. Conversely, plant-*Mucoromycotina* and plant-*Cantharellales* interactions appear to be highly specialized (high *H2’* values), and plant-*Sebacinales* and plant-*Helotiales* interactions show intermediate levels of specialization (Fig. 5a). These patterns of specialization are robust when excluding the rare plant species represented by less than 3 samples (Supplementary Fig. 12). Looking at the specialization of the individual plant species, both normalized degree and *d’* indicate that most plant species are more specialized toward their fungi than expected when interactions are randomly distributed based on species abundances (marginal null models; Supplementary Fig. 13). However, these results do not hold with the shuffle-sample null models, indicating that many plant-fungus specializations are driven by sample effect rather than by species effect.

**Figure 5:**
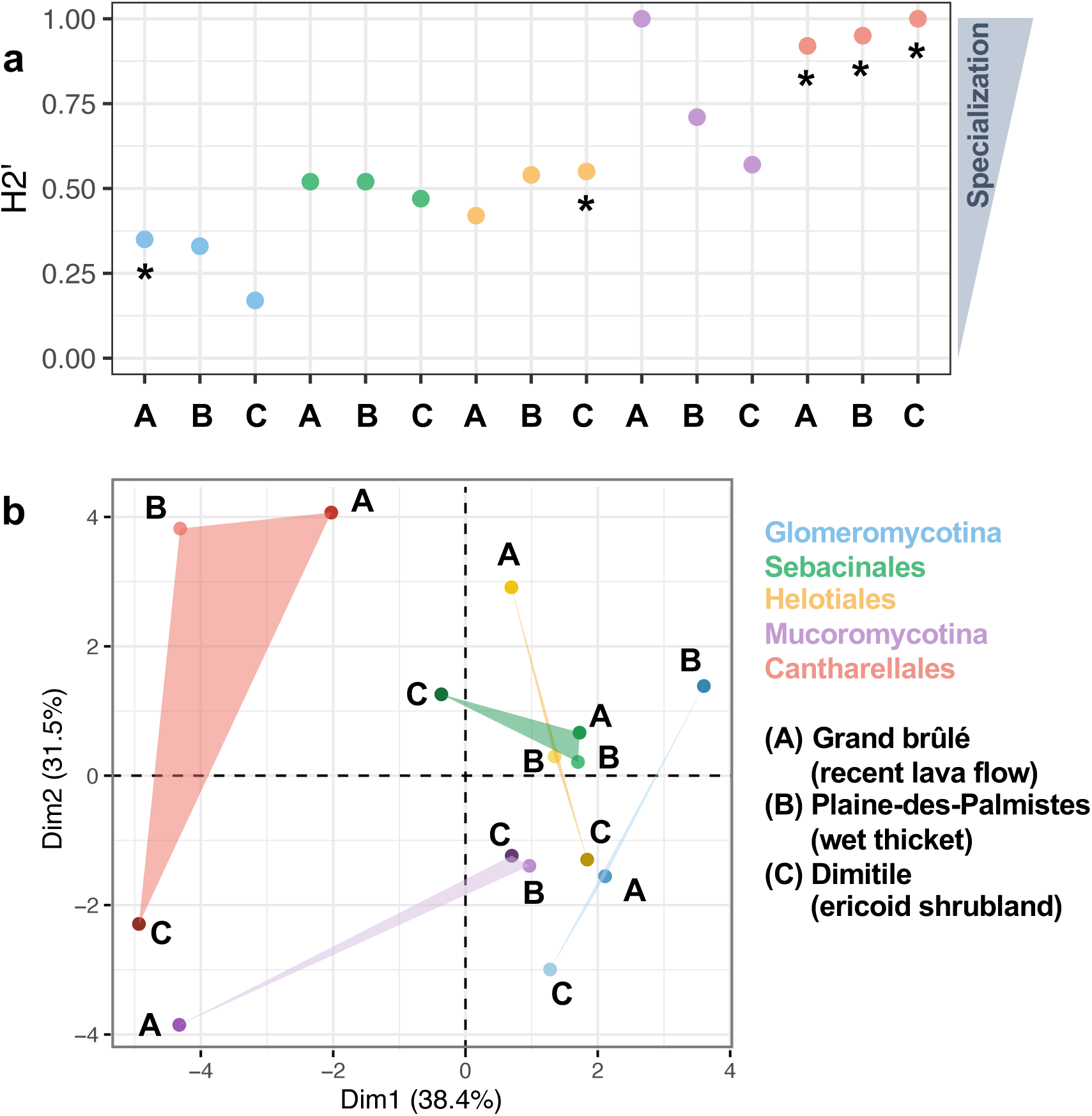
Fungal sharing in the plant-fungus networks varies across the different fungal lineages. **(a) Interaction specializations (H2’) are lower in plant-*Glomeromycotina* networks than in other plant-fungus networks.** For each plant-fungus network (with *Glomeromycotina*, *Mucoromycotina*, *Sebacinales*, *Helotiales*, or *Cantharellales*) in each sampled community (Grand brûlé (A), Plaine-des-Palmistes (B), or Dimitile (C)), a colored dot indicates the network-level interaction specialization (H2’). The significance of the H2’ values was evaluated using null models maintaining marginal sums or shuffle-sample null models: all the H2’ values were significant for the marginal sums null models, and asterisks indicate when the H2’ values are significant, based on the shuffle-sample null models (see Supplementary Figure 11 for details). **(b) Motif frequencies significantly differ between the plant-fungus networks.** Principal coordinate analyses (PCoA) of the bipartite motif frequencies (the “building blocks” of the network containing from 2 to 5 species) of each plant-fungus network (*Glomeromycotina*, *Mucoromycotina*, *Sebacinales*, *Helotiales*, or *Cantharellales*) in each sampled community (Grand brûlé (A), Plaine-des-Palmistes (B), or Dimitile (C)). The colored triangle areas represent the proximity within the sampled communities for the different groups of fungi.

When comparing the motif frequencies of the different networks using PCoA, *i.e.* the different frequencies of the “building blocks” of the networks, we find that plant-fungus network tends to cluster by fungal lineage (Fig. 5b). In particular, we observe that motifs constituted of plants and fungi that are densely interacting, indicating a lot of sharing, as motifs 11 or 12, tend to be more frequent in plant-*Glomeromycotina* networks (Supplementary Fig. 14).

To ensure that the low level of specialization observed in plant-*Glomeromycotina* interactions does not arise from the use of the 18S rRNA marker, we reproduced the analyses using the more variable ITS2 marker and observe similar patterns (Supplementary Fig. 15). Similarly, replicating the plant-*Sebacinales* network analyses with the 18S rRNA marker confirms that plant-*Sebacinales* interactions are on average more specialized than the plant-*Glomeromycotina* ones (Supplementary Fig. 16). Thus, the observed trends are not sensitive to the marker used.

We do not find a strong specialization in lycopod-*Glomeromycotina* interactions in the sampled community (Supplementary Fig. 17). Indeed, compared with the surrounding plant species, *Lycopodiella cernua* in Grand brûlé, *Lycopodiella caroliniana* in Plaine-des-Palmistes, or *Lycopodium clavatum* in Dimitile tend to be relatively more associated with *Glomeromycotina* partners that are well connected to other plant species in the networks, as illustrated by the high frequencies of motifs 19, 26, or 32 (Supplementary Fig. 17). In contrast, concerning associations with non-*Glomeromycotina* fungi, lycopod species tend to interact more frequently with specific fungi that are not connected with any other plant species in the network, like the motifs 17, 20, 23, or 33, which correspond to lycopod-specific fungi (Supplementary Fig. 17).

## Discussion

In this study, we thoroughly characterize the plant-associated fungi that may form mycorrhizal interactions within three contrasted local communities including distantly related plant species. We find that these plant communities across La Réunion island are mainly colonized by five lineages of putative mycorrhizal fungi. Against our expectations, we report only a weak evolutionary conservatism in these plant-fungus interactions. Instead, we notice frequent fungal sharing between phylogenetically distant plant lineages. When looking in detail at the different fungal lineages, we observe striking differences in terms of specialization and network structure, *i.e.* in the way the fungi are shared by plant species. Interestingly, while the composition of the root mycobiomes varies according to the sampled communities, the plant-fungus network structures do not seem to be impacted by such environmental variations.

### Characterizing the composition of the root mycobiomes

We notice that the ITS2 region, amplified with the ITS86F and ITS4 primers, and the 18S rRNA gene, amplified with the AMAD-f and AMDG-r primers, characterize different aspects of the root mycobiota. While the 18S rDNA offers better visualization of all the fungal lineages, the more resolutive ITS2 region enables better identification of *Dikarya*. In particular, we notice that *Mucoromycotina* (*Endogonales*) are almost always missing when using the common ITS86F and ITS4 primer pair, whereas they appear to be frequent endophytes and major mycorrhizal symbionts [13, 18]. These results therefore encourage systematically characterizing the composition of the root mycobiome by targeting both sets of regions. Yet, additional fungal lineages may still not be represented in our dataset, like the *Tulasnellaceae,* which can form mycorrhizal associations with orchids but often fail to amplify with generalist primers [50,73,74]. This suggests that even two generalist and complementary primer sets might still not be sufficient to retrieve the whole endophytic fungal community.

Compared with other studies, which characterized plant-fungus interaction networks in communities dominated by angiosperms in temperate habitats [35–38], we analyze plant-fungus interactions in tropical communities comprising diverse plant groups including “early-diverging” lineages. Yet, we also detect the same groups of mycorrhizal fungi that are typically found in temperate habitats. *Glomeromycotina* abundantly colonize most plant lineages, including bryophytes, lycopods, ferns, and many angiosperms [5, 7]. Similarly, we observe that *Mucoromycotina* are frequently associated with a range of plants [13], with the exception of ferns, as previously suggested by [18]. We also confirm that *Sebacinales* are major root endophytes [29, 75] and frequently colonize lycopods [19]. Similarly, *Helotiales* and *Cantharellales* are also retrieved as root endophytes of many plant species. More surprisingly, we detect an abundant *Mucoromycotina* colonization in *Orchidaceae*. Such an association has been previously detected in epiphytic *Orchidaceae* [76] and further works using the 18S rRNA marker as well as using microscopy methods should be pursued to confirm the permanent character of this association.

Most single plants are usually colonized by several fungi, among which some may be mycorrhizal. For instance, we notice that dual colonization by *Mucoromycotina* and *Glomeromycotina* are particularly frequent, especially in lycopods. Experiments have demonstrated that such dual symbioses can both be functional and have complementary nutritional roles [9]. However, here, our molecular detection of a fungus in a plant root can either correspond to a functioning mycorrhiza, an opportunistic or beneficial endophyte, or even (but less likely given the high abundances) a sporulating fungus [7, 42]. Testing which colonization corresponds to a mycorrhiza, and whether multiple fungal colonizations have all a functional role and are complementary will be a future step requiring targeted experimental manipulations.

We observe an important variability in the mycobiome composition between samples from the same plant species in a given community. Consequently, some interactions between a plant species and a fungal OTU are only supported by one or two root samples. This therefore explains why many apparent plant-fungus specializations at the species level are not significant when using the shuffle-sample null models. In addition, the rarefaction plots show that even >10 sampled roots per plant species are often not sufficient to get the whole diversity of the associated fungi. Yet, sample-specific OTUs represent only ∼7% of the root mycobiota, suggesting that only some rare fungal taxa sporadically colonizing some plant species are missing. In addition, replicating our analyses while excluding under-sampled plant species confirmed the robustness of our results in terms of network structure and specialization. This indicates that the incomplete characterization of the root mycobiota composition is unlikely to bias our conclusions.

### Effect of the community and the plant taxonomic group on root mycobiome composition

We find a strong effect of the sampled communities on the composition of the root mycobiomes, suggesting that the different environmental conditions of our three sampled communities in terms of humidity, disturbance, and elevation may influence the fungal distributions and the root mycobiota. Extensively sampling more communities covering the environmental gradients would allow unravelling the different effects of the environmental variables and excluding the potential effect of dispersal limitation or random variations expected under models of stochastic colonizations. In addition, we observe that mycobiota composition varies significantly across plots within a given community. This suggests that there are important variations at small spatial scales in the assembly of the root mycobiome [77, 78].

Among the strong community effects, we report that *Mucoromycotina* fungi are relatively abundant in the wet thickets (Plaine-des-Palmistes) and mostly absent from other sampled communities. Consequently, the root mycobiome composition of *Lycopodiella cernua* presents a clear shift according to its environment: while this plant mainly associates with *Glomeromycotina* and *Sebacinales* in Grand brûlé, *Mucoromycotina* represent up to 90% of its reads in Plaine-des-Palmistes. This raises the question of whether the occurrence of *Mucoromycotina* is favored in wet conditions. *Mucoromycotina* have been observed colonizing *Horneophyton lignieri*, a 407 million-year-old plant [8] that generally preferred sandy and organic-rich substrates, although it has been reported that this plant could develop in wet conditions [79].

The significances of the PERMANOVA indicate that, as expected, plant-fungus interactions are evolutionarily conserved in the different plant lineages. Yet, the relatively low R^2^ values of these PERMANOVA suggest that the different plant taxonomic groups only explain a small part of the compositional variations of the plant root mycobiomes. This is likely due to two non-exclusive explanations. First, although two mycobiota from the same plant species are on average more alike than two mycobiota from different plant species, we observe an important variability in the mycobiome composition between samples. Second, we detect frequent sharing of the fungal OTUs between co-occurring plant species, including across different taxonomic groups. Plant taxonomy has therefore only a limited influence of root mycobiome composition in diverse tropical communities.

### Levels of fungal sharing vary according to the fungal lineages

In local communities, against our initial expectations, we observe frequent fungal sharing between co-occurring plant species, including between phylogenetically distant plants. Therefore, we find little evidence for the statement “ancient plants with ancient fungi”, resulting from the recurrent observations of interactions between “early-diverging” liverworts and “early-diverging” non-*Glomeraceae* lineages [15]. All our analyses indicate that fungal sharing is particularly important for *Glomeromycotina*, which present very little specialization, as already often reported in local communities of angiosperms [3, 35]. Conversely, the other fungal lineages, especially *Mucoromycotina* and *Cantharellales*, are more specialized and often more sporadic in their interactions with plants. *Sebacinales* and *Helotiales* present intermediate levels of specialization, confirming that they are widespread root endophytes (*e.g.* [29, 75] for *Sebacinales* and [40] for *Helotiales*).

Strong differences in specialization might reflect the different evolutionary origins of these plant-associated fungi. Indeed, *Glomeromycotina* are thought to be ancestral plant symbionts that obligately associate with them [5,6,8,80]. Although some plants have lost their dependence on *Glomeromycotina* through time [81], they often retain the ability to occasionally host sparse *Glomeromycotina* fungi [7, 82], which could explain why *Glomeromycotina* tend to colonize many plant species with very low specificity. Conversely, *Sebacinales*, *Helotiales*, and *Cantharellales* are younger clades that have more recently acquired their ability to interact with plants [5]. In addition, many of these lineages are still saprotrophs [12,83,84]: these fungal lineages are therefore less dependent on plants than obligate biotrophic ones [85]. Thus, plant colonization can be more facultative for them and often requires a minimal plant-fungus specificity to be established [3], which could explain the higher specialization we have observed for these lineages. Next works should focus on the genomic determinants of such plant-fungus interactions [83, 86].

We also find that plant-*Mucoromycotina* interactions are quite specialized, facultative, and mainly limited to wet environments. Yet, *Mucoromycotina* are increasingly recognized as ancestral plant symbionts [5,8,13,87]: These associations might have played a primordial role in land colonization, without experiencing an important diversification of their ecological niches later.

Against what was suggested by a recent metanalysis of plant-*Glomeromycotina* interactions on a global scale [41], we observe that lycopod sporophytes are well connected to other plant species by fungal sharing. Nevertheless, compared to other plant species, we also notice the propensity of lycopod species to interact with lycopod-specific fungal OTUs, especially from *Mucoromycotina* and *Sebacinales*. Such lycopod-specific fungal OTUs have been hypothesized to sustain parental nurture between the autotrophic lycopod sporophytes and their achlorophyllous and subterranean gametophytes that rely on mycorrhizal fungi for their nutrition [88–91]. Future works should particularly focus on these achlorophyllous gametophytes to investigate what fungi provide them nutrients and to assess whether or not parental nurture occurs thanks to lycopod-specific fungi [91]. During our study, we have only found one gametophyte of *Lycopodiella cernua* close to Plaine-des-Palmistes, and this gametophyte was abundantly and specifically colonized by a single *Mucoromycotina* OTU also present in *Lycopodiella cernua* sporophytes.

### Structural distinctiveness in the plant-fungus networks

The different levels of specialization of the main fungal lineages resulted in different network structures, as illustrated by both motif and structural analyses. In particular, plant-*Glomeromycotina* networks tend to exhibit significant nestedness, confirming a pattern frequently observed in local communities of angiosperms [35,37,92]. Other plant-fungus networks, especially those comprising *Mucoromycotina* or *Cantharellales*, tend to have less connected, un-nested, and even modular structures, reflecting the higher specificity of these plant-fungus interactions. Our results thus support the idea that non-*Glomeromycotina* plant-fungus networks tend to have un-nested structures, as previously observed in local communities of angiosperms [43, 93] or in the liverwort-*Mucoromycotina* network on a global scale [94]. By separately looking at the main fungal lineages, our approach thus better captured the peculiar patterns of interactions with plants of these different fungal lineages, which can be missed when merging and studying all fungi in the same framework [42].

Some of the structural differences in the plant-fungus networks may be driven by differences in the fungal traits that affect the colonization of plant roots and the subsequent interaction. For instance, the *Cantharellales* present a large panel of ecological strategies [95] and plant-*Cantharellales* interactions can range from mutualism to antagonism (*e.g.* the pathogen *Rhizoctonia solani*). Such variability may generate modularity in the network [96] and might explain why plant-*Cantharellales* tend to form the most modular networks. In addition, we notice that lineages that have retained saprophytic abilities, like *Cantharellales* and *Mucoromycotina*, form more modular networks than obligate biotrophic endophytes, like the *Glomeromycotina* and some *Sebacinales*, suggesting that the resulting network structures are partially determined by the fungal niches. Improving the characterization of the individual fungal ecological niches [97] and accounting for fungal traits [98] will allow us to investigate the patterns of interactions at finer taxonomic and functional scales.

Despite the contrasted environmental conditions of our three sampled communities, we have observed consistent network structures for each fungal lineage across the different sampled communities. This suggests that even if the environmental conditions impact the relative abundance of the main fungi, they do not strongly influence the network structure. In other words, how fungi interact with plants and are shared between them likely results from the intrinsic properties of each fungal lineage, and not from environmental conditions. Our result contrasts with a recent metanalysis of plant-fungus interactions reporting that the mean annual rainfall has more influence on the level of nestedness of plant-fungus interaction networks than the fungal lineage involved [43]. Yet, while we only compare species-level networks at the level of the whole plant community, Põlme *et al.* [43] merged in their analyses very heterogeneous types of networks. Indeed, many networks they included were either individual-level networks or species-level networks restricted to a specific plant clade (*e.g.* only the *Orchidaceae*-fungus networks). Future works should then investigate why species-level *versus* individual-level networks may differentially respond to environmental variations.

## Conclusion

By characterizing plant-fungus interactions in tropical communities, we reveal that fungal sharing is widespread in local communities, even among distantly related plants. Our study demonstrates the distinctiveness in terms of specialization and network structure of the main fungal lineages detected. We suggest that this distinction is likely underpinned by the peculiar functioning and ecologies of these plant-fungus symbioses. This finding opens the door for future works aiming to characterize the functions of the root mycobiota in plant-plant interactions, which can range from nutrition to protection. Altogether, our findings highlight the importance of systematically considering the whole network of interactions (the “interactome”) rather than isolated macroorganisms and their associated microbes (the “holobiome”).

### Declarations

#### Availability of data and material

Scripts used for generating the OTU tables are available at https://github.com/BPerezLamarque/Scripts. OTU tables, associated metadata, and R script for replicating the network analyses are publicly accessible through the Open Science Framework (osf) portal: osf.io/bskxc.

#### Competing interests

The authors declare no conflict of interest.

#### Funding

This work was supported by the Agence Nationale de la Recherche (ANR-19-CE02-0002). BPL was supported by a doctoral fellowship from the École Normale Supérieure de Paris attributed to BPL and the École Doctorale FIRE – Programme Bettencourt. HM acknowledges support from the European Research Council (grant CoG-PANDA). C.S-D acknowledges the Fondation ARS Cuttoli-Paul Appell/Fondation de France for supporting her work on fossil fungi (grant 00103178).

#### Authors’ contributions

All authors designed the study. BPL, HM, and FM performed the fieldwork; BPL and FM did the molecular works; BPL performed the analyses and wrote a first draft of the manuscript and all authors contributed substantially to the writing.

## Acknowledgments

The authors thank Benoît Lequette, Jean-Marie Pausé, Audrey Valery, and Claudine Ah-Peng for support during the field work. The *Parc National de La Réunion* authorized sampling in La Réunion (N° DIR-I-2019-149). Logistic support was provided by the field station of Marelongue, funded by the P.O.E., Reunion National Park and OSU Reunion. The authors thank the Service de Systématique Moléculaire (UMS 2700) for technical support, as well as Chantal Griveau, Amélia Bourceret, and Céline Bonillo. They also thank Mélanie Roy and Philippe Vandenkoornhuyse for helpful discussions, as well as the Editor and two anonymous reviewers for constructive comments.

## Notes

### Competing Interest Statement

The authors have declared no competing interest.

