## Supplementary Fig for "Structure and specialization of mycorrhizal networks in phylogenetically diverse tropical communities"

**Supplementary Methods (3)**
**Supplementary Tables (1-8)**
**Supplementary Figures (1-18)**
**Supplementary References**

### Supplementary Methods:

#### Supplementary Methods 1: Molecular work

We amplified two nuclear regions of the rDNA operon, the 18S rRNA gene and the ITS2 region, using two sets of tagged primers: AMADf-AMDGr pair (Berruti *et al.*, 2017) and ITS86F-ITS4 pair (White *et al.*, 1990; Turenne *et al.*, 1999), that respectively amplify a fragment of 380 and 280 base pairs on average. PCRs were performed using the AmpliTaq Gold™ 360 kit (Thermo Fisher Scientific) following the manufacturer's instructions and ran for a total of 35 cycles at an annealing temperature of 50°C for the 18S rRNA marker and 56.5°C for the ITS2 one. To ensure better characterization of the mycobiota, the PCR step was replicated at least 2 times. Each sample was characterized by a unique combination of tagged primers (following Taberlet *et al.*, 2018; Petrolli *et al.*, 2021), and some combinations of primer pairs were left empty to evaluate the amount of tagged index hopping between samples (Zinger *et al.*, 2019). In addition, negative PCR controls were also conducted (i) to identify the possible external contaminants and (ii) to evaluate the baseline cross-contamination between samples during the library preparation.

For both libraries (constituted of either the 18S rRNA or the ITS2 amplicons), PCR products were purified using magnetic bead-based clean-up (NucleoMag™ NGS Clean-up and Size Select, Macherey-Nagel), quantified using Qubit (Qubit dsDNA High Sensitivity Assay Kit, Invitrogen)), and (up to) 3 ng of DNA amplicons per sample (including controls) were sequenced using Illumina 2x250 bp MiSeq technology (v3 chemistry, Fasteris, Geneva).

**Supplementary Methods 2: Bioinformatics**

The 18S rRNA reads and the ITS2 reads were separately processed using a pipeline based on VSEARCH (Rognes *et al.*, 2016) available on GitHub (https://github.com/BPerezLamarque/Scripts/). Pair reads were assembled, quality checked (removing reads that had on average more than 2 errors), demultiplexed (using cutadapt (Martin, 2011)), and clustered into operational taxonomic units (OTUs) using two different methods. We first used Swarm v2 (Mahé *et al.*, 2015) with the fastidious option, a clustering approach that does not rely on a global threshold of similarity but instead uses local thresholds and amplicon abundances. Secondly, we performed a classical 97% OTU clustering using VSEARCH. We removed the chimera in both sets of OTUs using *uchime_denovo* in VSEARCH and assigned taxonomy to each OTU using *usearch_global* (BLAST algorithm). For the latter step, we used the Silva database r138 (Quast *et al.*, 2013) for the taxonomic assignations of the 18S rRNA OTUs and the UNITE database v8.2 (Nilsson *et al.*, 2019) for those of the ITS2 OTUs. Because the Silva database contains only a few Agaricomycetes sequences, we also assigned taxonomy at the order level to these Agaricomycetes 18S rRNA OTUs using the Fungal 18S RefSeq Targeted Loci Project (NCBI BioProject - PRJNA39195). We finally built an OTU table for each marker and removed OTUs present in less than 10 reads. *Glomeromycotina* characterized using the 18S rRNA marker are often assigned to a *virtual taxa* (VT) based on the MaarjAM database (Öpik *et al.*, 2010), however, the AMADf-AMDGr primer pair used in this study led to 18S rRNA amplicons that only partially overlap with *virtual taxa* and prevented us to do the assignations (Davison *et al.*, 2015).

Next, the decontam pipeline in R (R Core Team, 2020) was applied to filter out the contaminants of our OTU tables, using the read abundances within the negative controls as well as the abundance profile of each OTU (Davis *et al.*, 2018).

Finally, we reconstructed the phylogenetic tree of both fungi and plants. For the tree of the fungal OTUs, for each main fungal lineage, we aligned the OTU sequences using MAFFT (Katoh & Standley, 2013), trimmed them with trimAl (Capella-Gutierrez *et al.*, 2009), selected the best substitution model using ModelFinder (Kalyaanamoorthy *et al.*, 2017), and reconstrued the maximum-likelihood tree using IQ-TREE (Nguyen *et al.*, 2015) with 1,000 SH-aLRT and ultrafast bootstraps. For each main fungal lineage, a set of 3 OTUs were chosen as an outgroup (from a closely related fungal lineage) in order to root the phylogenetic trees.

For the plants, the plant mega-phylogeny from (Zanne *et al.*, 2014) was pruned using Phylomatic (http://phylodiversity.net/phylomatic/) to obtain the phylogenetic tree of the plant species sampled in our three sampled communities. Plant taxa that were not identified at the species levels were added as polytomies at the origin of the clade.

Given that both Swarm and 97% OTU clustering gave qualitatively similar results, only the results obtained with Swarm are reported here.

**Supplementary Methods 3: Evaluating cross-contaminations**

First, thanks to our range of negative controls, we evaluated the amount of index hopping (using tagged primers left empty in the entire design) and cross-contaminations (using PCR negative controls). For most of the empty combinations of tagged primers, index hopping was negligible and below 5 reads (Supplementary Fig. 1a). For most of the PCR negative controls, contaminations were below 100 reads, but some samples had abundant contaminant concentrations (Supplementary Fig. 1 b). The contaminants mainly corresponded to human-skin *Malassezia* OTUs or other saprotrophs. All these negative OTUs were removed using the decontam pipelines (Davis *et al.*, 2018). Less than 1% of the reads found in the negative controls were fungi that may form mycorrhizal interactions, suggesting that cross-contamination was low. Thus, we considered that discarding the OTUs present in less than 5 reads in a sample represented a good balance to avoid the consideration of spurious interactions likely coming from cross-contaminations or index hopping.

Second, we verified that samples that were physically close in a plate (the organization of the plates was kept identical from the DNA extraction to the final amplicon pooling) did not tend to present a more similar composition than expected by chance because of cross-contaminations. To do so, we evaluated the similarities of the composition using Bray-Curtis dissimilarities computed using the *vegdist* function of the vegan R-package (Oksanen *et al.*, 2016) and tested whether they were correlated with the Euclidian physical distances in each PCR plate using Mantel tests. We found no significant correlations (Mantel test: p-value>0.05) suggesting that cross-contaminations were overall negligible and did not significantly influence our results.

#### Supplementary tables:

**Supplementary Table 1: The recurrence of plant root colonization varies according to the fungal lineages**

The total number of individual plant root systems sampled per plant species per sampled community (Grand brûlé, Plaine-des-Palmistes, or Dimitile) is indicated in the second column. Then among these samples, the following columns indicate the number of samples per plant species containing each type of symbiont (*Glomeromycotina, Mucoromycotina, Sebacinales, Helotiales*, or *Cantharellales*), based on the Swarm OTUs. Non-native plant species are marked with an asterisk (*). Plant species with an uncertain taxonomic assignation at the species level and that may be non-native are indicated with a double asterisk (**).

| Plant species | Grand brûlé | | | | | |
|---|---|---|---|---|---|---|
| | Number of samples | *Glomeromycotina* | *Mucoromycotina* | *Sebacinales* | *Helotiales* | *Cantharellales* |
| *Arundina graminifolia* * | 3 | 2 | 1 | 3 | 3 | 2 |
| *Clidemia hirta* * | 6 | 6 | 1 | 5 | 3 | 2 |
| *Cynodon dactylon* | 6 | 6 | 1 | 5 | 6 | 1 |
| *Desmodium incanum* * | 9 | 9 | 1 | 6 | 7 | 4 |
| *Dicranopteris linearis* | 6 | 6 | 0 | 4 | 6 | 3 |
| *Desmodium ramosissimum* * | 6 | 6 | 0 | 5 | 4 | 1 |
| *Costularia melicoides* | 3 | 2 | 0 | 3 | 3 | 1 |
| *Lycopodiella cernua* | 9 | 7 | 1 | 8 | 5 | 4 |
| *Nephrolepis abrupta* | 9 | 6 | 0 | 4 | 7 | 5 |
| *Nephrolepis biserrata* | 3 | 3 | 0 | 3 | 0 | 2 |
| *Psidium cattleianum* * | 4 | 4 | 0 | 3 | 2 | 1 |
| *Stachytarpheta jamaicensis* * | 3 | 3 | 0 | 1 | 1 | 2 |
| *Stenotaphrum dimidiatum* * | 3 | 3 | 0 | 3 | 0 | 1 |
| *Tristemma mauritianum* | 3 | 2 | 0 | 3 | 0 | 3 |

| Plant species | Plaine-des-Palmistes | | | | | |
|---|---|---|---|---|---|---|
| | Number of samples | *Glomeromycotina* | *Mucoromycotina* | *Sebacinales* | *Helotiales* | *Cantharellales* |
| *Polytrichum commune* | 9 | 5 | 4 | 5 | 9 | 1 |
| *Ipomoea sp. 1* ** | 3 | 3 | 0 | 3 | 2 | 2 |
| *Dicranopteris linearis* | 3 | 3 | 1 | 1 | 2 | 1 |
| *Erica reunionensis* | 6 | 6 | 5 | 4 | 6 | 3 |
| *Juncus effusus* | 3 | 3 | 2 | 3 | 1 | 3 |
| *Hubertia ambavilla* | 1 | 1 | 1 | 0 | 1 | 0 |
| *Lycopodiella caroliniana* | 6 | 5 | 5 | 5 | 3 | 3 |
| *Lycopodiella cernua* | 9 | 7 | 8 | 8 | 8 | 2 |
| *Machaerina iridifolia* | 9 | 8 | 5 | 5 | 6 | 4 |
| *Phyllanthus phillyreifolius* | 1 | 1 | 1 | 1 | 1 | 0 |
| *Rhynchospora rugosa* ** | 9 | 8 | 1 | 6 | 5 | 2 |
| *Stoebe passerinoides* | 2 | 2 | 2 | 0 | 2 | 0 |
| *Tristemma mauritianum* | 3 | 3 | 1 | 1 | 2 | 1 |
| *Benthamia africana* | 9 | 8 | 6 | 6 | 5 | 4 |

| Plant species | Dimitile | | | | | |
|---|---|---|---|---|---|---|
| | Number of samples | *Glomeromycotina* | *Mucoromycotina* | *Sebacinales* | *Helotiales* | *Cantharellales* |
| *Polytrichum commune* | 7 | 3 | 0 | 5 | 6 | 1 |
| *Cordyline mauritiana* | 1 | 1 | 0 | 0 | 1 | 0 |
| *Dicranopteris linearis* | 10 | 10 | 0 | 3 | 7 | 2 |
| *Erica reunionensis* | 14 | 14 | 6 | 8 | 7 | 0 |
| *Habenaria frappieri* | 3 | 1 | 2 | 1 | 3 | 2 |
| *Lycopodium clavatum* | 12 | 2 | 0 | 8 | 10 | 0 |
| *Phylica nitida* | 15 | 15 | 8 | 5 | 13 | 2 |
| *Hyparrhenia rufa* | 9 | 8 | 2 | 4 | 7 | 1 |
| *Stoebe passerinoides* | 14 | 14 | 4 | 3 | 9 | 1 |

**Supplementary Table 2: Number of reads and OTUs:**

For each marker, we indicated the number of fungal Swarm OTUs and their coverage per root sample (after having discarded the contaminants using decontam), as well as the number of fungal OTUs that may form mycorrhizal interactions and their coverage per root sample. OTUs having less than 10 reads in the database were removed as they likely result from PCR or sequencing errors.

Note that even if both libraries had similar sizes, we significantly retrieved more fungal reads using the ITS2 marker than the 18S rRNA marker, because the AMADf-AMGDr primer pair also importantly amplified the plant 18S rRNA gene, whereas the ITS86F-ITS4 primer pair was more specific toward fungi only.

| OTU clustering | ITS2 region | 18S rRNA |
|---|---|---|
| Number of fungal OTUs | 5,236 | 4,371 |
| Total number of fungal reads | 14,105,171 | 4,523,549 |
| Mean number of fungal read per root sample (+/- the standard deviation) | 60,537 (+/-29,044) | 19,414 (+/-14,974) |
| Minimum and maximum number of fungal read per root sample | 7,346 – 209,608 | 18 – 77,640 |
| Number of fungal OTUs that may form mycorrhizal interactions | 622 | 1,177 |
| Total number of reads corresponding to OTUs that may form mycorrhizal interactions | 1,328,381 | 886,019 |
| Mean number of mycorrhizal fungal read per root sample (+/- s.d.) | 5,701 (+/-7,876) | 3,836 (+/-5,767) |
| Minimum and maximum number of mycorrhizal fungal read per root sample | 2 – 36,806 | 4 – 42,427 |

**Supplementary Table 3: PERMANOVA analyses revealed the significant effect of the plant taxonomic groups on the Bray-Curtis (a) or UniFrac (b) beta diversity between root samples for the different fungal lineages.**

The results of the PERMANOVA ($R^2$ and p-value based on 10,000 permutations) are indicated for all the fungi (based on the 18S rRNA or the ITS2 markers) or for each of the 5 main fungal lineages (*Glomeromycotina, Mucoromycotina, Sebacinales, Helotiales,* or *Cantharellales*) in each sampled community (Grand brûlé, Plaine-des-Palmistes, or Dimitile) using Swarm OTUs.

Significant correlations are highlighted in bold.

PERMANOVA could not be applied for *Mucoromycotina* in Grand brûlé because most OTUs were sample-specific.

Note that the non-significances of the PERMANOVA for the *Cantharellales* are likely due to the low number of corresponding OTUs in the different communities for this fungal lineage (Fig. 4).

**(a) Bray-Curtis dissimilarities**

| Fungi | Sampled community | Swarm OTUs | |
|---|---|---|---|
| | | $R^2$ | p-value |
| All 18S rRNA fungi | Grand brûlé | **0.15** | **0.0001** |
| | Plaine-des-Palmistes | **0.15** | **0.0002** |
| | Dimitile | **0.19** | **0.0001** |
| All ITS2 fungi | Grand brûlé | **0.08** | **0.04** |
| | Plaine-des-Palmistes | **0.17** | **0.0001** |
| | Dimitile | **0.15** | **0.0002** |
| *Glomeromycotina* | Grand brûlé | **0.11** | **0.0013** |
| | Plaine-des-Palmistes | **0.13** | **0.0361** |
| | Dimitile | **0.15** | **0.0005** |
| *Mucoromycotina* | Grand brûlé | NA | NA |
| | Plaine-des-Palmistes | **0.20** | **0.002** |
| | Dimitile | **0.22** | **0.0371** |
| *Sebacinales* | Grand brûlé | **0.09** | **0.0391** |
| | Plaine-des-Palmistes | 0.16 | 0.0768 |
| | Dimitile | **0.26** | **0.007** |
| *Helotiales* | Grand brûlé | 0.09 | 0.3255 |
| | Plaine-des-Palmistes | **0.18** | **0.0035** |
| | Dimitile | **0.18** | **0.0002** |
| *Cantharellales* | Grand brûlé | 0.16 | 0.1315 |
| | Plaine-des-Palmistes | 0.26 | 0.3109 |
| | Dimitile | 0.52 | 0.2617 |

**(b) UniFrac distances**

| Fungi | Sampled community | Swarm OTUs | |
|---|---|---|---|
| | | $R^2$ | p-value |
| All 18S rRNA fungi | Grand brûlé | **0.19** | **0.0003** |
| | Plaine-des-Palmistes | **0.23** | **0.0001** |
| | Dimitile | **0.22** | **0.0001** |
| All ITS2 fungi | Grand brûlé | **0.09** | **0.012** |
| | Plaine-des-Palmistes | **0.16** | **0.0007** |
| | Dimitile | **0.17** | **0.0008** |
| *Glomeromycotina* | Grand brûlé | **0.10** | **0.048** |
| | Plaine-des-Palmistes | 0.11 | 0.24 |
| | Dimitile | **0.16** | **0.008** |
| *Mucoromycotina* | Grand brûlé | NA | NA |
| | Plaine-des-Palmistes | 0.19 | 0.08 |
| | Dimitile | **0.33** | **0.006** |
| *Sebacinales* | Grand brûlé | 0.08 | 0.23 |
| | Plaine-des-Palmistes | 0.13 | 0.44 |
| | Dimitile | 0.26 | 0.065 |
| *Helotiales* | Grand brûlé | 0.10 | 0.27 |
| | Plaine-des-Palmistes | **0.23** | **0.006** |
| | Dimitile | **0.25** | **0.0009** |
| *Cantharellales* | Grand brûlé | **0.24** | **0.035** |
| | Plaine-des-Palmistes | 0.17 | 0.66 |
| | Dimitile | 0.64 | 0.10 |

**Supplementary Table 4: High proportion of shared OTUs between different plants species and different plant taxonomic groups.**

The proportion of shared OTUs between plant species or between plant taxonomic groups is indicated for all the fungi (based on the 18S rRNA or the ITS2 markers) or for each of the 5 main fungal lineages (*Glomeromycotina, Mucoromycotina, Sebacinales, Helotiales*, or *Cantharellales*) in each sampled community (Grand brûlé, Plaine-des-Palmistes, or Dimitile) using Swarm OTUs. These shared OTUs represent >90% of the total fungal reads.

| Fungi | Marker | Community | Proportion of fungal OTUs shared between plant species | Proportion of fungal OTUs shared between plant taxonomic groups |
|---|---|---|---|---|
| all | 18S | Grand brûlé | 0.43 | 0.38 |
| all | 18S | Plaine-des-Palmistes | 0.42 | 0.41 |
| all | 18S | Dimitile | 0.39 | 0.37 |
| all | ITS | Grand brûlé | 0.58 | 0.49 |
| all | ITS | Plaine-des-Palmistes | 0.52 | 0.51 |
| all | ITS | Dimitile | 0.48 | 0.47 |
| *Cantharellales* | ITS | Grand brûlé | 0.62 | 0.5 |
| *Cantharellales* | ITS | Plaine-des-Palmistes | 0.43 | 0.43 |
| *Cantharellales* | ITS | Dimitile | 0.29 | 0.29 |
| *Glomeromycotina* | 18S | Grand brûlé | 0.52 | 0.41 |
| *Glomeromycotina* | 18S | Plaine-des-Palmistes | 0.68 | 0.68 |
| *Glomeromycotina* | 18S | Dimitile | 0.5 | 0.48 |
| *Helotiales* | ITS | Grand brûlé | 0.58 | 0.58 |
| *Helotiales* | ITS | Plaine-des-Palmistes | 0.33 | 0.33 |
| *Helotiales* | ITS | Dimitile | 0.34 | 0.34 |
| *Mucoromycotina* | 18S | Grand brûlé | 0 | 0 |
| *Mucoromycotina* | 18S | Plaine-des-Palmistes | 0.38 | 0.38 |
| *Mucoromycotina* | 18S | Dimitile | 0.42 | 0.42 |
| *Sebacinales* | ITS | Grand brûlé | 0.54 | 0.49 |
| *Sebacinales* | ITS | Plaine-des-Palmistes | 0.68 | 0.64 |
| *Sebacinales* | ITS | Dimitile | 0.38 | 0.38 |

**Supplementary Table 5: The connectances of the plant-fungus networks are significantly lower than expected if plant-fungus interactions happen at random.** The connectance (the proportion of realized interactions in the network) is indicated for each plant-fungus network (*Glomeromycotina, Mucoromycotina, Sebacinales, Helotiales*, or *Cantharellales*) in each sampled community (Grand brûlé, Plaine-des-Palmistes, or Dimitile) using Swarm OTUs.

The significance of the connectance values was evaluated using shuffle-sample binary null models (the quasiswap null model keeps the connectance constant). We considered the network to be more connected (resp. less connected) than expected by chance if p<2.5%, where p is the fraction of the null model values that are greater than or equal to (resp. lower than or equal to) the observed value.

| Fungi | Site | Connectance | Connectance (binary null models) |
|---|---|---|---|
| Glomeromycotina | Grand brûlé | 0.23 | **Lower connectance (p=0)** |
| Glomeromycotina | Plaine des palmistes | 0.34 | Non significant (p>0.1) |
| Glomeromycotina | Dimitile | 0.27 | **Lower connectance (p=0)** |
| Mucoromycotina | Grand brûlé | 0.2 | Non significant (p=1) |
| Mucoromycotina | Plaine des palmistes | 0.14 | **Lower connectance (p=0.001)** |
| Mucoromycotina | Dimitile | 0.35 | Non significant (p>0.082) |
| Sebacinales | Grand brûlé | 0.2 | Non significant (p>0.035) |
| Sebacinales | Plaine des palmistes | 0.23 | Non significant (p>0.066) |
| Sebacinales | Dimitile | 0.23 | Non significant (p>0.393) |
| Helotiales | Grand brûlé | 0.29 | Non significant (p>0.057) |
| Helotiales | Plaine des palmistes | 0.18 | **Lower connectance (p=0)** |
| Helotiales | Dimitile | 0.23 | **Lower connectance (p=0.002)** |
| Cantharellales | Grand brûlé | 0.23 | Non significant (p>0.273) |
| Cantharellales | Plaine des palmistes | 0.21 | **Lower connectance (p=0.001)** |
| Cantharellales | Dimitile | 0.21 | Non significant (p>0.046) |

**Supplementary Table 6: The nestedness of the plant-fungus networks varies according to the fungal lineages**

Nestedness was computed for each plant-fungus network (*Glomeromycotina, Mucoromycotina, Sebacinales, Helotiales*, or *Cantharellales*) in each sampled community (Grand brûlé, Plaine-des-Palmistes, or Dimitile) using Swarm OTUs. The significance of the nestedness values was evaluated using both quasiswap (a) and shuffle-sample (b) null models and using abundance, incidence, or binary networks. Weighted NODF values were computed for quantified networks, whereas NODF2 values were computed for binary networks. We considered the network to be more nested (resp. less nested) than expected by chance if p<2.5%, where p is the fraction of the null model values that are greater than or equal to (resp. lower than or equal to) the observed value.

(a) quasiswap null models:

| **Fungi** | **Site** | **Nestedness (abundance null models)** | **Nestedness (incidence null models)** | **Nestedness (binary null models)** |
|---|---|---|---|---|
| Glomeromycotina | Grand brûlé | **Nested (p=0)** | **Nested (p=0)** | Non significant (p>0.387) |
| Glomeromycotina | Plaine des palmistes | Non significant (p>0.077) | Non significant (p>0.034) | Non significant (p>0.175) |
| Glomeromycotina | Dimitile | **Nested (p=0)** | **Nested (p=0)** | Non significant (p>0.234) |
| Mucoromycotina | Grand brûlé | Non significant (p=1) | Non significant (p=1) | Non significant (p=1) |
| Mucoromycotina | Plaine des palmistes | Non significant (p>0.093) | Non significant (p>0.135) | Non significant (p>0.442) |
| Mucoromycotina | Dimitile | **Nested (p=0.004)** | Non significant (p>0.268) | Non significant (p>0.43) |
| Sebacinales | Grand brûlé | Non significant (p>0.359) | Non significant (p>0.299) | Non significant (p>0.304) |
| Sebacinales | Plaine des palmistes | Non significant (p>0.089) | Non significant (p>0.159) | **Anti-nested (p=0.012)** |
| Sebacinales | Dimitile | **Nested (p=0.009)** | Non significant (p>0.487) | Non significant (p>0.395) |
| Helotiales | Grand brûlé | Non significant (p>0.33) | Non significant (p>0.264) | Non significant (p>0.099) |
| Helotiales | Plaine des palmistes | **Nested (p=0)** | Non significant (p>0.136) | Non significant (p>0.496) |
| Helotiales | Dimitile | **Nested (p=0)** | Non significant (p>0.052) | **Nested (p=0.017)** |
| Cantharellales | Grand brûlé | **Anti-nested (p=0.002)** | Non significant (p>0.481) | Non significant (p>0.379) |
| Cantharellales | Plaine des palmistes | Non significant (p>0.374) | Non significant (p>0.288) | **Anti-nested (p=0.025)** |
| Cantharellales | Dimitile | Non significant (p>0.883) | Non significant (p>0.728) | Non significant (p>0.119) |

(b) shuffle-sample null models:

| Fungi | Site | Nestedness (abundance null models) | Nestedness (incidence null models) | Nestedness (binary null models) |
|---|---|---|---|---|
| Glomeromycotina | Grand brûlé | Non significant (p>0.356) | **Nested (p=0.024)** | Non significant (p>0.276) |
| Glomeromycotina | Plaine des palmistes | Non significant (p>0.068) | Non significant (p>0.456) | Non significant (p>0.21) |
| Glomeromycotina | Dimitile | Non significant (p>0.072) | **Nested (p=0.001)** | Non significant (p>0.037) |
| Mucoromycotina | Grand brûlé | Non significant (p=1) | Non significant (p=1) | Non significant (p=1) |
| Mucoromycotina | Plaine des palmistes | Non significant (p>0.148) | Non significant (p>0.106) | Non significant (p>0.209) |
| Mucoromycotina | Dimitile | Non significant (p>0.368) | Non significant (p>0.437) | Non significant (p>0.394) |
| Sebacinales | Grand brûlé | Non significant (p>0.432) | Non significant (p>0.43) | Non significant (p>0.257) |
| Sebacinales | Plaine des palmistes | Non significant (p>0.403) | Non significant (p>0.237) | Non significant (p>0.264) |
| Sebacinales | Dimitile | Non significant (p>0.15) | Non significant (p>0.191) | Non significant (p>0.234) |
| Helotiales | Grand brûlé | Non significant (p>0.081) | Non significant (p>0.228) | **Anti-nested (p=0.025)** |
| Helotiales | Plaine des palmistes | Non significant (p>0.25) | Non significant (p>0.141) | Non significant (p>0.362) |
| Helotiales | Dimitile | Non significant (p>0.176) | Non significant (p>0.412) | Non significant (p>0.08) |
| Cantharellales | Grand brûlé | Non significant (p>0.042) | Non significant (p>0.487) | Non significant (p>0.312) |
| Cantharellales | Plaine des palmistes | Non significant (p>0.056) | Non significant (p>0.148) | **Anti-nested (p=0.003)** |
| Cantharellales | Dimitile | Non significant (p>0.037) | Non significant (p=1) | **Anti-nested (p=0.022)** |

**Supplementary Table 7: The checkerboard score (C-score) of the plant-fungus networks varies according to the fungal lineages**

The checkerboard score (C-score) is indicated for each plant-fungus (binary) network (*Glomeromycotina, Mucoromycotina, Sebacinales, Helotiales*, or *Cantharellales*) in each sampled community (Grand brûlé, Plaine-des-Palmistes, or Dimitile) using Swarm OTUs.

The significance of the checkerboard scores was evaluated using both quasiswap (a) and shuffle-sample (b) null models and using abundance, incidence, or binary networks. We considered the network to have a significantly higher checkerboard score (resp. lower checkerboard score) than expected by chance if p<2.5%, where p is the fraction of the null model values that are greater than or equal to (resp. lower than or equal to) the observed value.

(a) quasiswap null models:

| **Fungi** | **Site** | **Cscore** | **Cscore (abundance null models)** | **Cscore (incidence null models)** | **Cscore (binary null models)** |
|---|---|---|---|---|---|
| Glomeromycotina | Grand brûlé | 0.3 | **Anti-checkerboard (p=0)** | **Anti-checkerboard (p=0)** | Non significant (p>0.287) |
| Glomeromycotina | Plaine des palmistes | 0.17 | **Anti-checkerboard (p=0)** | Non significant (p>0.128) | Non significant (p>0.193) |
| Glomeromycotina | Dimitile | 0.17 | **Anti-checkerboard (p=0)** | **Anti-checkerboard (p=0.005)** | Non significant (p>0.039) |
| Mucoromycotina | Grand brûlé | 1 | Non significant (p=1) | Non significant (p=1) | Non significant (p=1) |
| Mucoromycotina | Plaine des palmistes | 0.63 | **Anti-checkerboard (p=0)** | Non significant (p>0.08) | Non significant (p>0.376) |
| Mucoromycotina | Dimitile | 0.28 | Non significant (p>0.036) | Non significant (p>0.144) | Non significant (p>0.11) |
| Sebacinales | Grand brûlé | 0.43 | **Anti-checkerboard (p=0.001)** | Non significant (p>0.047) | Non significant (p>0.34) |
| Sebacinales | Plaine des palmistes | 0.54 | Non significant (p>0.392) | Non significant (p>0.224) | **Checkerboard (p=0.018)** |
| Sebacinales | Dimitile | 0.42 | Non significant (p>0.099) | Non significant (p>0.354) | Non significant (p>0.137) |
| Helotiales | Grand brûlé | 0.29 | Non significant (p>0.107) | Non significant (p>0.277) | Non significant (p>0.303) |
| Helotiales | Plaine des palmistes | 0.29 | **Anti-checkerboard (p=0)** | **Anti-checkerboard (p=0.007)** | **Anti-checkerboard (p=0.019)** |
| Helotiales | Dimitile | 0.25 | **Anti-checkerboard (p=0)** | **Anti-checkerboard (p=0.002)** | Non significant (p>0.4) |
| Cantharellales | Grand brûlé | 0.53 | **Anti-checkerboard (p=0.015)** | Non significant (p>0.226) | Non significant (p>0.07) |
| Cantharellales | Plaine des palmistes | 0.47 | **Anti-checkerboard (p=0.022)** | Non significant (p>0.129) | Non significant (p>0.029) |
| Cantharellales | Dimitile | 0.87 | Non significant (p>0.694) | Non significant (p>0.597) | Non significant (p>0.587) |

(b) shuffle-sample null models:

| Fungi | Site | Cscore | Cscore (abundance null models) | Cscore (incidence null models) | Cscore (binary null models) |
|---|---|---|---|---|---|
| Glomeromycotina | Grand brûlé | 0.3 | **Checkerboard (p=0.003)** | **Checkerboard (p=0.003)** | **Checkerboard (p=0.003)** |
| Glomeromycotina | Plaine des palmistes | 0.17 | Non significant (p>0.228) | Non significant (p>0.228) | Non significant (p>0.228) |
| Glomeromycotina | Dimitile | 0.17 | Non significant (p>0.365) | Non significant (p>0.365) | Non significant (p>0.365) |
| Mucoromycotina | Grand brûlé | 1 | Non significant (p=1) | Non significant (p=1) | Non significant (p=1) |
| Mucoromycotina | Plaine des palmistes | 0.63 | Non significant (p>0.263) | Non significant (p>0.263) | Non significant (p>0.263) |
| Mucoromycotina | Dimitile | 0.28 | Non significant (p>0.215) | Non significant (p>0.215) | Non significant (p>0.215) |
| Sebacinales | Grand brûlé | 0.43 | Non significant (p>0.493) | Non significant (p>0.493) | Non significant (p>0.493) |
| Sebacinales | Plaine des palmistes | 0.54 | Non significant (p>0.042) | Non significant (p>0.042) | Non significant (p>0.042) |
| Sebacinales | Dimitile | 0.42 | Non significant (p>0.028) | Non significant (p>0.028) | Non significant (p>0.028) |
| Helotiales | Grand brûlé | 0.29 | **Checkerboard (p=0.018)** | **Checkerboard (p=0.018)** | **Checkerboard (p=0.018)** |
| Helotiales | Plaine des palmistes | 0.29 | Non significant (p>0.368) | Non significant (p>0.368) | Non significant (p>0.368) |
| Helotiales | Dimitile | 0.25 | Non significant (p>0.293) | Non significant (p>0.293) | Non significant (p>0.293) |
| Cantharellales | Grand brûlé | 0.53 | Non significant (p>0.371) | Non significant (p>0.371) | Non significant (p>0.371) |
| Cantharellales | Plaine des palmistes | 0.47 | **Checkerboard (p=0.004)** | **Checkerboard (p=0.004)** | **Checkerboard (p=0.004)** |
| Cantharellales | Dimitile | 0.87 | Non significant (p>0.051) | Non significant (p>0.051) | Non significant (p>0.051) |

**Supplementary Table 8: The modularity of the plant-fungus networks varies according to the fungal lineages**

The modularity value (proportion of interactions within modules) is indicated for each plant-fungus network (*Glomeromycotina, Mucoromycotina, Sebacinales, Helotiales,* or *Cantharellales*) in each sampled community (Grand brûlé, Plaine-des-Palmistes, or Dimitile) using Swarm OTUs.

The significance of the modularity values was evaluated using both quasiswap (a) and shuffle-sample (b) null models and using abundance, incidence, or binary networks.

We considered the network to be more modular (resp. less modular) than expected by chance if p<2.5%, where p is the fraction of the null model values that are greater than or equal to (resp. lower than or equal to) the observed value.

(a) quasiswap null models:

| **Fungi** | **Site** | **Modularity (abundance null models)** | **Modularity (incidence null models)** | **Modularity (binary null models)** |
|---|---|---|---|---|
| Glomeromycotina | Grand brûlé | M=0.37; Non significant (p>0.107) | M=0.29; Non significant (p>0.117) | **M=0.32; Modular (p=0)** |
| Glomeromycotina | Plaine des palmistes | M=0.32; Non significant (p>0.091) | M=0.21; Non significant (p>0.139) | M=0.25; Non significant (p>0.177) |
| Glomeromycotina | Dimitile | M=0.19; Non significant (p>0.145) | M=0.19; Non significant (p>0.073) | M=0.32; Non significant (p>0.486) |
| Mucoromycotina | Grand brûlé | M=0.71; Non significant (p>0.512) | M=0.75; Non significant (p=1) | M=0.75; Non significant (p=1) |
| Mucoromycotina | Plaine des palmistes | M=0.59; Non significant (p>0.086) | M=0.46; Non significant (p>0.133) | M=0.51; Non significant (p>0.42) |
| Mucoromycotina | Dimitile | M=0.47; Non significant (p>0.038) | M=0.35; Non significant (p>0.174) | M=0.38; Non significant (p>0.406) |
| Sebacinales | Grand brûlé | M=0.46; Non significant (p>0.288) | **M=0.4; Modular (p=0.017)** | M=0.37; Non significant (p>0.135) |
| Sebacinales | Plaine des palmistes | M=0.4; Non significant (p>0.373) | M=0.38; Non significant (p>0.086) | **M=0.4; Modular (p=0.009)** |
| Sebacinales | Dimitile | M=0.27; Non significant (p>0.346) | M=0.29; Non significant (p>0.48) | M=0.47; Non significant (p>0.465) |
| Helotiales | Grand brûlé | **M=0.16; Modular (p=0.02)** | M=0.27; Non significant (p>0.209) | M=0.37; Non significant (p>0.048) |
| Helotiales | Plaine des palmistes | M=0.4; Non significant (p>0.051) | M=0.36; Non significant (p>0.093) | M=0.41; Non significant (p>0.1) |
| Helotiales | Dimitile | **M=0.42; Modular (p=0.017)** | M=0.34; Non significant (p>0.224) | M=0.4; Non significant (p>0.263) |
| Cantharellales | Grand brûlé | **M=0.69; Modular (p=0)** | M=0.49; Non significant (p>0.217) | M=0.47; Non significant (p>0.33) |
| Cantharellales | Plaine des palmistes | **M=0.55; Modular (p=0.017)** | M=0.44; Non significant (p>0.488) | M=0.55; Non significant (p>0.295) |
| Cantharellales | Dimitile | M=0.52; Non significant (p>0.116) | M=0.69; Non significant (p>0.186) | M=0.72; Non significant (p>0.084) |

(b) shuffle-sample null models:

| Fungi | Site | Modularity (abundance null models) | Modularity (incidence null models) | Modularity (binary null models) |
|---|---|---|---|---|
| Glomeromycotina | Grand brûlé | **M=0.37; Modular (p=0.009)** | **M=0.29; Modular (p=0)** | **M=0.32; Modular (p=0.001)** |
| Glomeromycotina | Plaine des palmistes | M=0.32; Non significant (p>0.127) | M=0.21; Non significant (p>0.028) | M=0.25; Non significant (p>0.467) |
| Glomeromycotina | Dimitile | M=0.19; Non significant (p>0.063) | M=0.19; Non significant (p>0.144) | M=0.32; Non significant (p>0.143) |
| Mucoromycotina | Grand brûlé | **M=0.71; Modular (p=0)** | M=0.75; Non significant (p=1) | M=0.75; Non significant (p=1) |
| Mucoromycotina | Plaine des palmistes | M=0.59; Non significant (p>0.325) | M=0.46; Non significant (p>0.087) | **M=0.51; Modular (p=0.017)** |
| Mucoromycotina | Dimitile | **M=0.47; Modular (p=0.014)** | **M=0.35; Modular (p=0.023)** | M=0.38; Non significant (p>0.3) |
| Sebacinales | Grand brûlé | M=0.46; Non significant (p>0.188) | M=0.4; Non significant (p>0.056) | M=0.37; Non significant (p>0.348) |
| Sebacinales | Plaine des palmistes | M=0.4; Non significant (p>0.072) | M=0.38; Non significant (p>0.169) | M=0.4; Non significant (p>0.23) |
| Sebacinales | Dimitile | M=0.27; Non significant (p>0.472) | M=0.29; Non significant (p>0.411) | M=0.47; Non significant (p>0.077) |
| Helotiales | Grand brûlé | M=0.16; Non significant (p>0.082) | M=0.27; Non significant (p>0.221) | M=0.37; Non significant (p>0.408) |
| Helotiales | Plaine des palmistes | M=0.4; Non significant (p>0.129) | **M=0.36; Modular (p=0.015)** | **M=0.41; Modular (p=0.007)** |
| Helotiales | Dimitile | **M=0.42; Modular (p=0.009)** | **M=0.34; Modular (p=0.001)** | M=0.4; Non significant (p>0.209) |
| Cantharellales | Grand brûlé | **M=0.69; Modular (p=0.017)** | M=0.49; Non significant (p>0.104) | M=0.47; Non significant (p>0.153) |
| Cantharellales | Plaine des palmistes | M=0.55; Non significant (p>0.067) | **M=0.44; Modular (p=0.013)** | M=0.55; Non significant (p>0.082) |
| Cantharellales | Dimitile | M=0.52; Non significant (p>0.241) | **M=0.69; Modular (p=0.025)** | **M=0.72; Modular (p=0.014)** |

#### Supplementary Figures:

**Supplementary Figure 1: Negative controls indicate low baseline contamination in our dataset**

(a, c) Numbers of reads found for the 8 combinations of tagged primers left empty indicating the mean number of index hopping for the 18S rRNA marker (a) or the ITS2 marker (c).

(b, d) Number of reads found for the PCR negative controls indicating the mean number of reads corresponding to contaminations for the 18S rRNA marker (d) or the ITS2 marker (d).

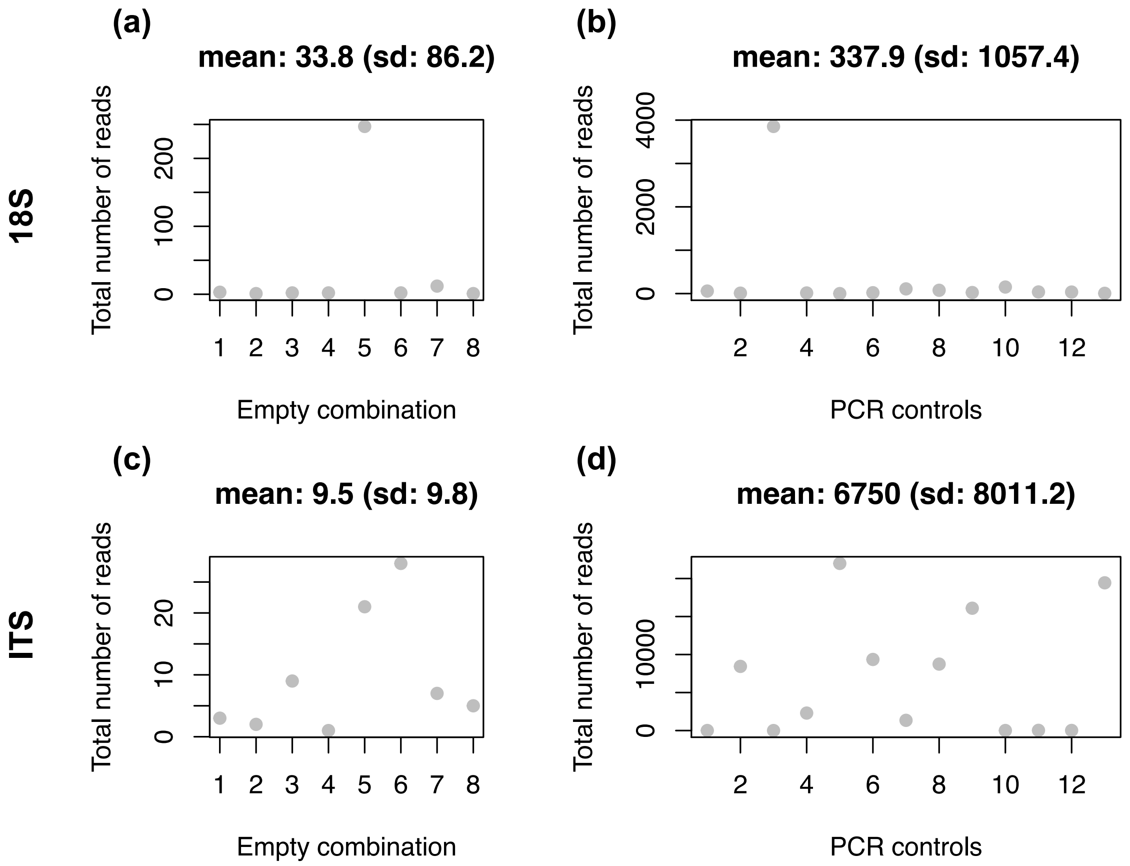
(a)
mean: 33.8 (sd: 86.2)
(b)
mean: 337.9 (sd: 1057.4)
18S
Total number of reads
Empty combination
PCR controls
(c)
mean: 9.5 (sd: 9.8)
(d)
mean: 6750 (sd: 8011.2)
ITS

**Supplementary Figure 2: Number of fungal reads per sample:**

Each boxplot represents the number of reads corresponding to fungal lineages that may form mycorrhizal interactions in the plant root samples of a given plant species using the 18S rRNA (a) or the ITS2 (b) markers. Each panel corresponds to a sampled community (Grand brûlé, Plaine-des-Palmistes, or Dimitile).

Boxplots present the median surrounded by the first and third quartiles, and whiskers extend to the extreme values but no further than 1.5 of the inter-quartile range.

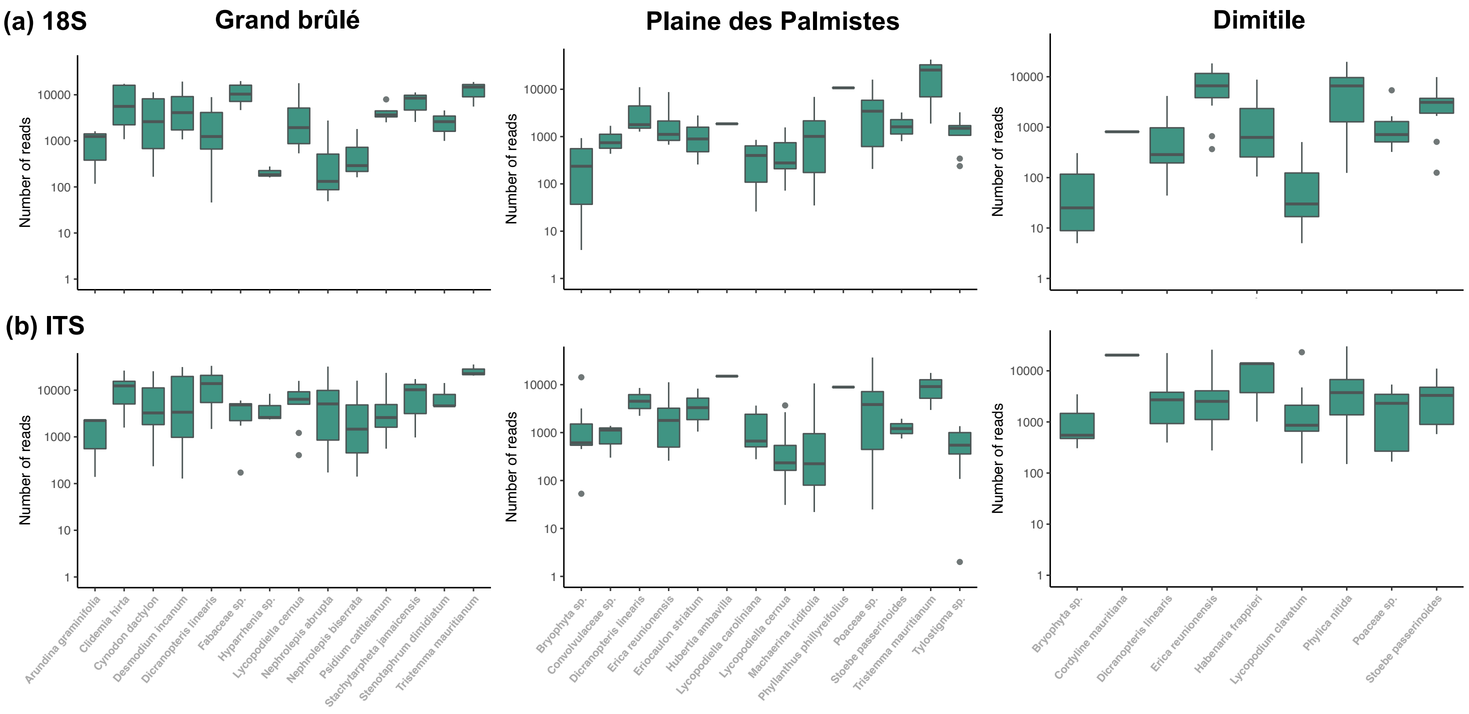

**Supplementary Figure 3: Root mycobiome compositions vary according to the main plant taxonomic groups**

For each plant root sample belonging to a given plant group (bryophytes, lycopods, ferns, monocots (excluding *Orchidaceae*), eudicots (excluding *Ericaceae*), *Orchidaceae*, or *Ericaceae*), the relative abundance of the fungi is indicated according to the 18S rRNA (a) or ITS2 (b) markers.

The class and the order of each fungus are represented in colors. Rare taxa (representing less than 0.1% of the data are represented in black). The sampled community where each sample came from is indicated at the bottom of the bar charts. Samples with less than 20 reads were discarded (empty space).

(I) Bryophytes

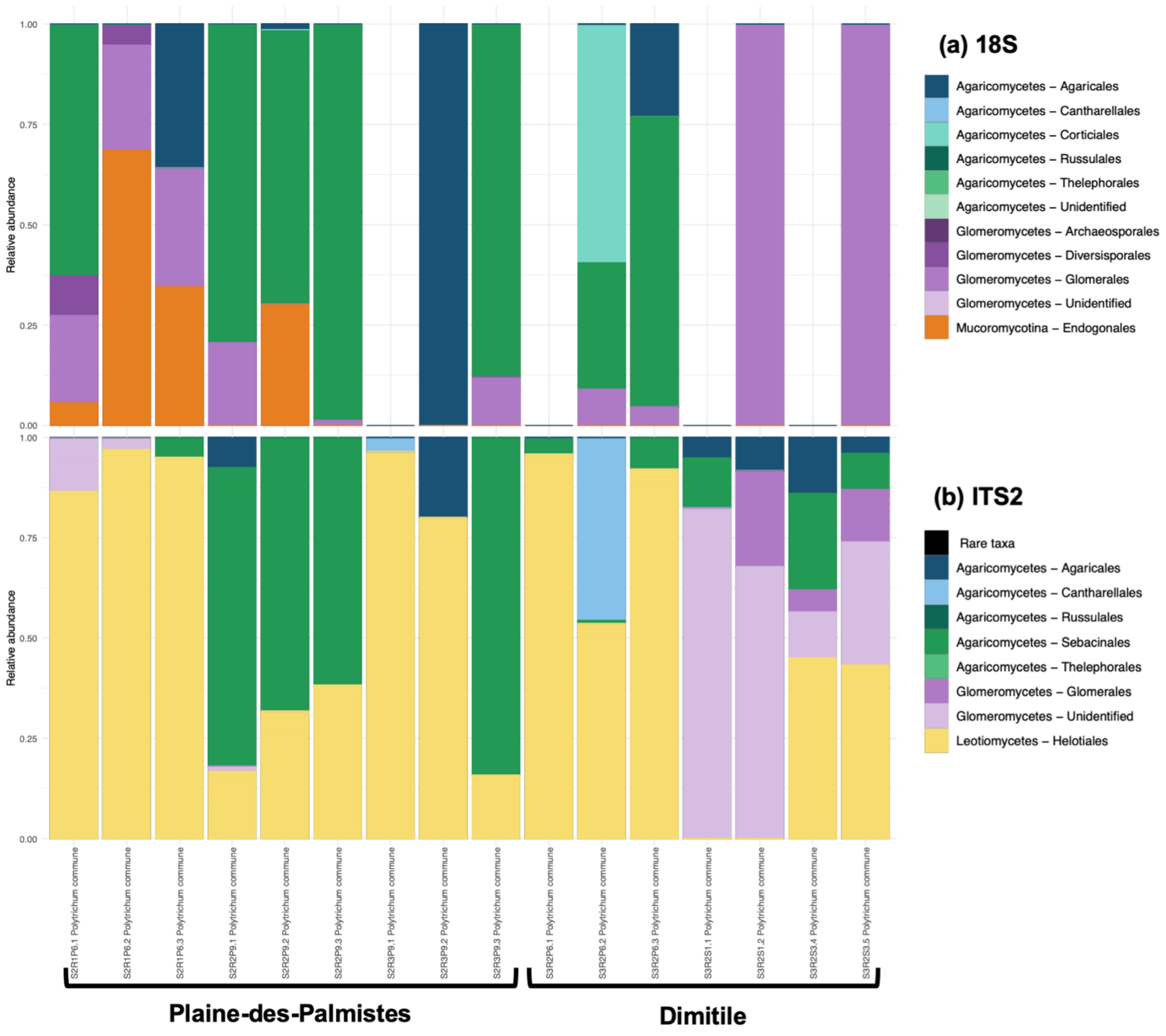

(II) Lycopods

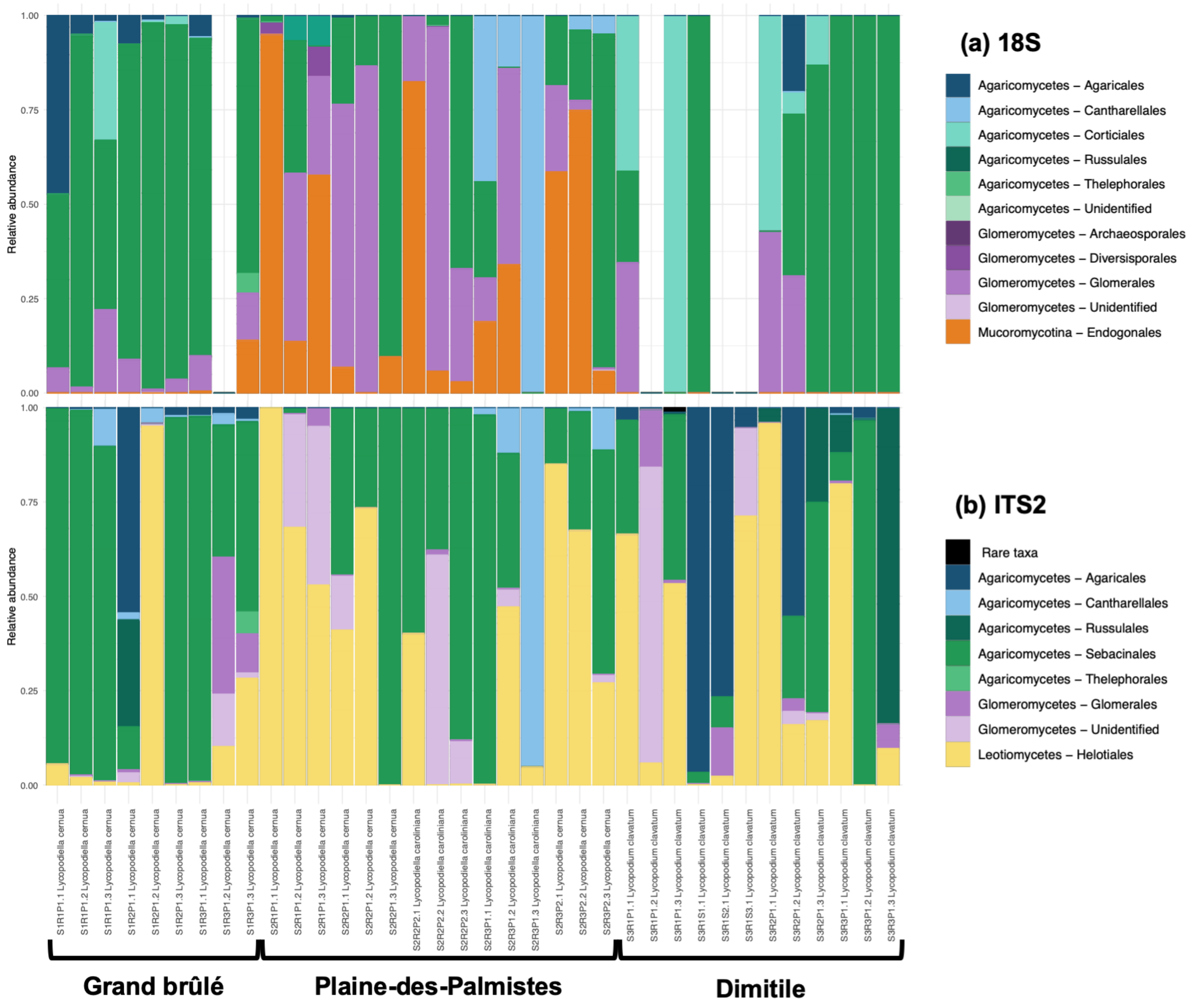
(a) 18S
Agaricomycetes – Agaricales
Agaricomycetes – Cantharellales
Agaricomycetes – Corticiales
Agaricomycetes – Russulales
Agaricomycetes – Thelephorales
Agaricomycetes – Unidentified
Glomeromycetes – Archaeosporales
Glomeromycetes – Diversisporales
Glomeromycetes – Glomerales
Glomeromycetes – Unidentified
Mucoromycotina – Endogonales
(b) ITS2
Rare taxa
Agaricomycetes – Agaricales
Agaricomycetes – Cantharellales
Agaricomycetes – Russulales
Agaricomycetes – Sebacinales
Agaricomycetes – Thelephorales
Glomeromycetes – Glomerales
Glomeromycetes – Unidentified
Leotiomycetes – Helotiales
Relative abundance
1.00
0.75
0.50
0.25
0.00
S1R1P1.1 Lycopodiella cernua
S1R1P1.2 Lycopodiella cernua
S1R1P1.3 Lycopodiella cernua
S1R2P1.1 Lycopodiella cernua
S1R2P1.2 Lycopodiella cernua
S1R2P1.3 Lycopodiella cernua
S1R3P1.1 Lycopodiella cernua
S1R3P1.2 Lycopodiella cernua
S1R3P1.3 Lycopodiella cernua
S2R1P1.1 Lycopodiella cernua
S2R1P1.2 Lycopodiella cernua
S2R1P1.3 Lycopodiella cernua
S2R2P1.1 Lycopodiella cernua
S2R2P1.2 Lycopodiella cernua
S2R2P1.3 Lycopodiella cernua
S2R2P2.1 Lycopodiella caroliniana
S2R2P2.2 Lycopodiella caroliniana
S2R2P2.3 Lycopodiella caroliniana
S2R3P1.1 Lycopodiella caroliniana
S2R3P1.2 Lycopodiella caroliniana
S2R3P1.3 Lycopodiella caroliniana
S2R3P2.1 Lycopodiella cernua
S2R3P2.2 Lycopodiella cernua
S2R3P2.3 Lycopodiella cernua
S3R1P1.1 Lycopodium clavatum
S3R1P1.2 Lycopodium clavatum
S3R1P1.3 Lycopodium clavatum
S3R1S1.1 Lycopodium clavatum
S3R1S2.1 Lycopodium clavatum
S3R1S3.1 Lycopodium clavatum
S3R2P1.1 Lycopodium clavatum
S3R2P1.2 Lycopodium clavatum
S3R2P1.3 Lycopodium clavatum
S3R3P1.1 Lycopodium clavatum
S3R3P1.2 Lycopodium clavatum
S3R3P1.3 Lycopodium clavatum
Grand brûlé
Plaine-des-Palmistes
Dimitile

(III) Ferns

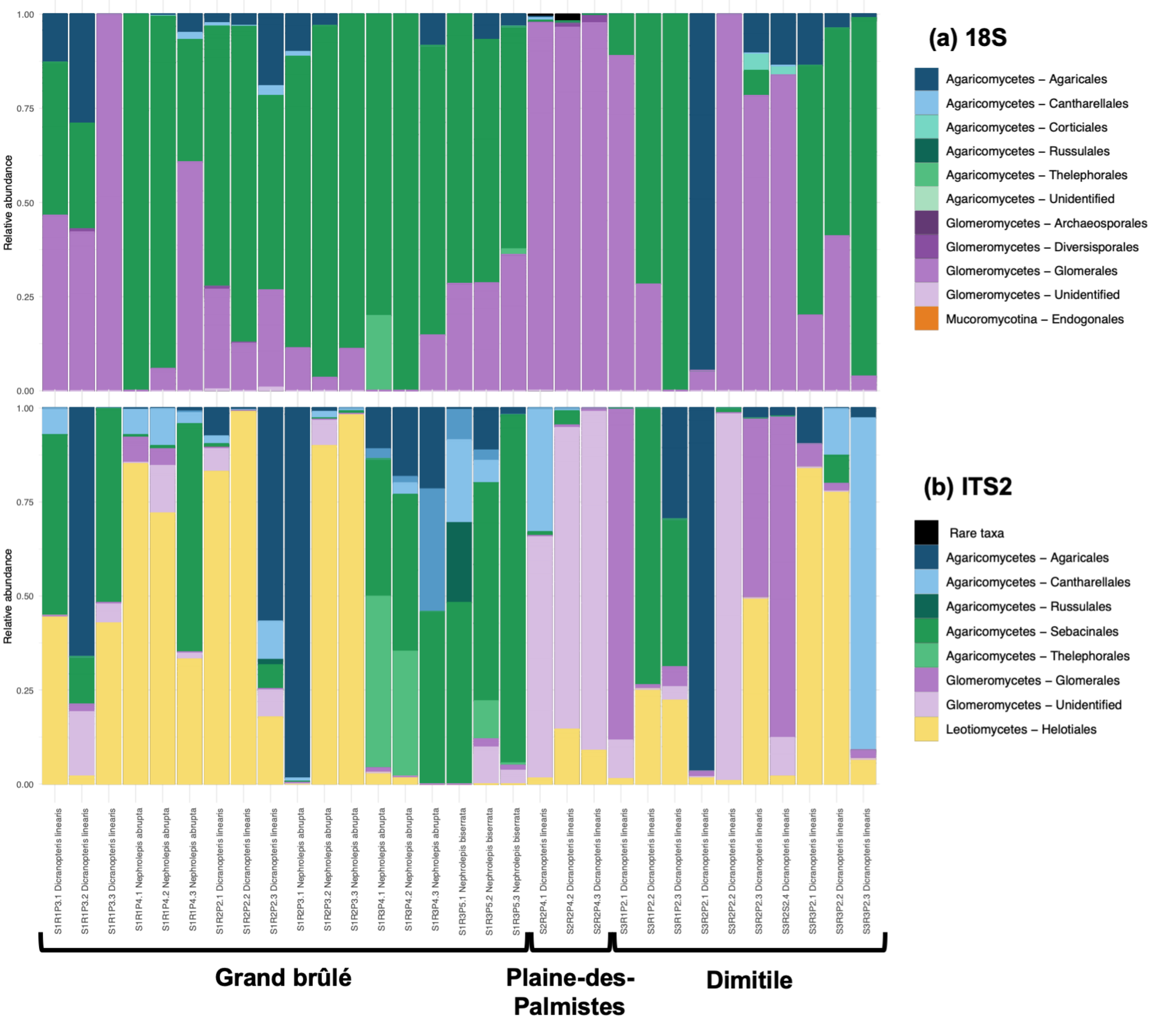
(a) 18S
Agaricomycetes – Agaricales
Agaricomycetes – Cantharellales
Agaricomycetes – Corticiales
Agaricomycetes – Russulales
Agaricomycetes – Thelephorales
Agaricomycetes – Unidentified
Glomeromycetes – Archaeosporales
Glomeromycetes – Diversisporales
Glomeromycetes – Glomerales
Glomeromycetes – Unidentified
Mucoromycotina – Endogonales
(b) ITS2
Rare taxa
Agaricomycetes – Agaricales
Agaricomycetes – Cantharellales
Agaricomycetes – Russulales
Agaricomycetes – Sebacinales
Agaricomycetes – Thelephorales
Glomeromycetes – Glomerales
Glomeromycetes – Unidentified
Leotiomycetes – Helotiales
Relative abundance
1.00
0.75
0.50
0.25
0.00
S1R1P3.1 Dicranopteris linearis
S1R1P3.2 Dicranopteris linearis
S1R1P3.3 Dicranopteris linearis
S1R1P4.1 Nephrolepis abrupta
S1R1P4.2 Nephrolepis abrupta
S1R1P4.3 Nephrolepis abrupta
S1R2P2.1 Dicranopteris linearis
S1R2P2.2 Dicranopteris linearis
S1R2P2.3 Dicranopteris linearis
S1R2P3.1 Nephrolepis abrupta
S1R2P3.2 Nephrolepis abrupta
S1R2P3.3 Nephrolepis abrupta
S1R3P4.1 Nephrolepis abrupta
S1R3P4.2 Nephrolepis abrupta
S1R3P4.3 Nephrolepis abrupta
S1R3P5.1 Nephrolepis biserrata
S1R3P5.2 Nephrolepis biserrata
S1R3P5.3 Nephrolepis biserrata
S2R2P4.1 Dicranopteris linearis
S2R2P4.2 Dicranopteris linearis
S2R2P4.3 Dicranopteris linearis
S3R1P2.1 Dicranopteris linearis
S3R1P2.2 Dicranopteris linearis
S3R1P2.3 Dicranopteris linearis
S3R2P2.1 Dicranopteris linearis
S3R2P2.2 Dicranopteris linearis
S3R2P2.3 Dicranopteris linearis
S3R2S2.4 Dicranopteris linearis
S3R3P2.1 Dicranopteris linearis
S3R3P2.2 Dicranopteris linearis
S3R3P2.3 Dicranopteris linearis
Grand brûlé
Plaine-des-Palmistes
Dimitile

(IV) Monocots (excluding *Orchidaceae*)

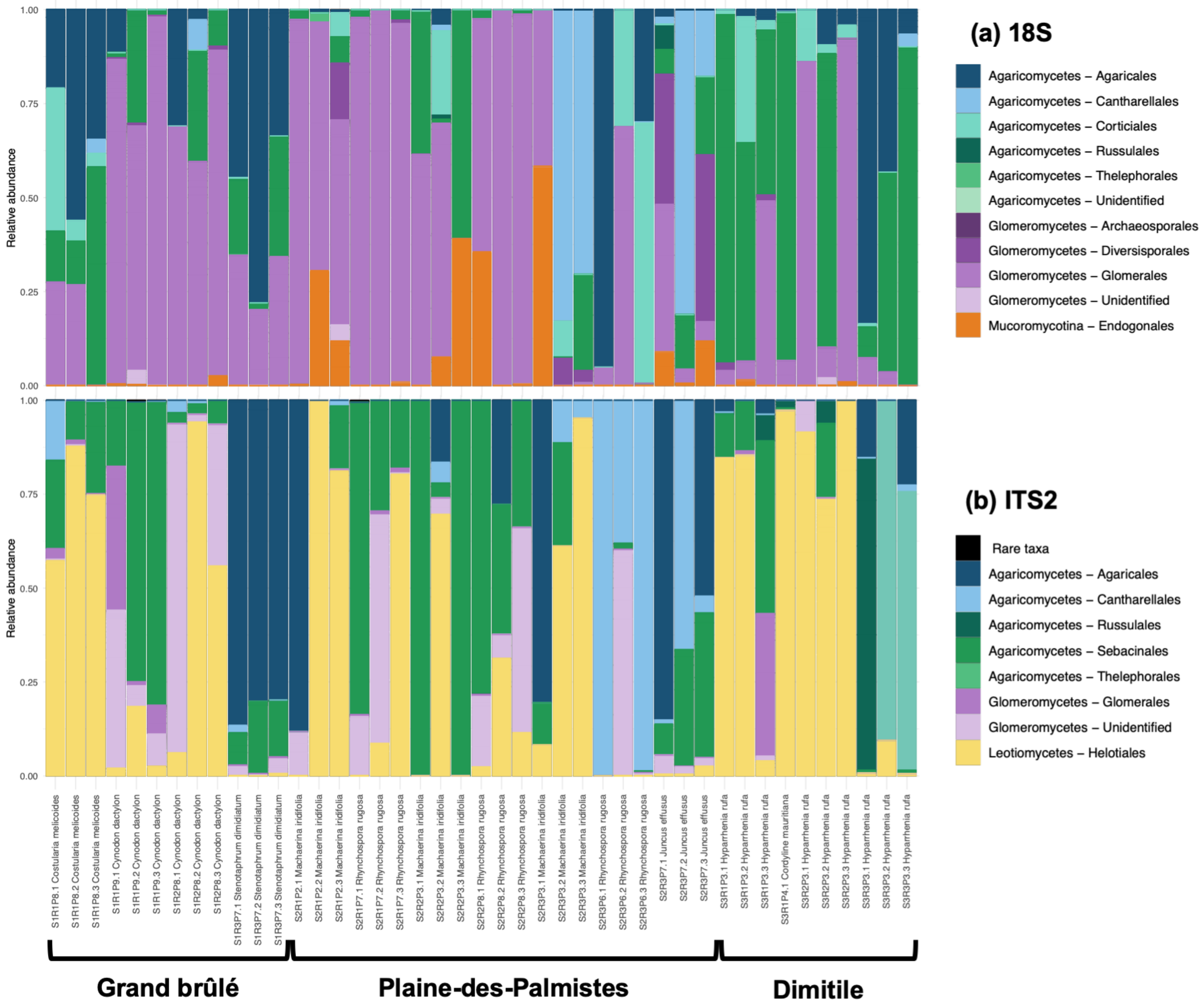

(V) Eudicots (excluding *Ericaceae*)

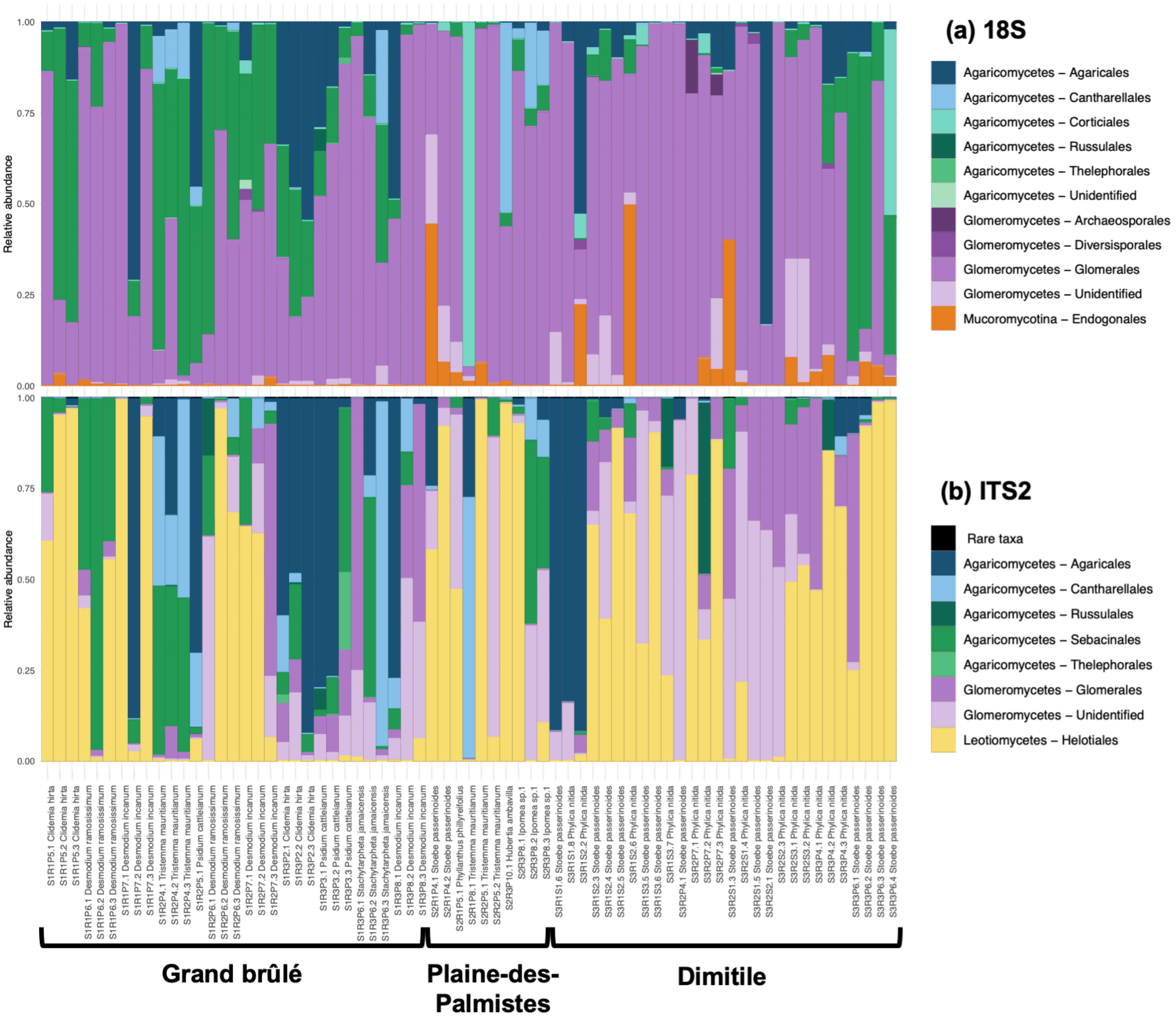

(VI) *Orchidaceae*

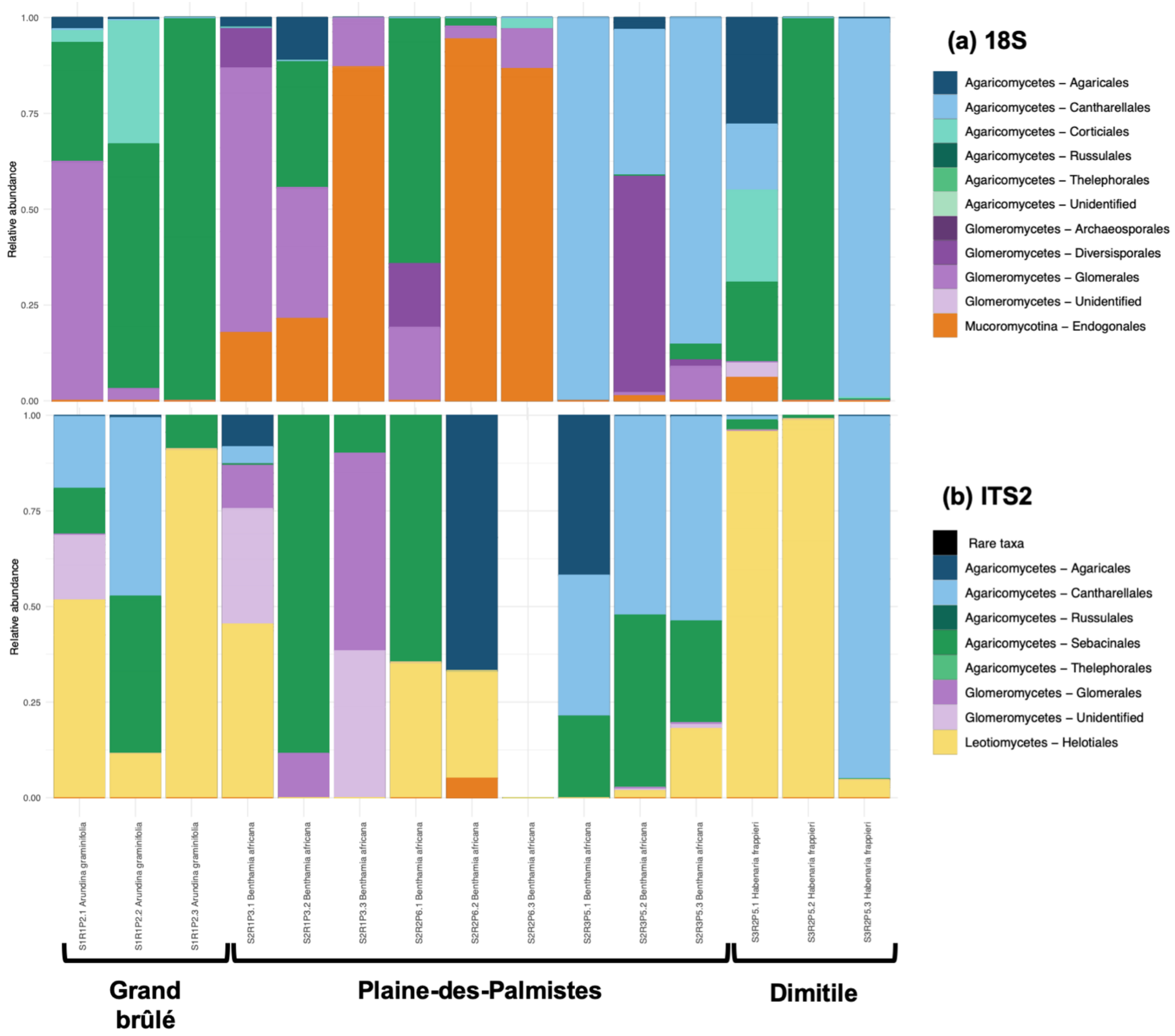

(VII) *Ericaceae*

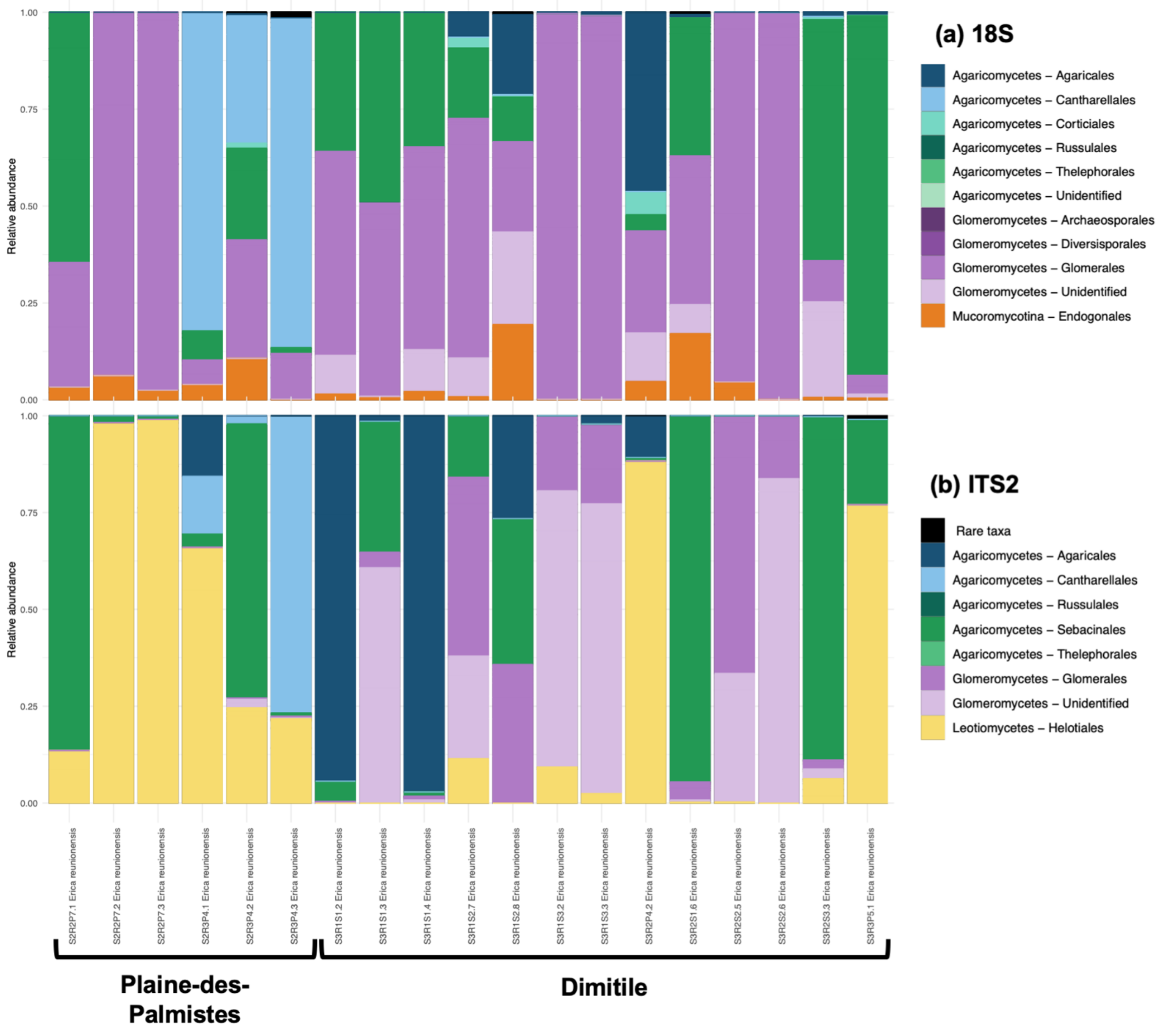

**Supplementary Figure 4: Relative read abundances and sample incidence are correlated measures of weighted interactions**

For each fungal lineage (*Glomeromycotina, Mucoromycotina, Sebacinales, Helotiales,* or *Cantharellales*) and in each sampled community (Grand brûlé, Plaine-des-Palmistes, or Dimitile), we tested the relationship between the weighted interactions measured as the relative read abundances (y-axis) or measured as the incidence among samples (x-axis). The significance of each relationship evaluated using a linear model is indicated at the top of each panel.

This was not tested for plant-*Mucoromycotina* networks in Grand brûlé given that each interaction was only found in one sample.

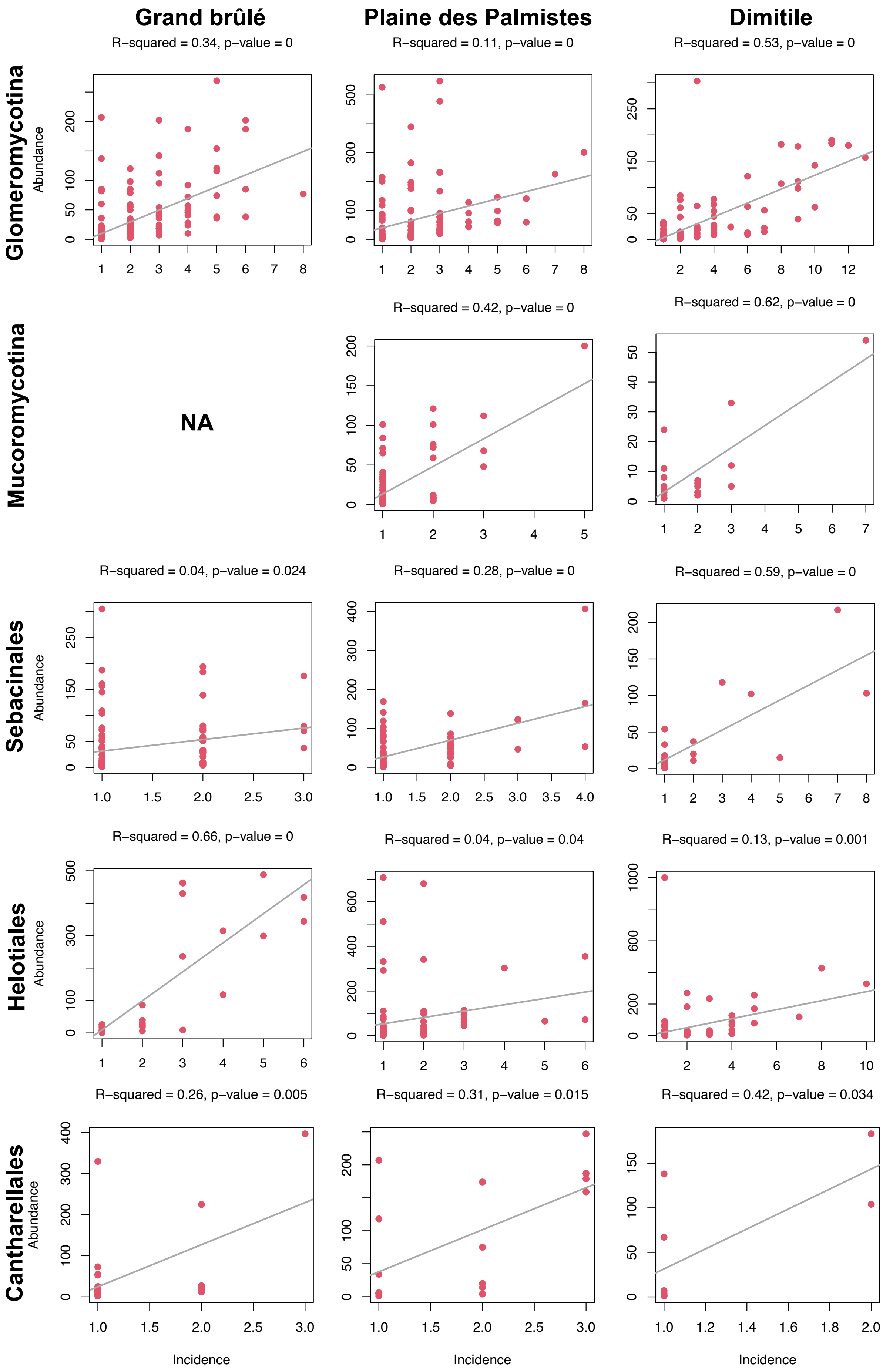
Grand brûlé
Plaine des Palmistes
Dimitile
R−squared = 0.34, p−value = 0
R−squared = 0.11, p−value = 0
R−squared = 0.53, p−value = 0
Glomeromycotina
Abundance
Mucoromycotina
NA
R−squared = 0.42, p−value = 0
R−squared = 0.62, p−value = 0
Sebacinales
R−squared = 0.04, p−value = 0.024
R−squared = 0.28, p−value = 0
R−squared = 0.59, p−value = 0
Helotiales
R−squared = 0.66, p−value = 0
R−squared = 0.04, p−value = 0.04
R−squared = 0.13, p−value = 0.001
Cantharellales
R−squared = 0.26, p−value = 0.005
R−squared = 0.31, p−value = 0.015
R−squared = 0.42, p−value = 0.034
Incidence

**Supplementary Figure 5: Rarefaction plots indicate that we did not document the whole root fungal diversity for many plant species**

Rarefactions plots indicated the mean number of fungal OTUs (OTU richness) or the mean phylogenetic diversity of the fungal OTUs (Faith’s index) found in each plant species as a function of the number of individuals sampled.

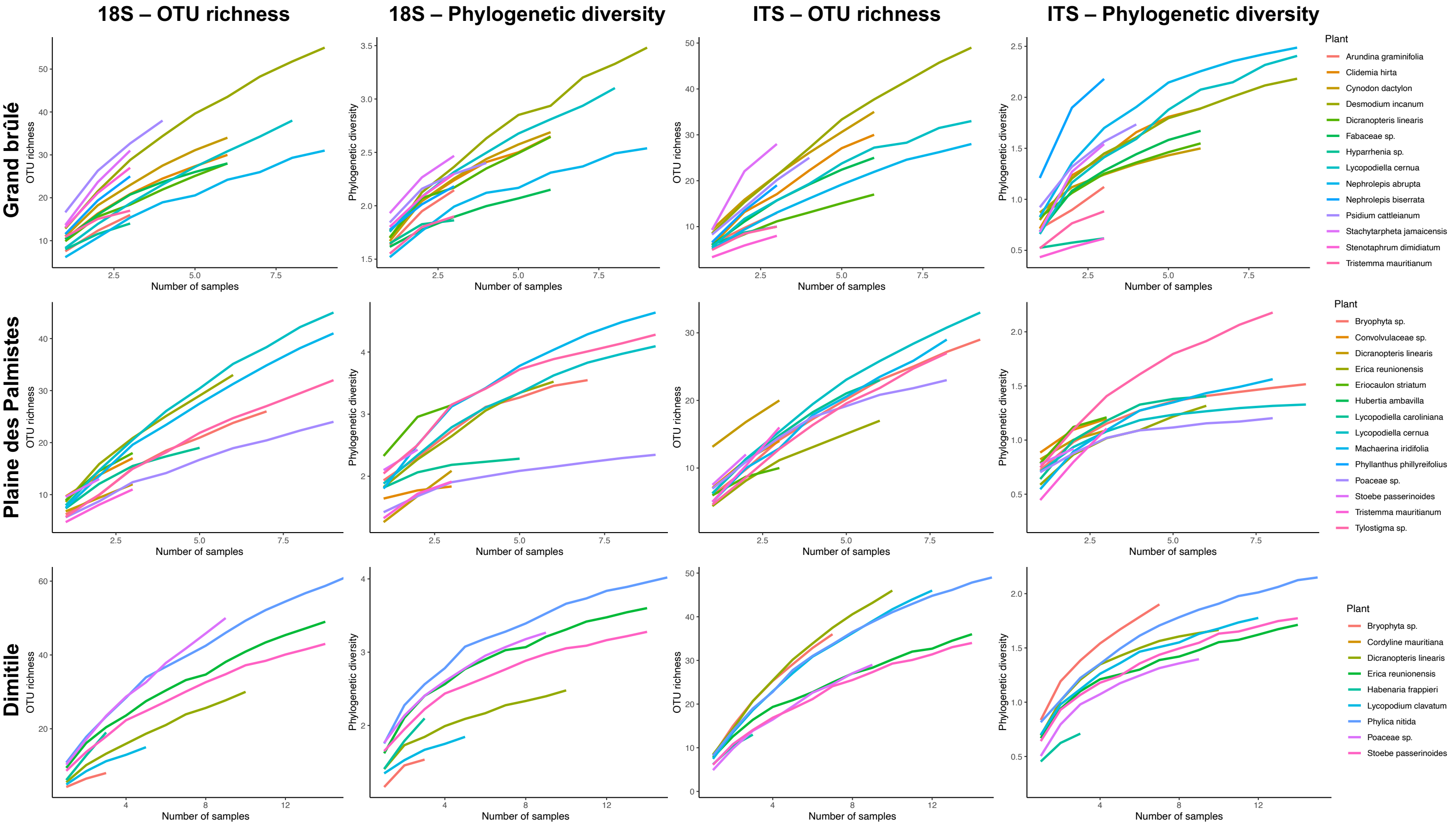

**Supplementary Figure 6: Principal coordinate analyses (PCoA) of the fungal mycobiomes associated with all plant roots across sampled communities (a-b) or with only the plant species simultaneously present in several sampled communities (c-d) show a strong clustering by sampled community (*i.e.* habitat).**

Each panel represents the projection of all the samples onto the two first axes of the PCoA performed on Bray-Curtis dissimilarities according to the 18S rRNA (a,c) or the ITS2 (b,d) markers. Each sample is colored according to its sampled community (Grand brûlé, Plaine-des-Palmistes, or Dimitile). The PCoA is then replicated between sampled from the same species found across several sampled communities (*Dicranopteris linearis, Erica reunionensis, Lycopodiella cernua,* and *Stoebe passerinoides*). Only results for the Bray-Curtis dissimilarities based on Swarm OTUs are represented (using UniFrac distances gave very similar results).

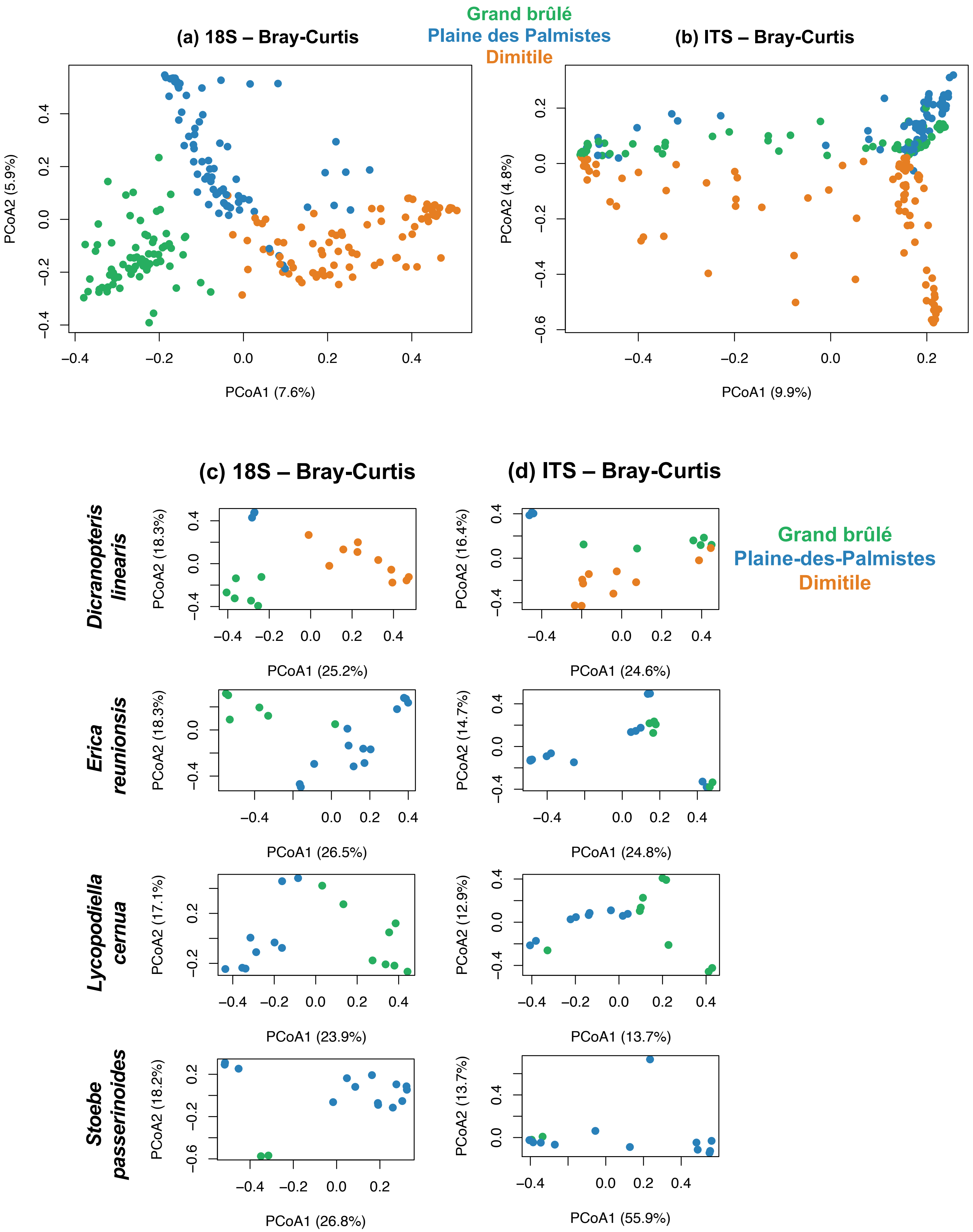

Grand brûlé
Plaine des Palmistes
Dimitile
(a) 18S – Bray-Curtis
(b) ITS – Bray-Curtis
PCoA2 (5.9%)
PCoA1 (7.6%)
PCoA2 (4.8%)
PCoA1 (9.9%)
(c) 18S – Bray-Curtis
(d) ITS – Bray-Curtis
Grand brûlé
Plaine-des-Palmistes
Dimitile
Dicranopteris linearis
PCoA2 (18.3%)
PCoA1 (25.2%)
PCoA2 (16.4%)
PCoA1 (24.6%)
Erica reunionsis
PCoA2 (18.3%)
PCoA1 (26.5%)
PCoA2 (14.7%)
PCoA1 (24.8%)
Lycopodiella cernua
PCoA2 (17.1%)
PCoA1 (23.9%)
PCoA2 (12.9%)
PCoA1 (13.7%)
Stoebe passerinoides
PCoA2 (18.2%)
PCoA1 (26.8%)
PCoA2 (13.7%)
PCoA1 (55.9%)

**Supplementary Figure 7: Root mycobiome compositions associated with root samples of *Lycopodiella cernua* significantly varied across the 2 sampled communities:**

For each *Lycopodiella cernua* root sample, the relative abundance of the fungi is indicated according to the 18S rRNA (top) or ITS2 (bottom) markers according to the sampled community (Grand brûlé or Plaine-des-Palmistes).

The class and the family of each fungus are represented in colors. Rare taxa (representing less than 0.1% of the data are represented in black).

Samples with less than 20 reads were discarded (empty space) and the number of reads is indicated at the top of each pie chart.

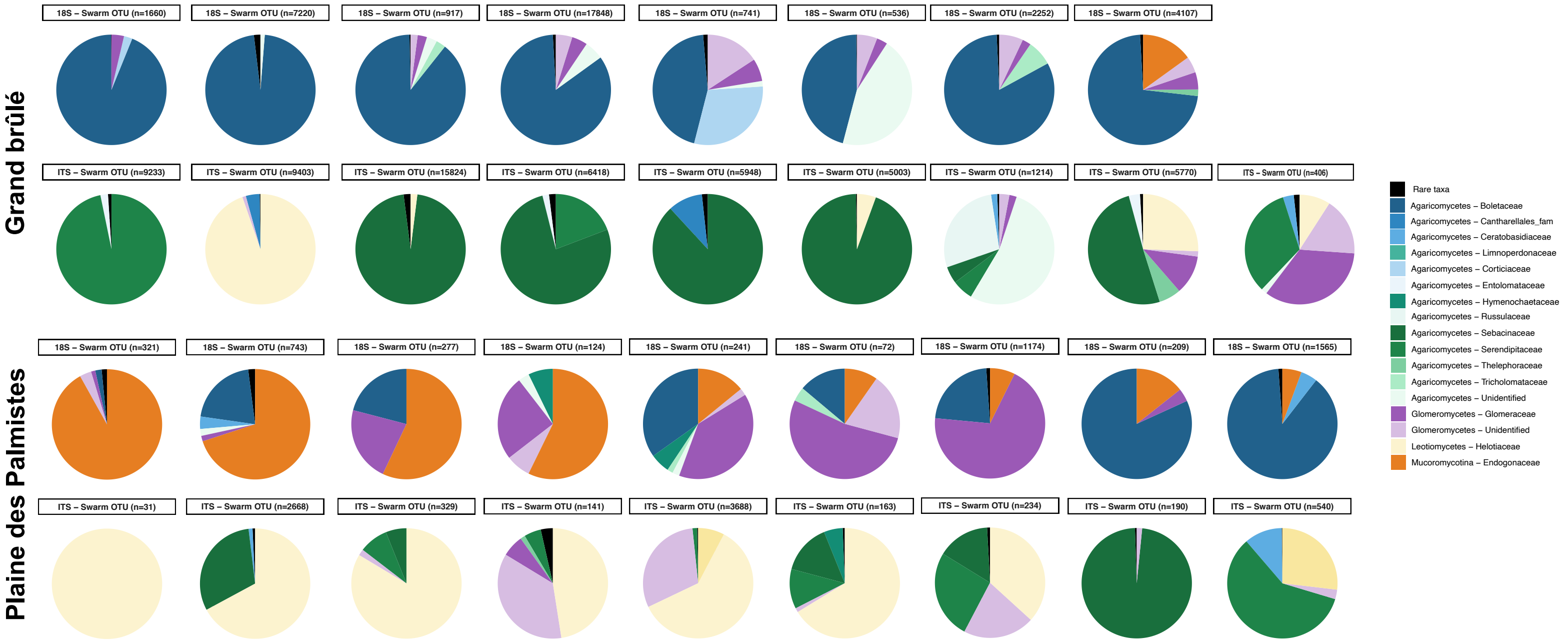

**Supplementary Figure 8: Principal coordinate analyses (PCoA) of the fungal mycobiomes associated with plant roots show an effect of both sampling plots (a) and plant taxonomic groups (b):**

Each panel represents the projection of the samples onto the two first axes of the PCoA performed on Bray-Curtis dissimilarities or on UniFrac distances according to the 18S rRNA (top panels) or the ITS2 (bottom panels) markers for each sampled community (Grand brûlé, Plaine-des-Palmistes, and Dimitile).

**(a)** Each root sample is colored according to its sampling plot (R1 in blue, R2 in red, or R3 in purple).

**(b)** Each root sample is colored according to the plant taxonomic group (bryophytes, lycopods, ferns, monocots (excluding *Orchidaceae*), eudicots (excluding *Ericaceae*), *Orchidaceae*, or *Ericaceae*).

#### (a) Sampling plot

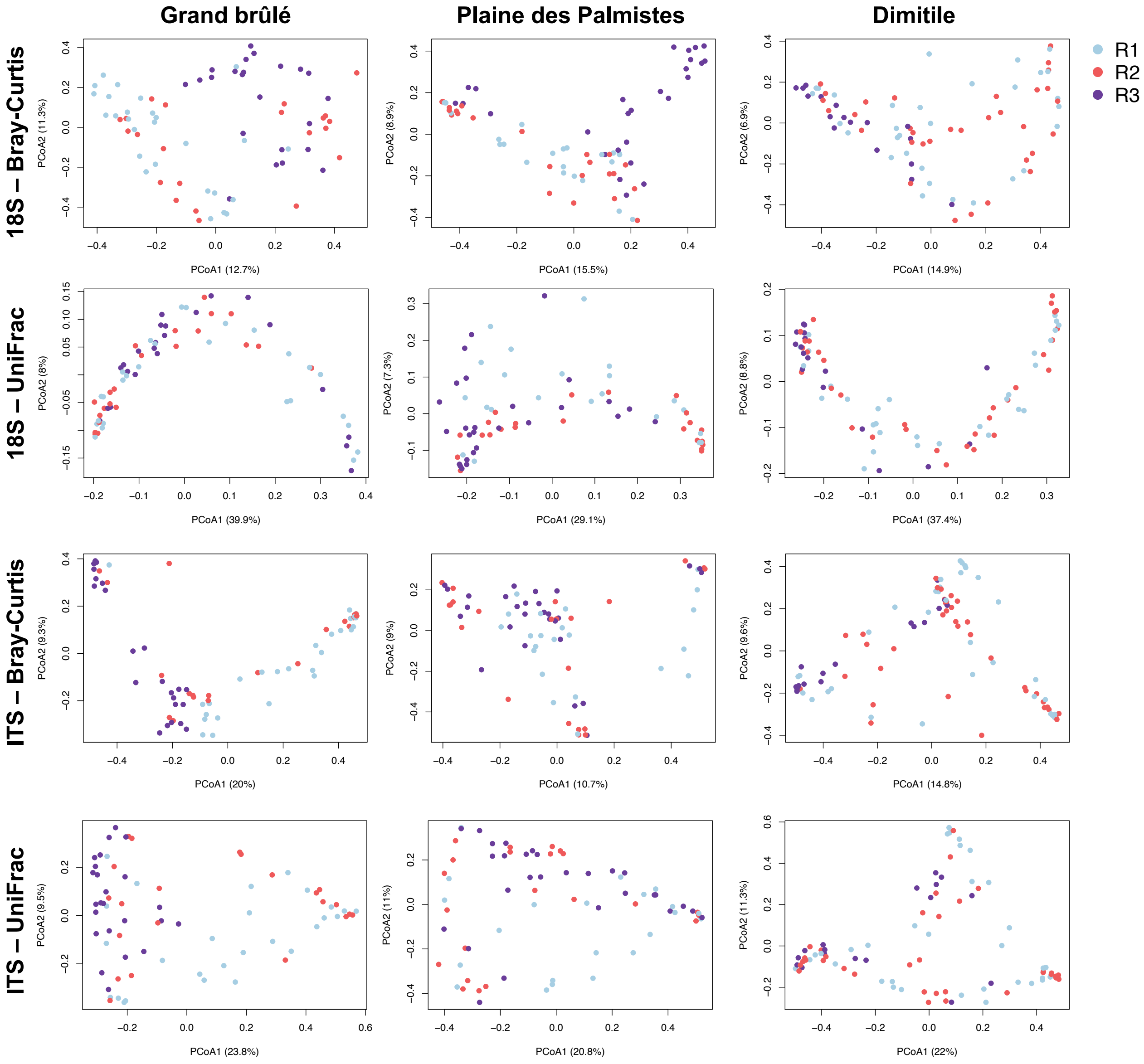

#### (b) Plant taxonomic group

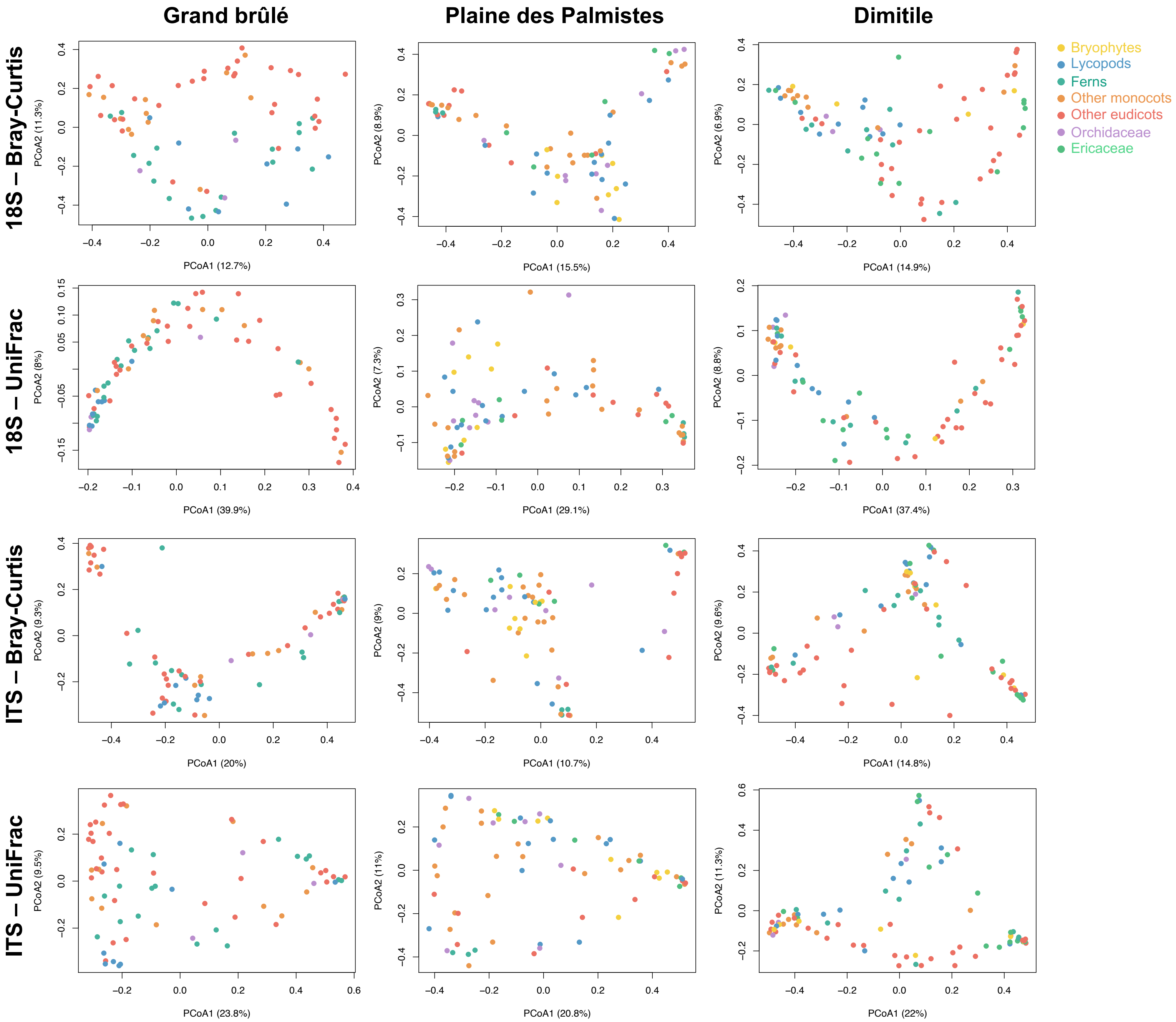

**Supplementary Figure 9: Plant-fungus network representations vary according to the fungal lineages:**

Each panel represents the plant-fungus network in a given sampled community and for a given clade of fungi: *Glomeromycotina* (I), *Mucoromycotina* (II), *Sebacinales* (III), *Helotiales* (IV), and *Cantharellales* (V). In each panel, plant species are in rows and fungal OTUs are in columns, and the elements of the matrix are colored according to the abundances of each plant-fungus interaction (*i.e.* relative number of reads): dark greens represent abundant interactions, whereas light greens represent rare interactions. Each network corresponds to one sampled community: Grand brûlé (a), Plaine-des-Palmistes (b), or Dimitile (c).

*Glomeromycotina* and *Mucoromycotina* networks were characterized using the 18S rRNA marker, whereas the *Sebacinales*, *Helotiales*, and *Cantharellales* networks were characterized using the ITS2 marker.

(I) *Glomeromycotina*

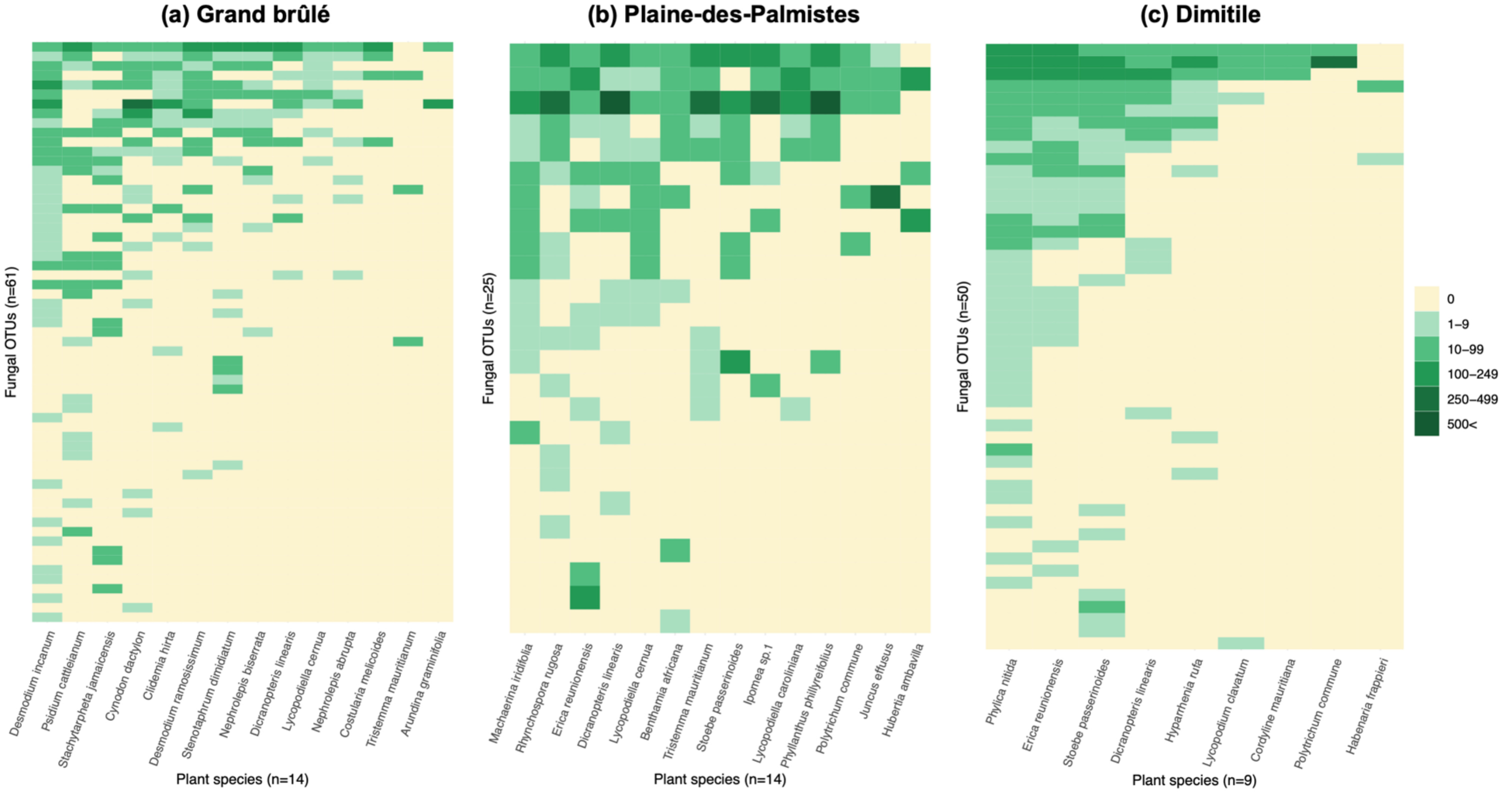

(II) *Mucoromycotina*

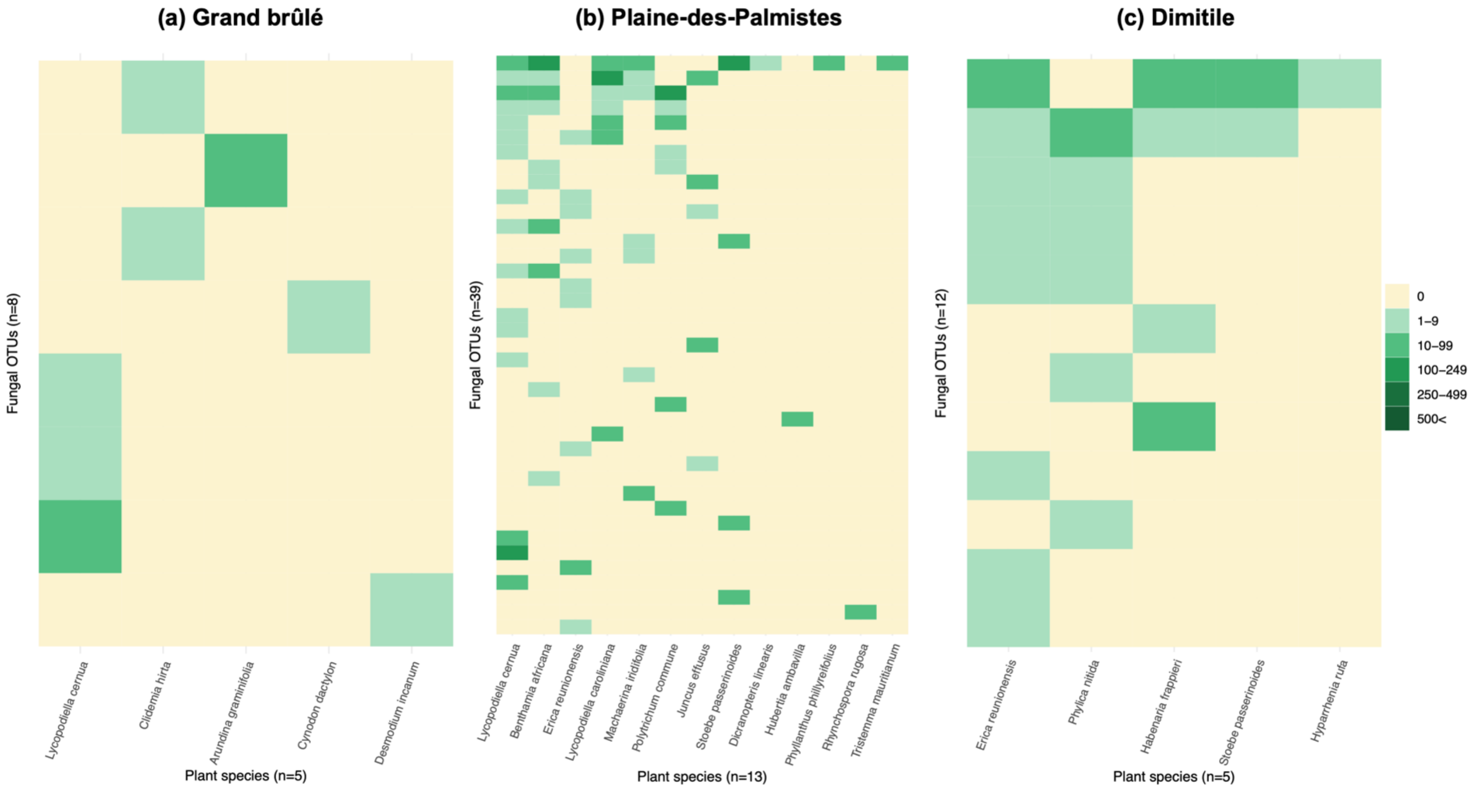


(III) *Sebacinales*

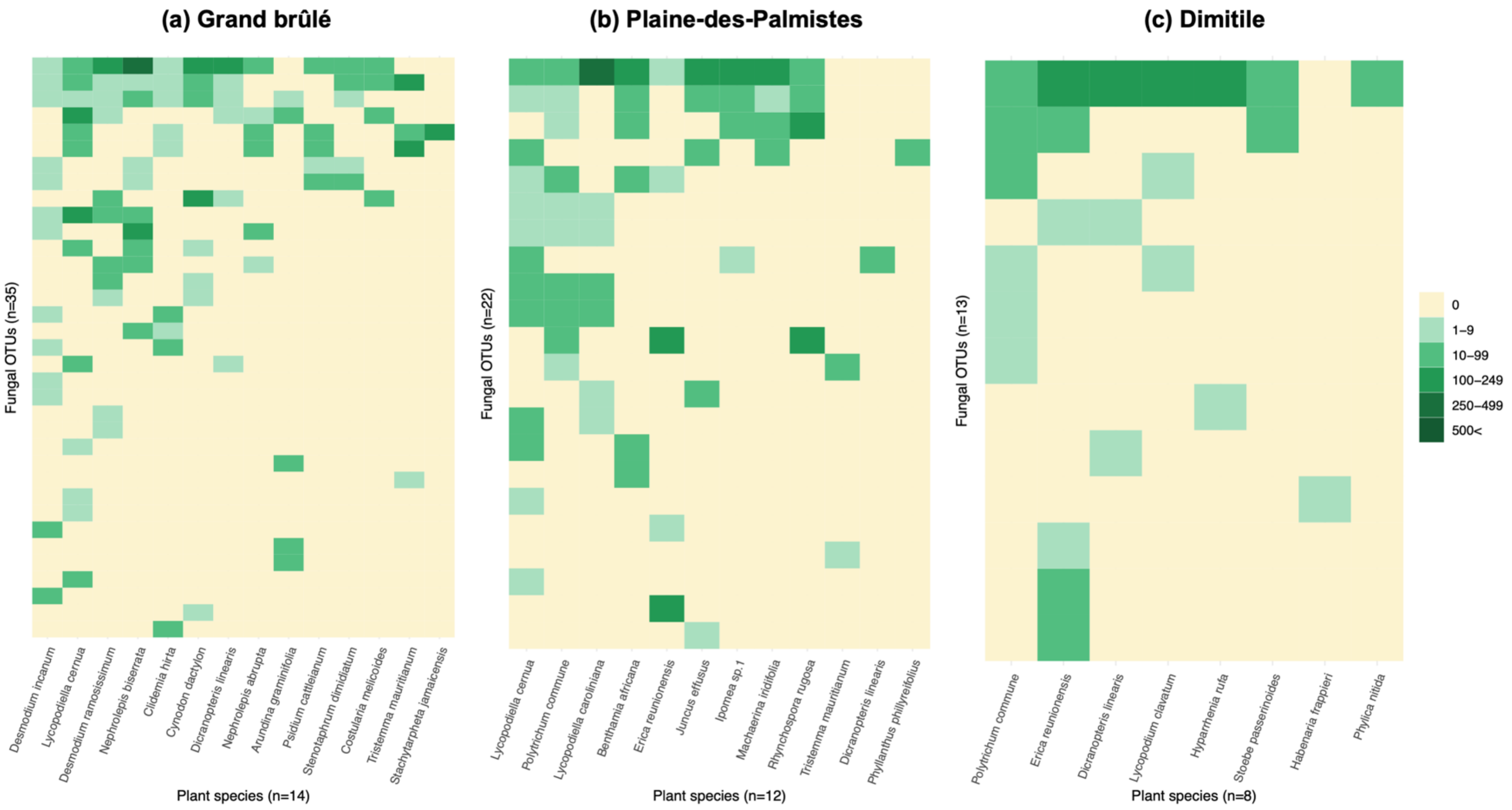

(IV) *Helotiales*

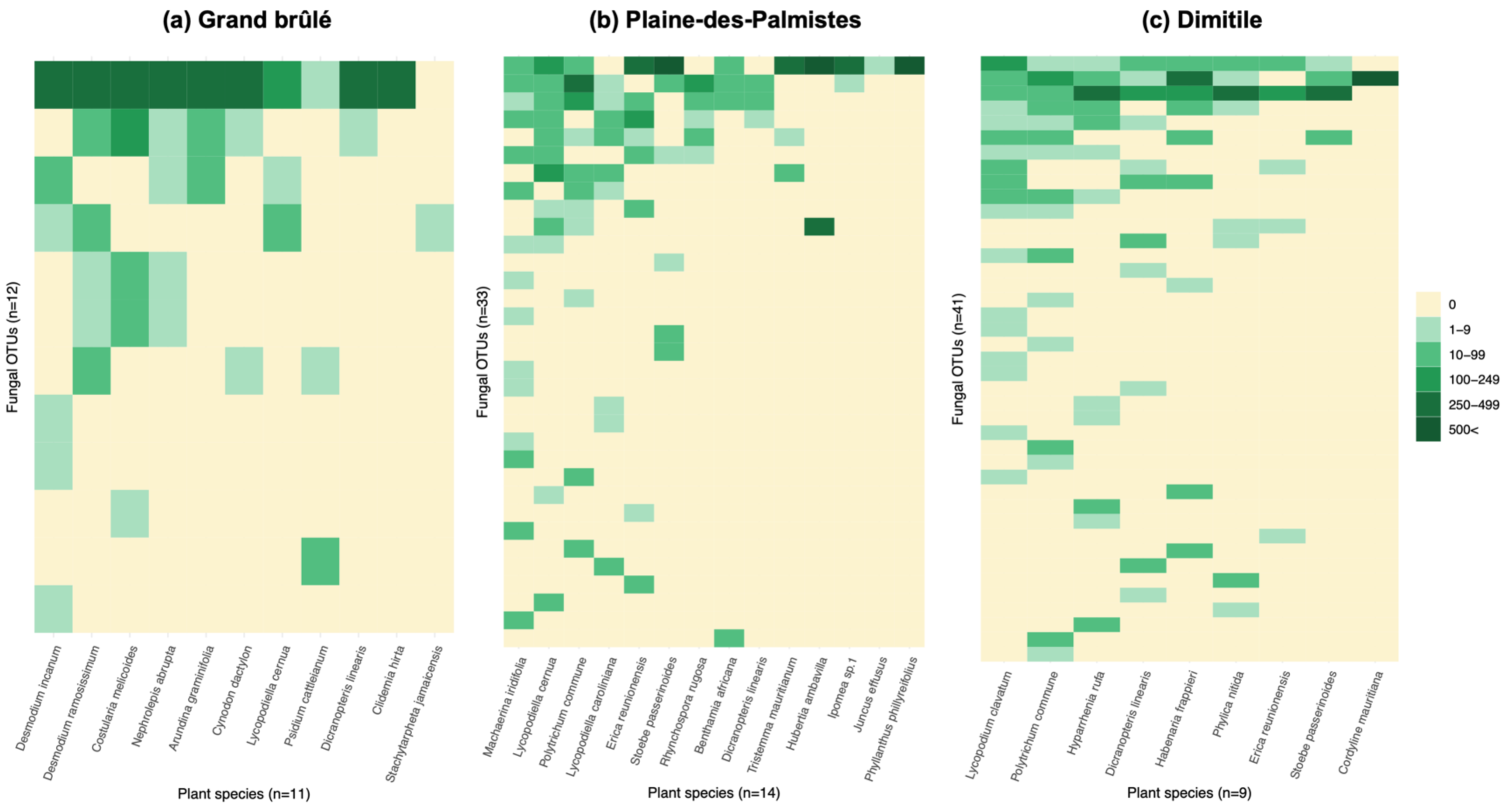


(V) *Cantharellales*

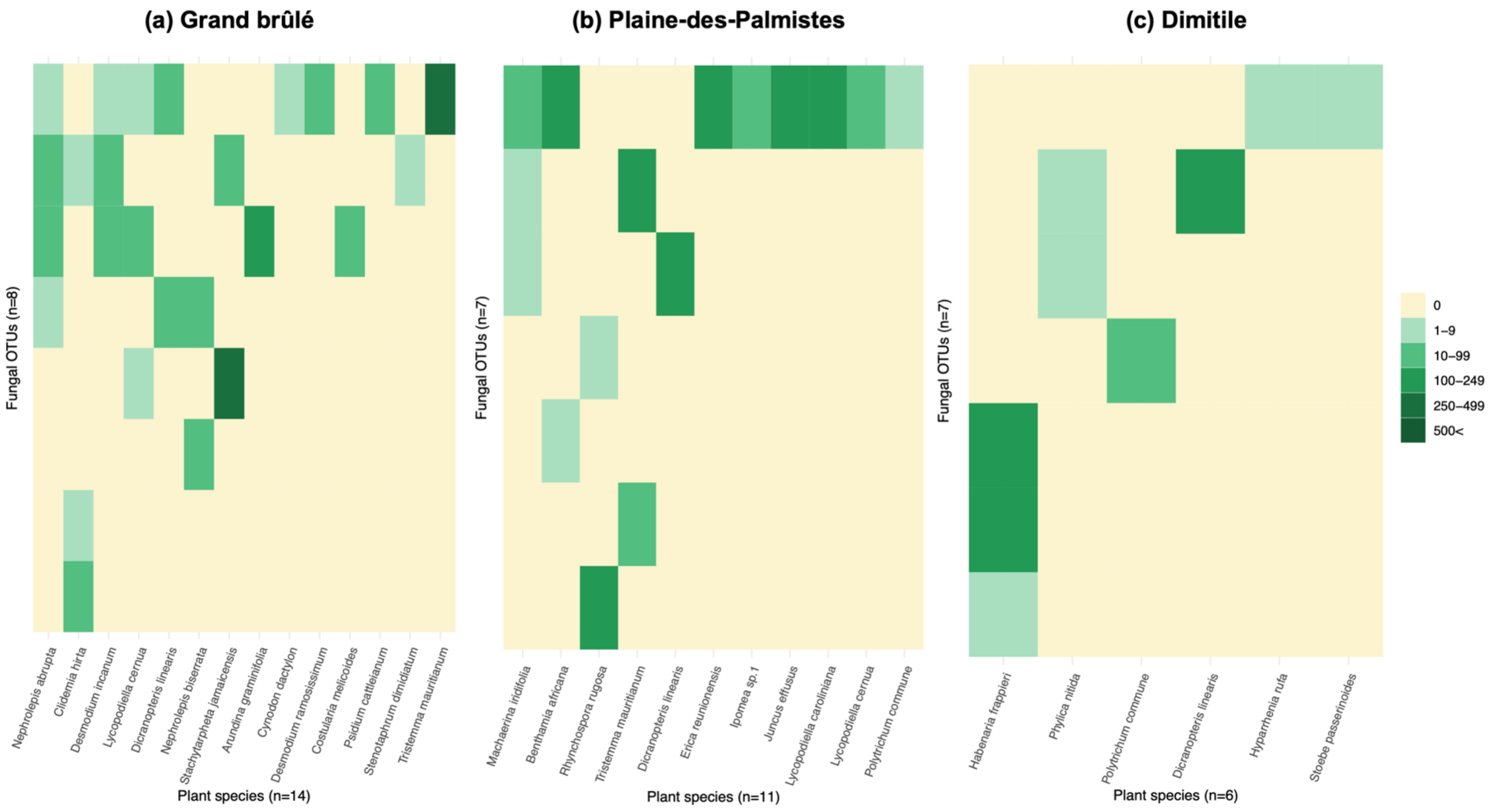

**Supplementary Figure 10: Plant-fungus network structures are robust to the exclusion of rare plant species.**

Species-level network representation in each sampled community (Plaine-des-Palmistes or Dimitile) for the different fungal groups (*Glomeromycotina, Helotiales, Sebacinales, Mucoromycotina*, or *Cantharellales*) when excluding rare plant species, that were only sampled once or twice. Because all plant species in Grand Brûlé were sampled at least 3 times, the results for this community were not influenced by the exclusion of rare species (the networks in Grand Brûlé are therefore not represented here). Colored round nodes represent plant species (colors indicate the main plant taxonomic groups) and grey squared nodes correspond to fungal OTUs. Grey links represent plant-fungus interactions and their widths are proportional to interaction abundances. The position of the nodes reflects the similarity in species interactions using the Fruchterman-Reingold layout algorithm [99] from the *igraph* R-package. Fungal lineages (in rows) are ordered according to their network structures: networks that tend to be nested are at the top, whereas networks that tend to be modular are at the bottom. For each network, the total read abundances (L), the connectance (C), and the ratio of interactions within modules (Q) are indicated. Q is computed from the most modular structure according to Beckett's algorithm for abundance networks, Q close to 0 indicates that most interactions are between modules (*i.e.* low modularity), while Q close to 1 indicates that most interactions are within modules (*i.e.* high modularity).

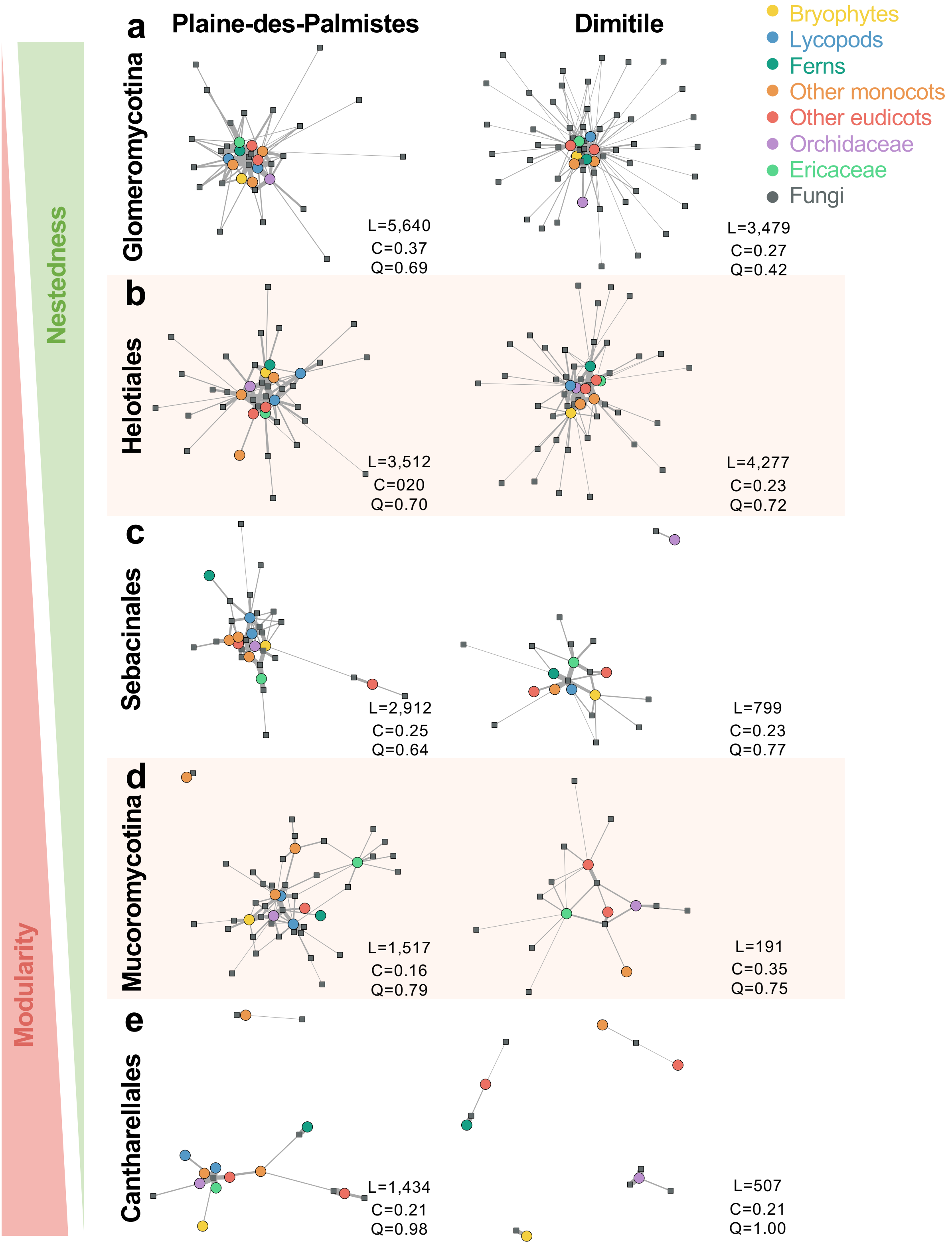

Plaine-des-Palmistes
Dimitile
Bryophytes
Lycopods
Ferns
Other monocots
Other eudicots
Orchidaceae
Ericaceae
Fungi
Nestedness
Modularity
a
Glomeromycotina
L=5,640
C=0.37
Q=0.69
L=3,479
C=0.27
Q=0.42
b
Helotiales
L=3,512
C=020
Q=0.70
L=4,277
C=0.23
Q=0.72
c
Sebacinales
L=2,912
C=0.25
Q=0.64
L=799
C=0.23
Q=0.77
d
Mucoromycotina
L=1,517
C=0.16
Q=0.79
L=191
C=0.35
Q=0.75
e
Cantharellales
L=1,434
C=0.21
Q=0.98
L=507
C=0.21
Q=1.00

**Supplementary Figure 11: Interaction specializations ($H_2'$) of the plant-fungus networks significantly vary according to the fungal lineages**

For each plant-fungus network (*Glomeromycotina, Mucoromycotina, Sebacinales, Helotiales*, or *Cantharellales*) in each sampled community (Grand brûlé (A), Plaine-des-Palmistes (B), or Dimitile (C)), a colored dot indicates the network-level interaction specialization ($H_2'$) obtained using abundance networks of Swarm OTUs.

The significance of the $H_2'$ values was evaluated using null models maintaining marginal sums (a) or shuffle-sample null models (b), and the distributions of the $H_2'$ values of the null models are represented with the black boxplots. Colored horizontal lines indicate the 95% confidence interval.

Boxplots present the median surrounded by the first and third quartiles, and whiskers extend to the extreme.

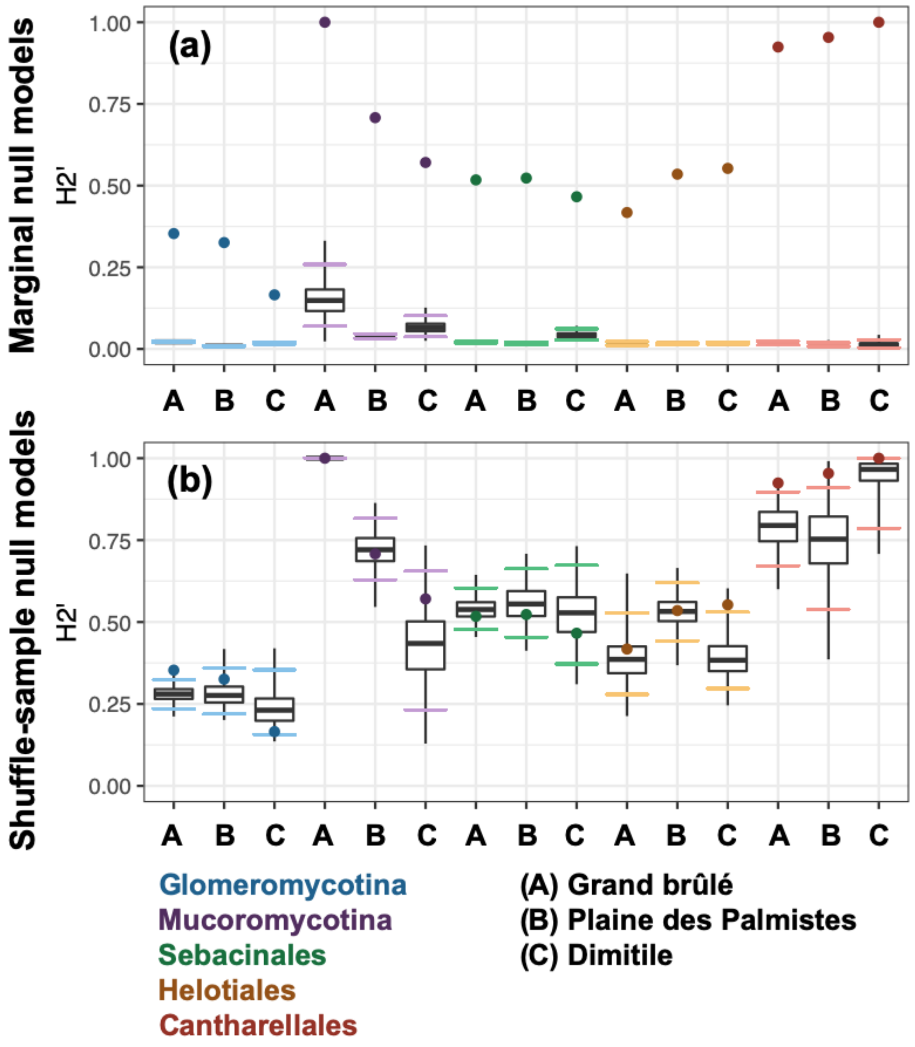

**Supplementary Figure 12: Patterns of plant-fungus specialization are robust to the exclusion of rare plant species.**

We replicated analyses of plant-fungus specialization while excluding rare plant species, that were only sampled once or twice. Because all plant species in Grand Brûlé were sampled at least 3 times, the results for this community were not influenced by the exclusion of rare species.

**(a) Interaction specializations ($H_2'$) are lower in plant-*Glomeromycotina* networks than in other plant-fungus networks.** For each plant-fungus network (with *Glomeromycotina, Mucoromycotina, Sebacinales, Helotiales*, or *Cantharellales*) in each sampled community (Grand brûlé (A), Plaine-des-Palmistes (B), or Dimitile (C)), a colored dot indicates the network-level interaction specialization ($H_2'$).

**(b) Motif frequencies significantly differ between the plant-fungus networks**. Principal coordinate analyses (PCoA) of the bipartite motif frequencies (the "building blocks" of the network containing from 2 to 5 species) of each plant-fungus network (*Glomeromycotina, Mucoromycotina, Sebacinales, Helotiales*, or *Cantharellales*) in each sampled community (Grand brûlé (A), Plaine-des-Palmistes (B), or Dimitile (C)). The colored triangle areas represent the proximity within the sampled communities for the different groups of fungi.

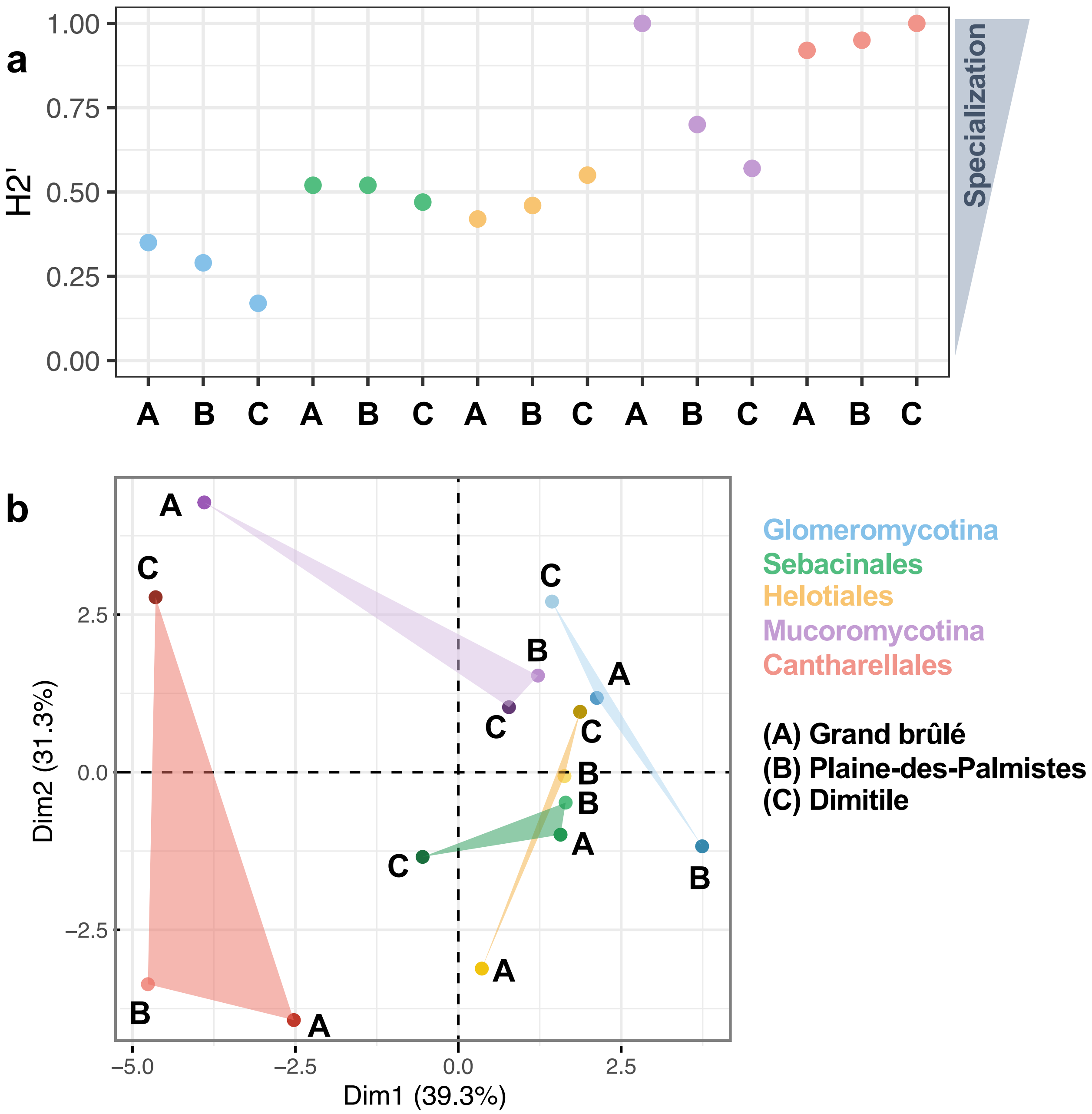

a
H2'
1.00
0.75
0.50
0.25
0.00
A B C A B C A B C A B C A B C
Specialization
b
Dim2 (31.3%)
Dim1 (39.3%)
2.5
0.0
−2.5
−5.0
−2.5
0.0
2.5
Glomeromycotina
Sebacinales
Helotiales
Mucoromycotina
Cantharellales
(A) Grand brûlé
(B) Plaine-des-Palmistes
(C) Dimitile

**Supplementary Figure 13: Plant species are specialized towards their fungi**

For each plant-fungus network (*Glomeromycotina* (I), *Mucoromycotina* (II), *Sebacinales* (III), *Helotiales* (IV), or *Cantharellales* (V)) in each sampled community (Grand brûlé, Plaine-des-Palmistes, or Dimitile), plant specialization was measured for each plant species using normalized degree (top panels) and d' index (bottom panels) and represented using red dots. Results were obtained for the abundance networks of Swarm OTUs.

The significance of the specialization values was evaluated using null models maintaining marginal sums or shuffle-sample null models, and the distribution of the specialization values of the null models are represented with the black boxplots. Orange horizontal lines indicate the 95% confidence interval.

Boxplots present the median surrounded by the first and third quartiles, and whiskers extend to the extreme values.

(I) *Glomeromycotina* – marginal null models

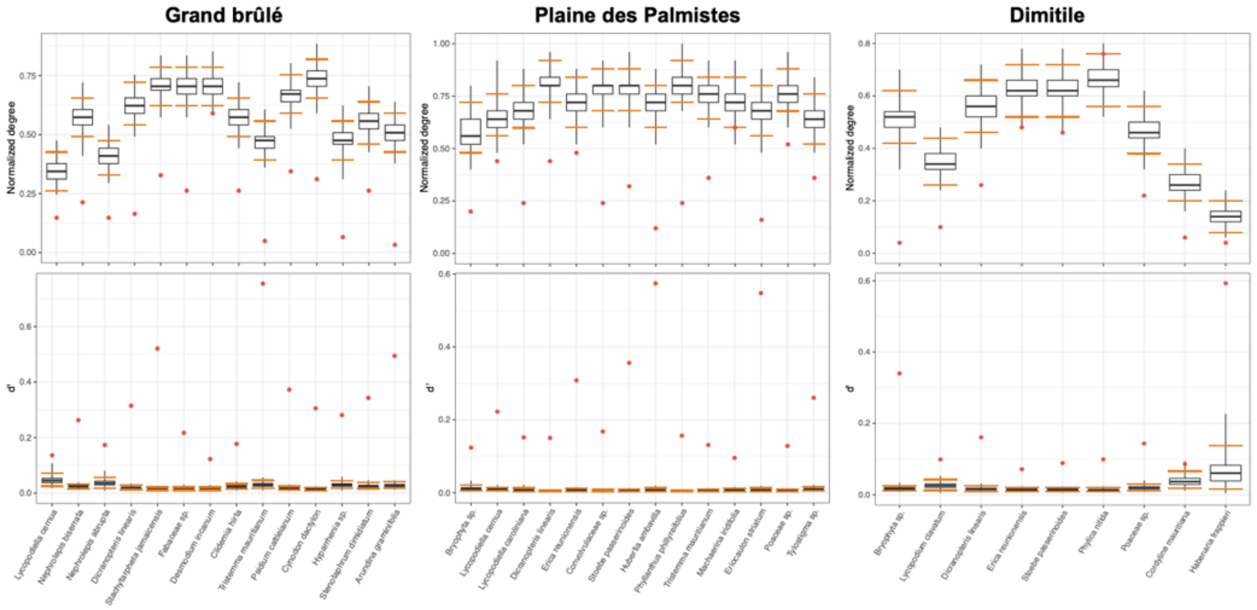


(I) *Glomeromycotina* – shuffle-sample null models

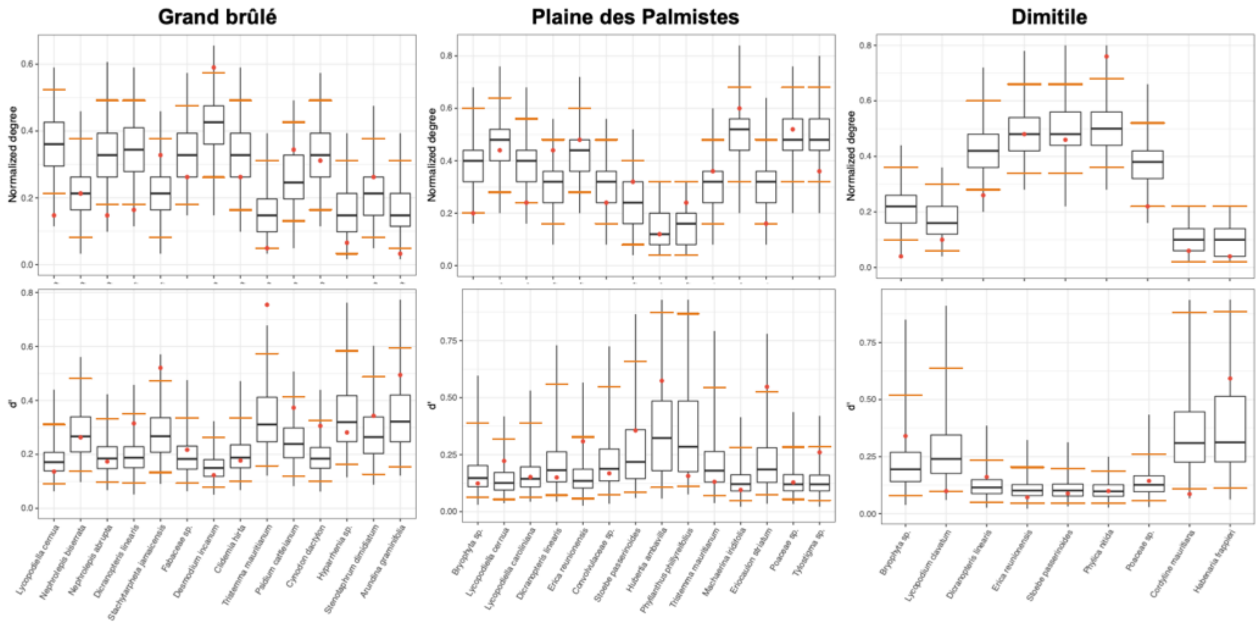

(II) *Mucoromycotina* – marginal null models

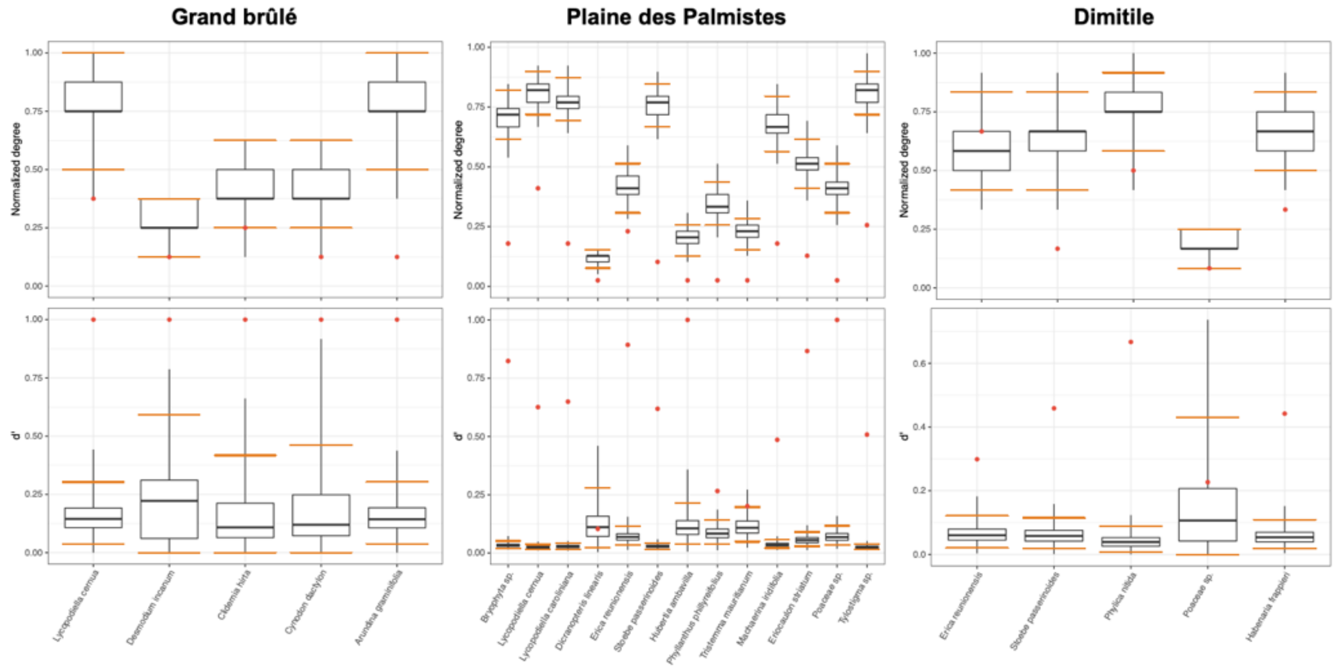


(II) *Mucoromycotina* – shuffle-sample null models

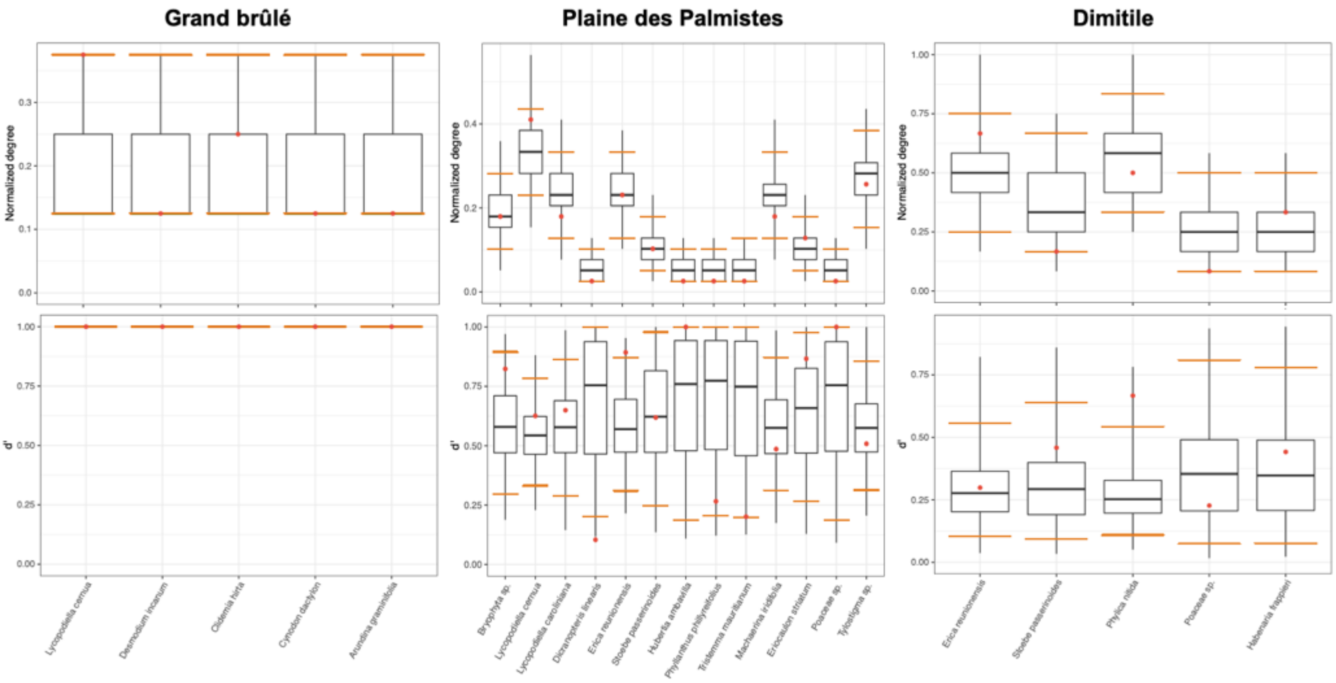

(III) *Sebacinales* – marginal null models

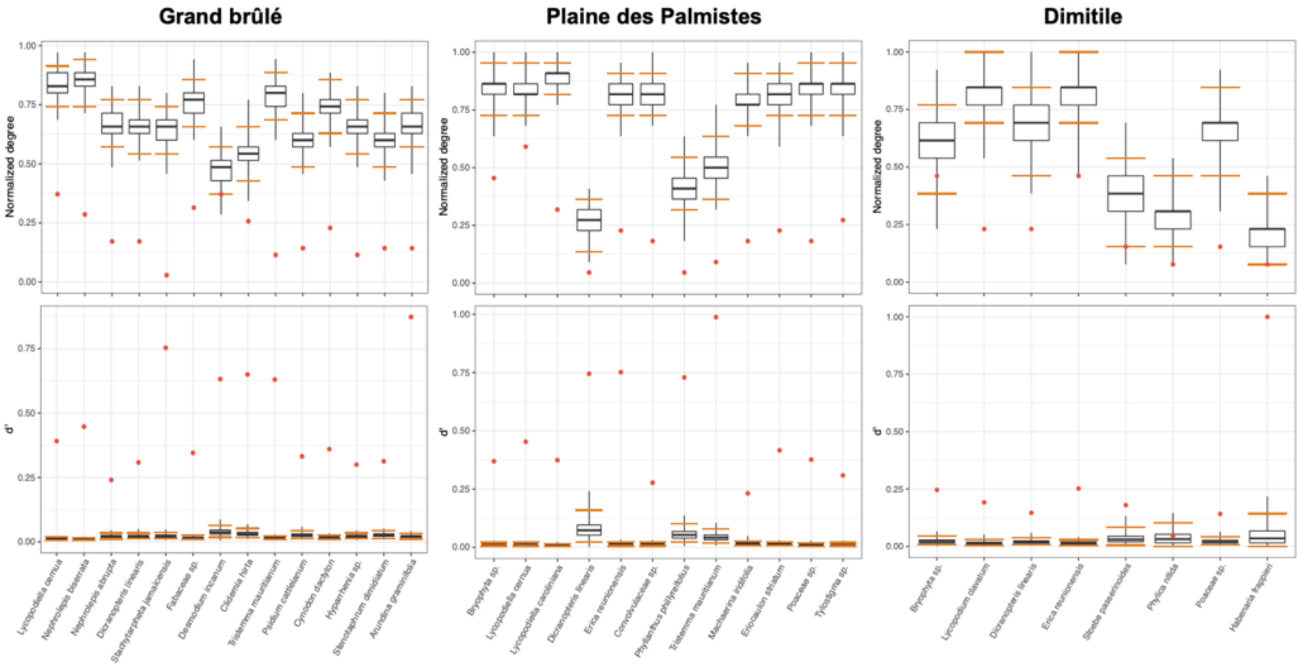


(III) *Sebacinales* – shuffle-sample null models

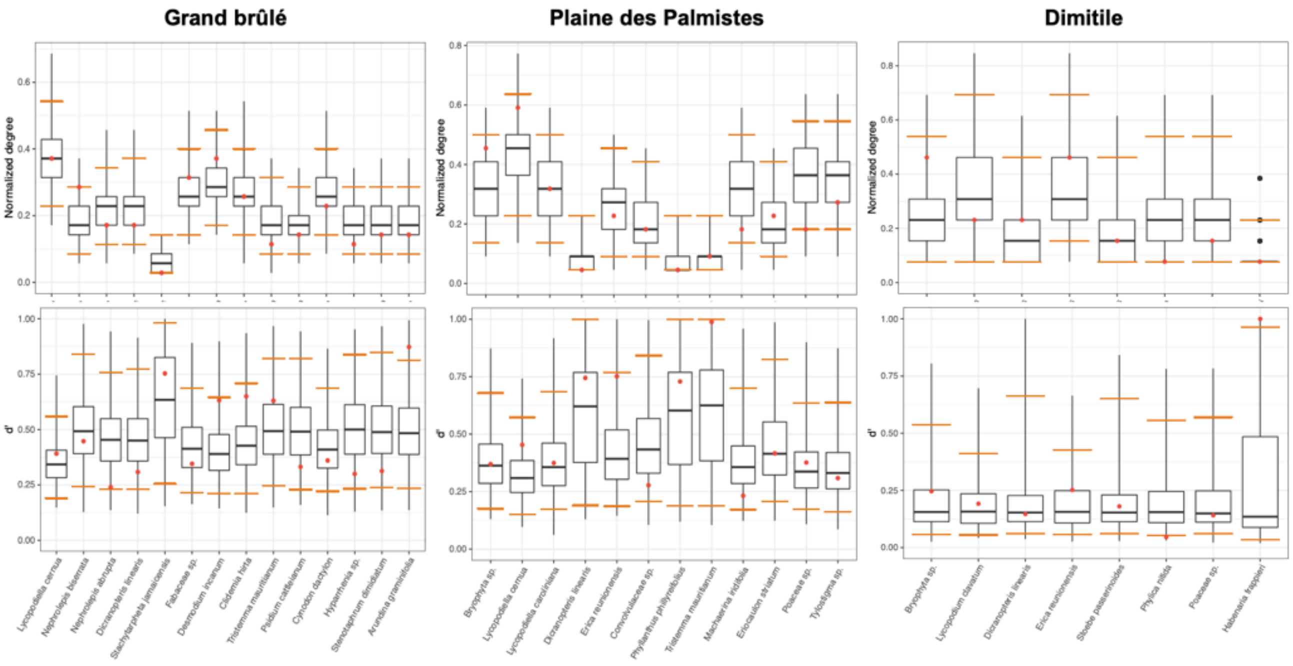

(IV) *Helotiales* – marginal null models

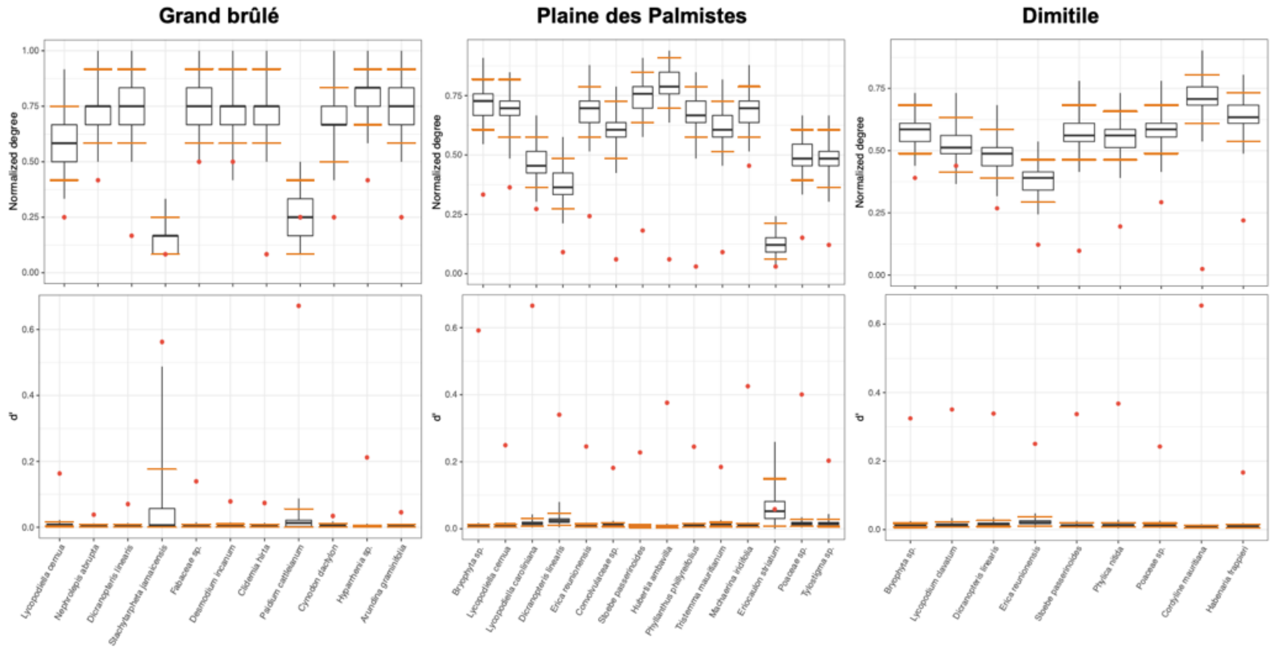


(IV) *Helotiales* – shuffle-sample null models

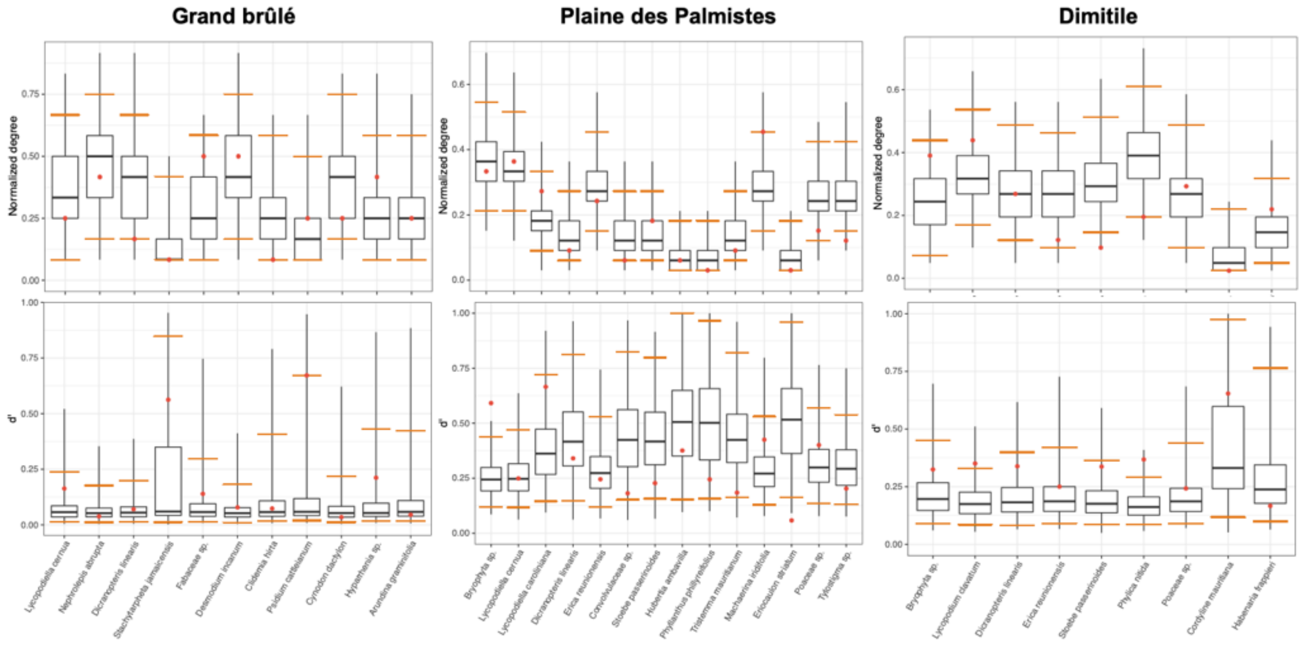

(IV) *Cantharellales* – marginal null models

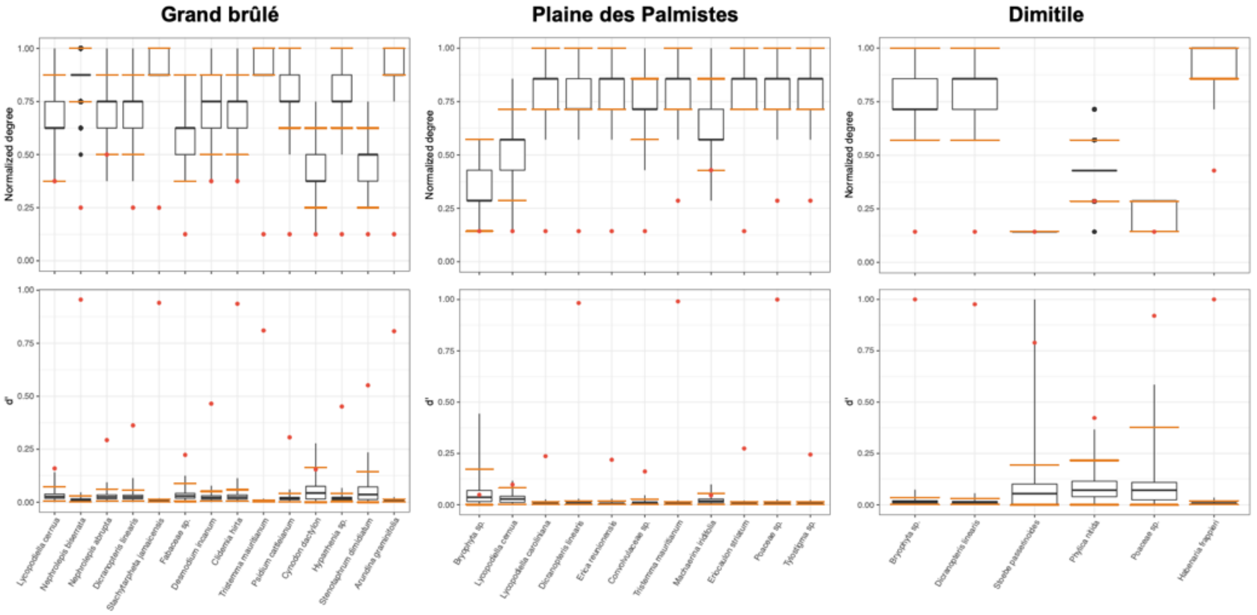


(IV) *Cantharellales* – shuffle-sample null models

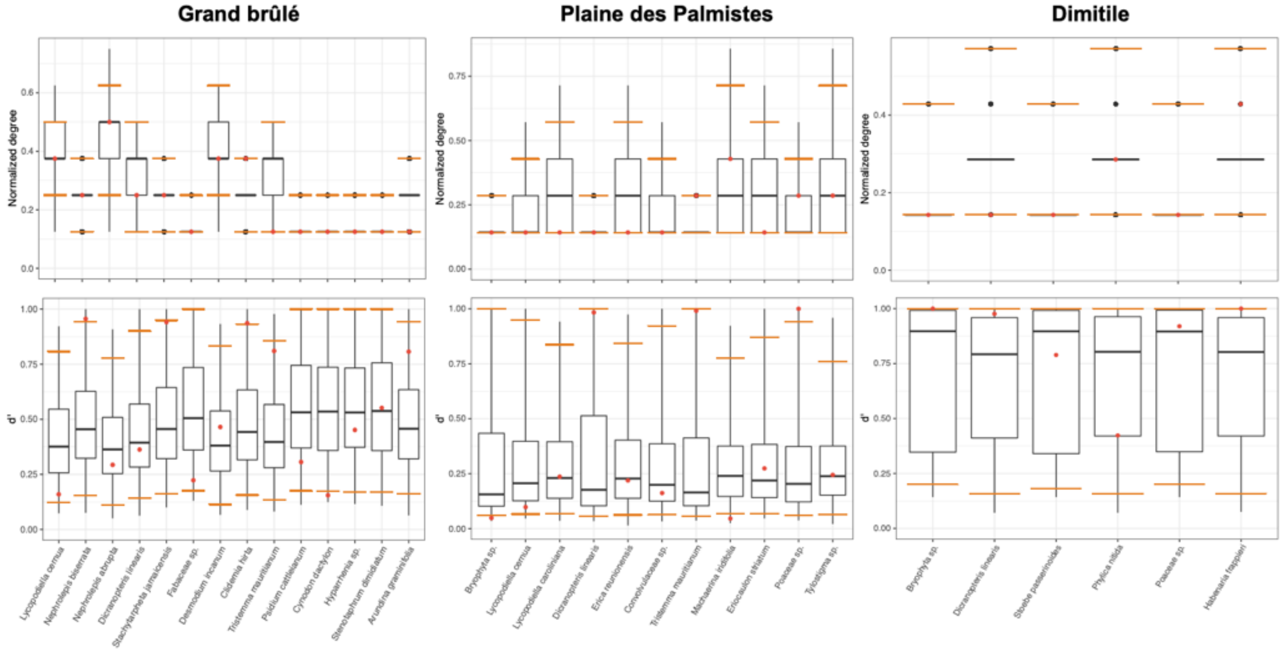

**Supplementary Figure 14: Some motif frequencies in the plant-fungus networks tend to vary between the different fungal lineages:**

For each plant-fungus binary network (*Glomeromycotina, Mucoromycotina, Sebacinales, Helotiales*, or *Cantharellales*) in each sampled community (Grand brûlé, Plaine-des-Palmistes, or Dimitile), motif frequencies were computed (for all motifs containing from 2 to 5 species) (a). Dots are colored according to the type of plant-fungus networks. Motifs are numbered from 1 to 17 and panel (b) presents the different motifs (adapted from Figure 1 of (Simmons *et al.*, 2019); the colors of the links are not informative here).

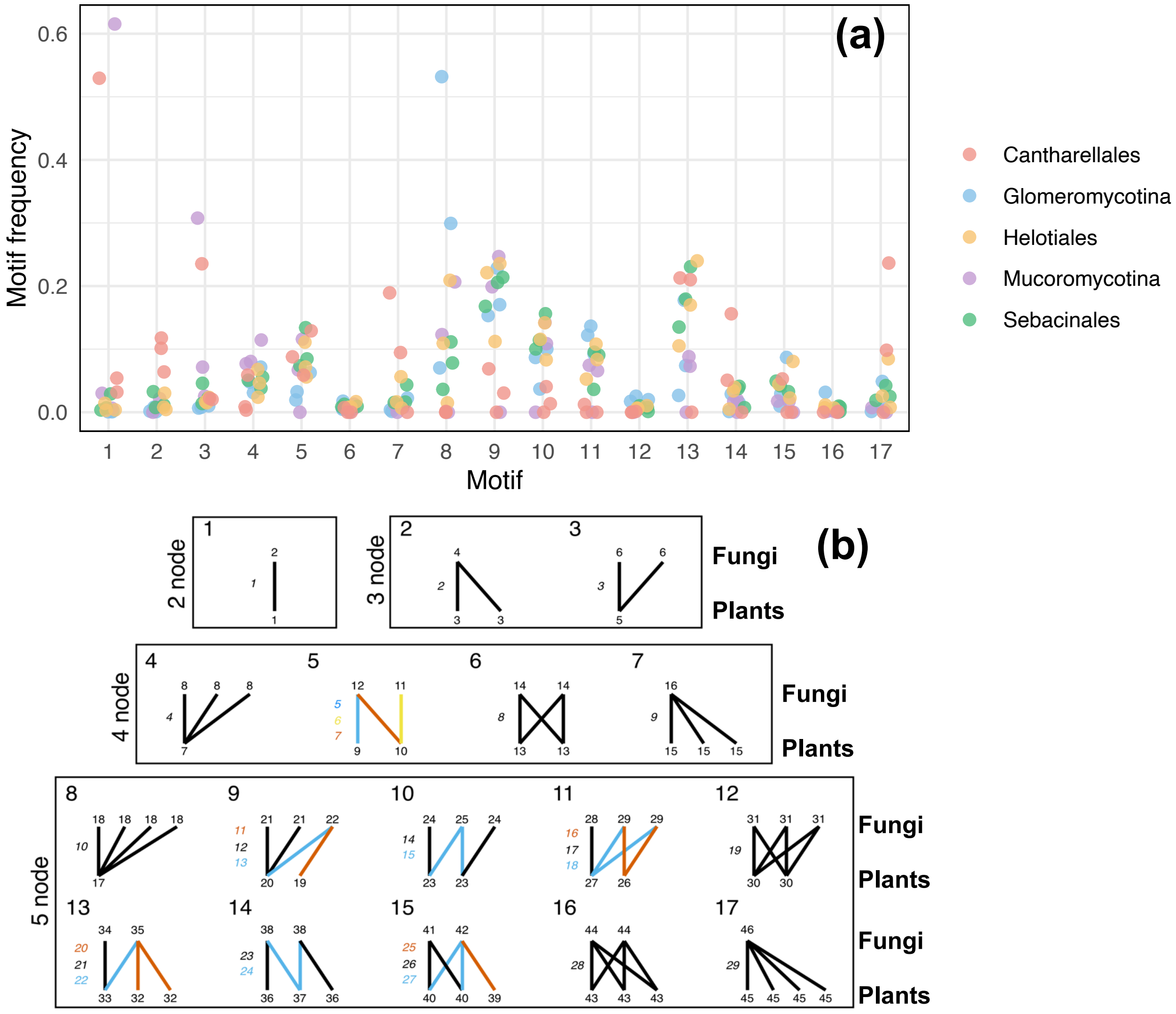

**Supplementary Figure 15: The level of specialization of plant-*Glomeromycotina* interaction remains on average low even when characterized using the ITS2 marker:** (a) Network representation of the plant-*Glomeromycotina* network in each sampled community (Grand brûlé, Plaine-des-Palmistes, or Dimitile). Colored round nodes represent plant species (colors separate the main plant functional groups) and grey squared nodes correspond to fungal OTUs. Grey links represent plant-fungus interactions. Network-level interaction specializations ($H_2'$) are indicated for each sampled community.

(b) For each plant-*Glomeromycotina* network in each sampled community (Grand brûlé, Plaine-des-Palmistes, or Dimitile), motif frequencies were computed (for all motifs containing from 2 to 5 species) and plotted in the following graph for Swarm OTUs. Motifs are numbered from 1 to 17 (adapted from Figure 1 of Simmons *et al.*, 2019).

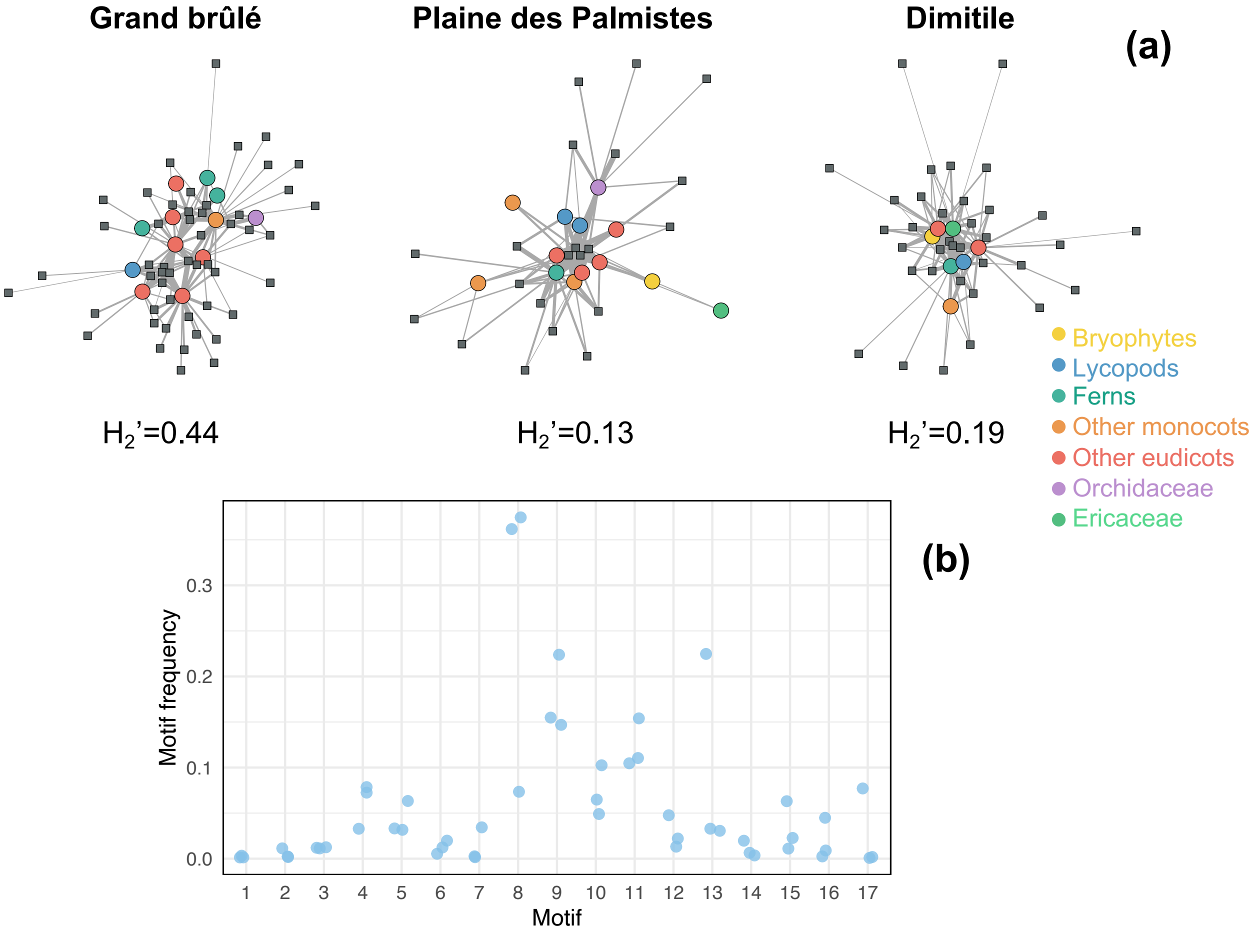

**Supplementary Figure 16: The level of specialization of plant-*Sebacinales* remains on average high when characterized using the 18S rRNA marker:**

(a) Network representation of the plant-*Sebacinales* network in each sampled community (Grand brûlé, Plaine-des-Palmistes, or Dimitile). Colored round nodes represent plant species (colors separate the main plant functional groups) and grey squared nodes correspond to fungal OTUs. Grey links represent plant-fungus interactions. Network-level interaction specializations ($H_2'$) are indicated for each sampled community.

(b) For each plant-*Sebacinales* network in each sampled community (Grand brûlé, Plaine-des-Palmistes, or Dimitile), motif frequencies were computed (for all motifs containing from 2 to 5 species) and plotted in the following graph for Swarm OTUs. Motifs are numbered from 1 to 17 (adapted from Figure 1 of Simmons *et al.*, 2019).

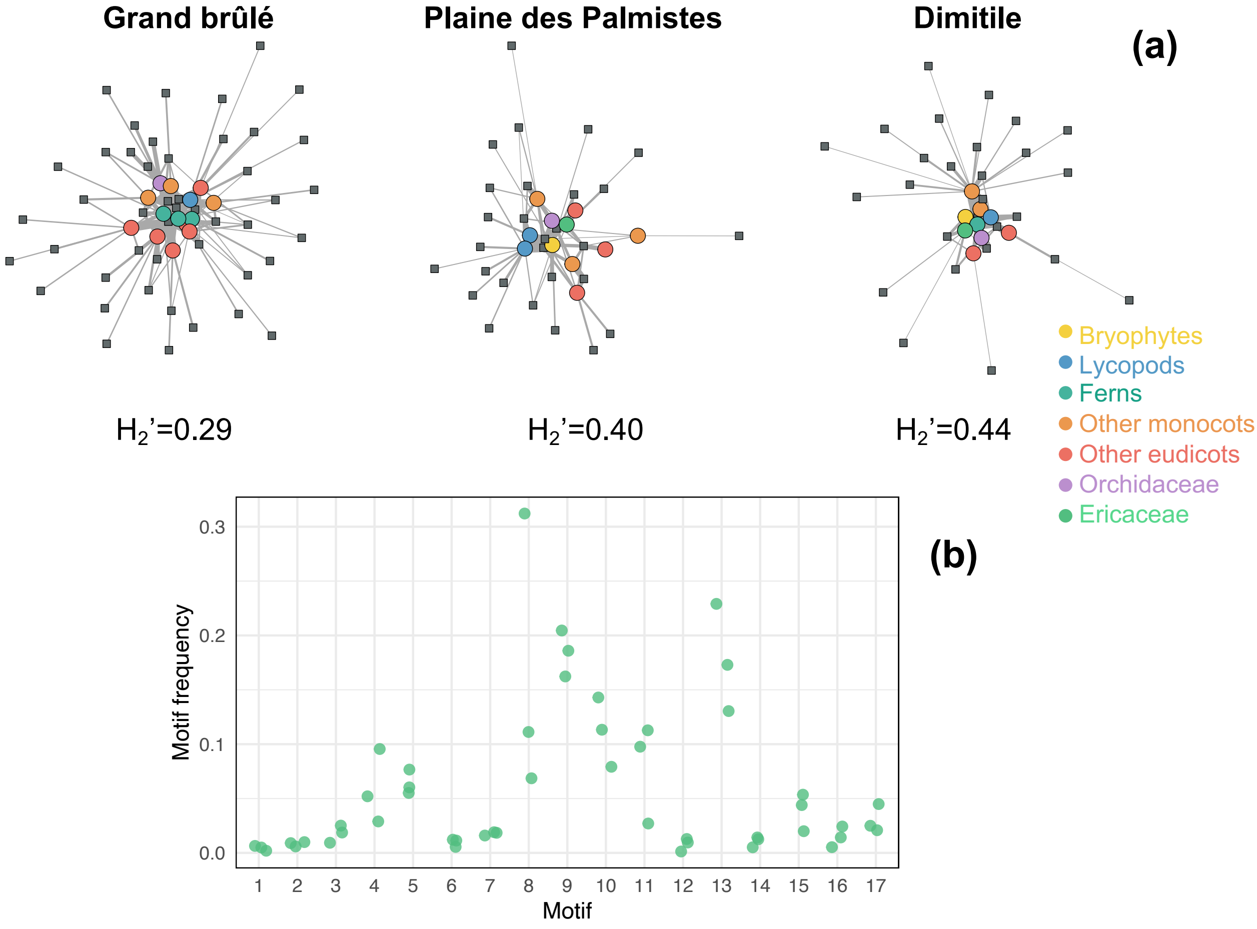

**Supplementary Figure 17: Motif frequencies for the different lycopod species indicate that lycopods are well connected to other plant species but that some lycopod-specific fungi are also frequent**

Motif position frequencies of each lycopod species in the different plant-fungus networks (*Glomeromycotina, Mucoromycotina, Sebacinales, Helotiales*, and *Cantharellales*) in each sampled community (Grand brûlé, Plaine-des-Palmistes, or Dimitile) are represented using orange bars, whereas the motif position frequencies for all the other plant species in the network are represented in grey. The different motifs are represented at the bottom (adapted from Figure 1 of (Simmons *et al.*, 2019)), where plant species occupy the "lower level" (motifs 1, 3, 5…). The colors of the link are not informative here.

The photos at the top represent the main lycopod species in each community.

***Lycopodiella cernua***
**in Grand brûlé**

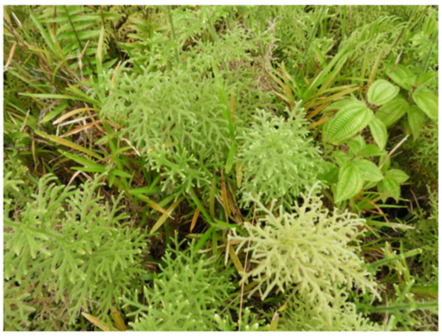

***Lycopodiella cernua* and**
***Lycopodiella caroliniana***
**in Plaine des Palmistes**

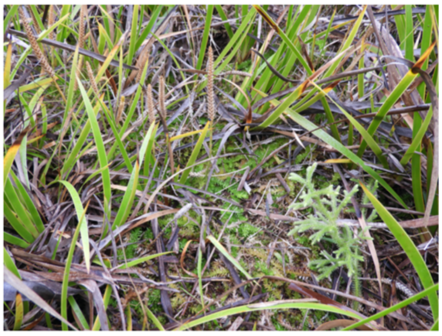

***Lycopodium clavatum***
**in Dimitile**

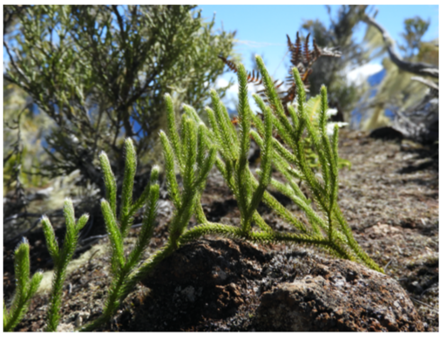

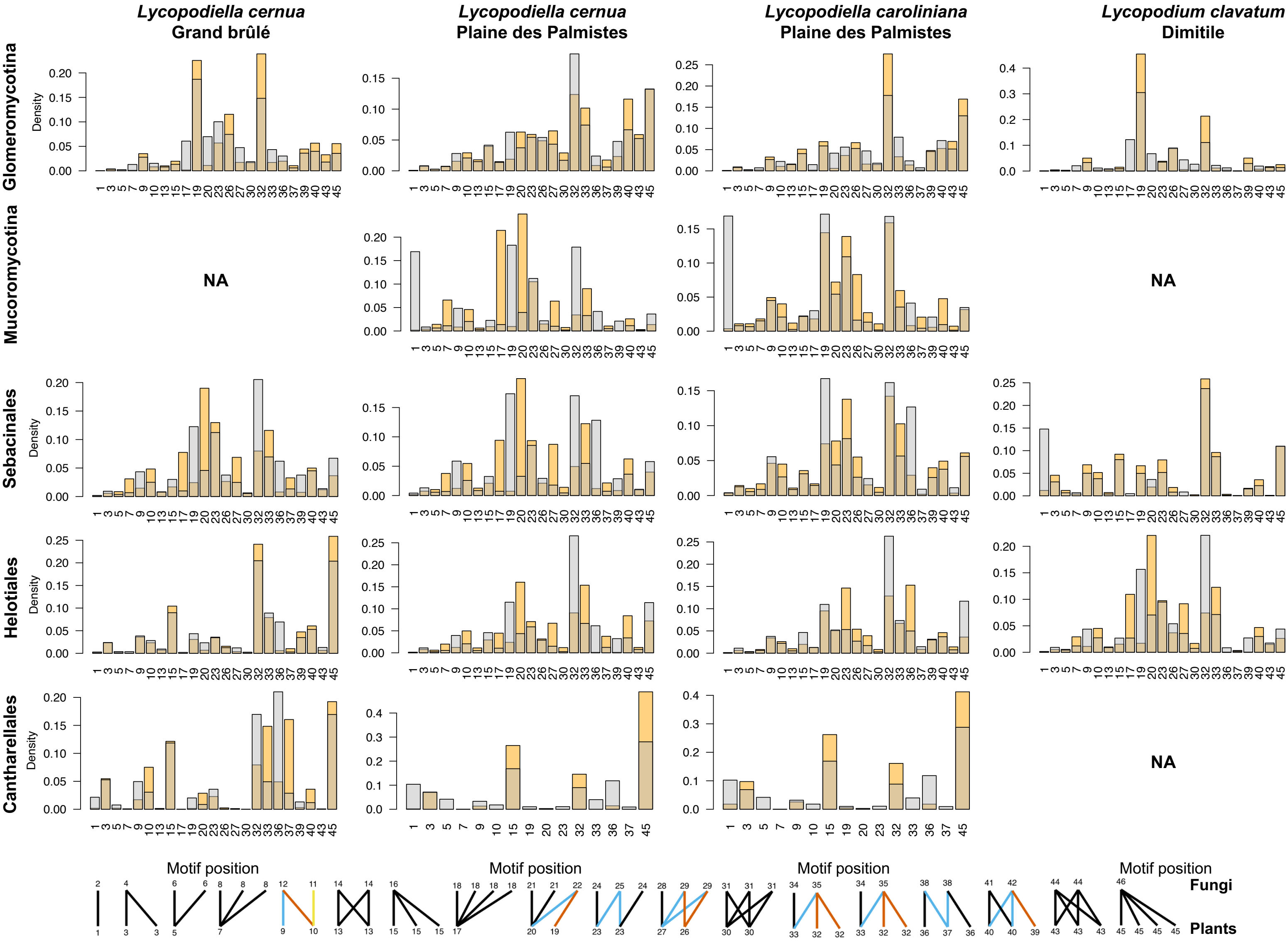

**Supplementary Figure 18: Representation of the plant-fungus networks with fungi taxonomically identified at the genus level:**

Network representation at the species level of the different plant-fungus networks (*Glomeromycotina, Mucoromycotina, Sebacinales,* or *Helotiales*) in each sampled community (Grand brûlé, Plaine-des-Palmistes, or Dimitile). Colored round nodes represent plant species (colors separate the main plant taxonomic groups) and color squared nodes correspond to fungal OTUs (colors indicate the fungal genera and families). Grey links represent plant-fungus interactions and their widths are proportional to interaction abundances. Networks were visualized using the *igraph* R-package.

*(a) Glomeromycotina*

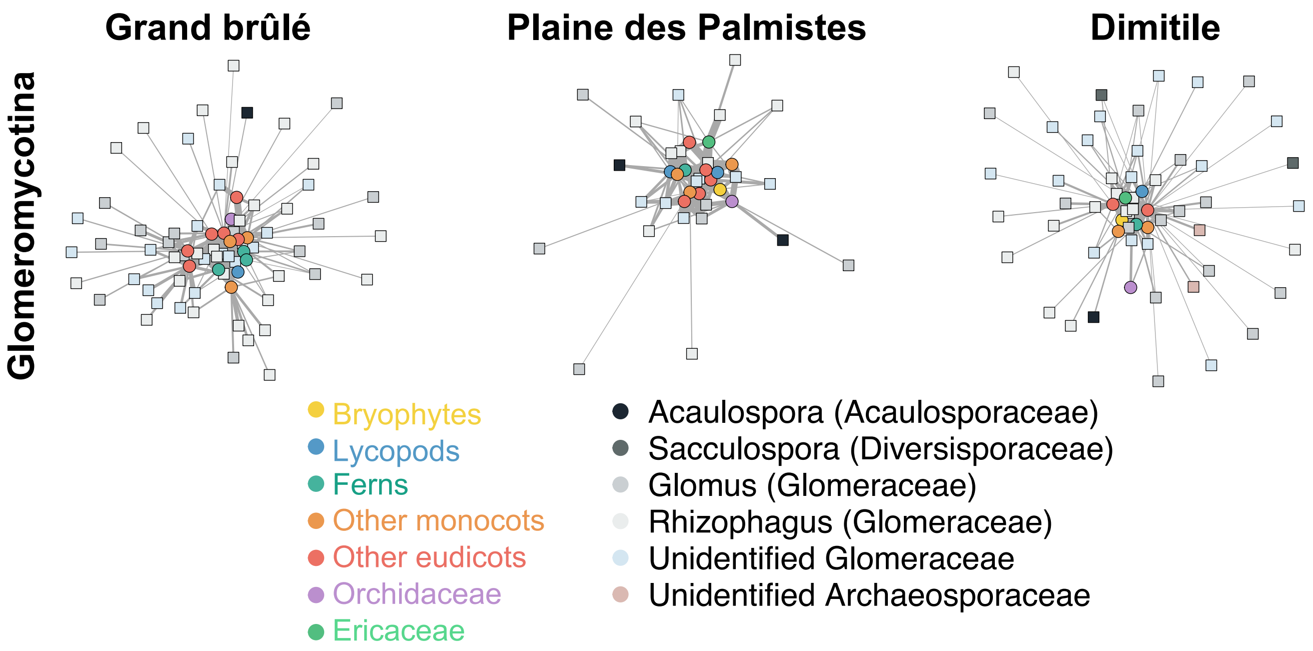

*(b) Mucoromycotina*

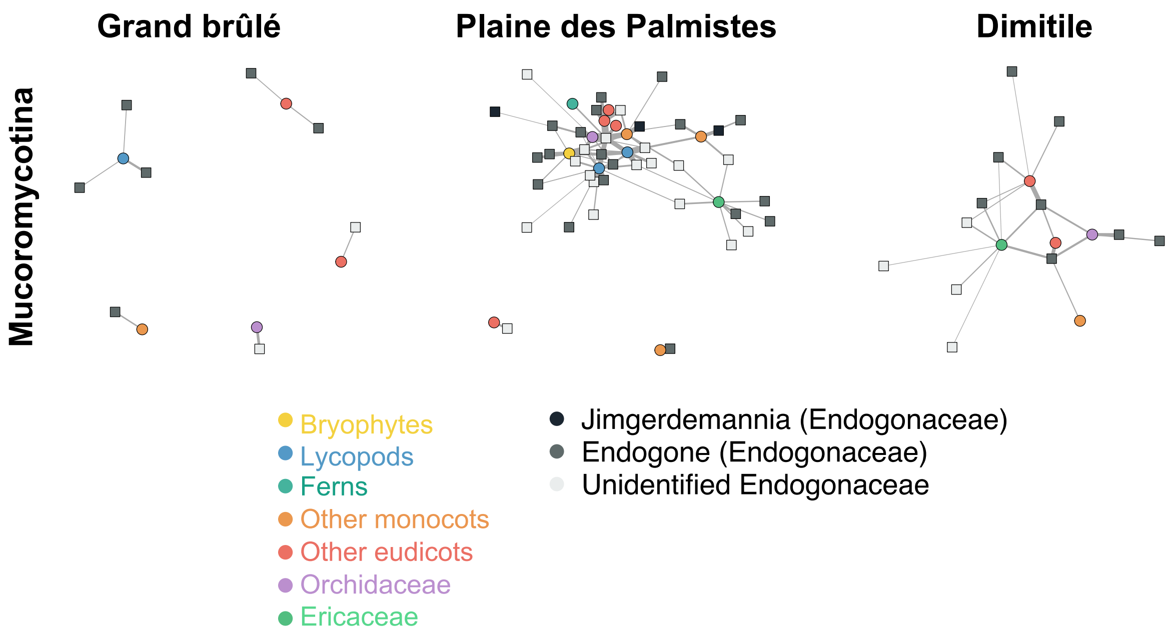


*(c) Sebacinales*

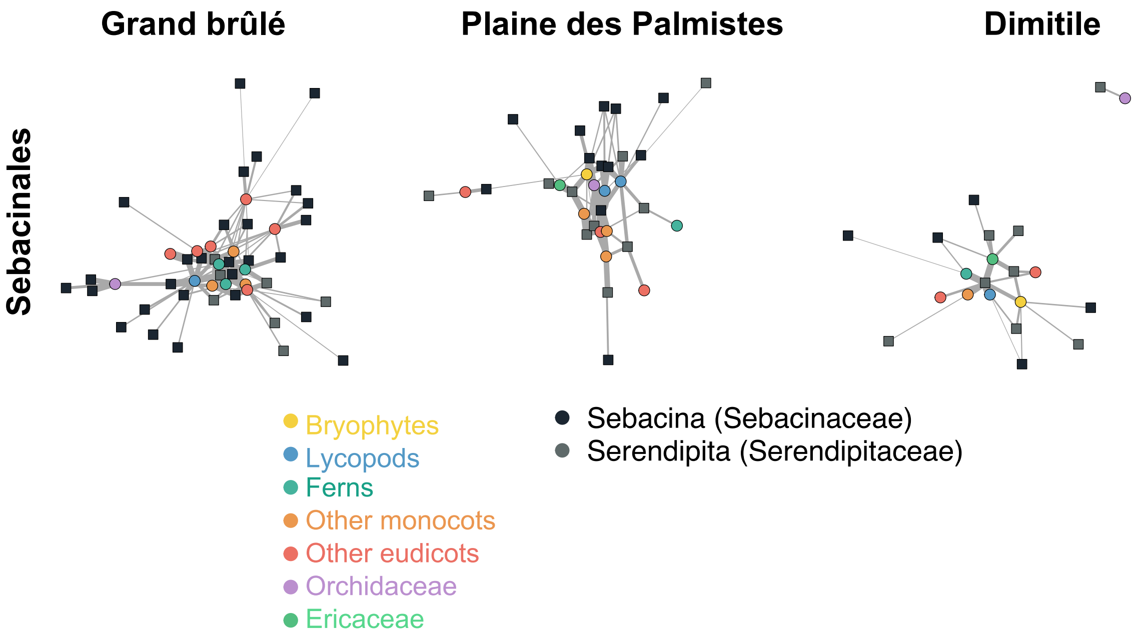

*(d) Helotiales*

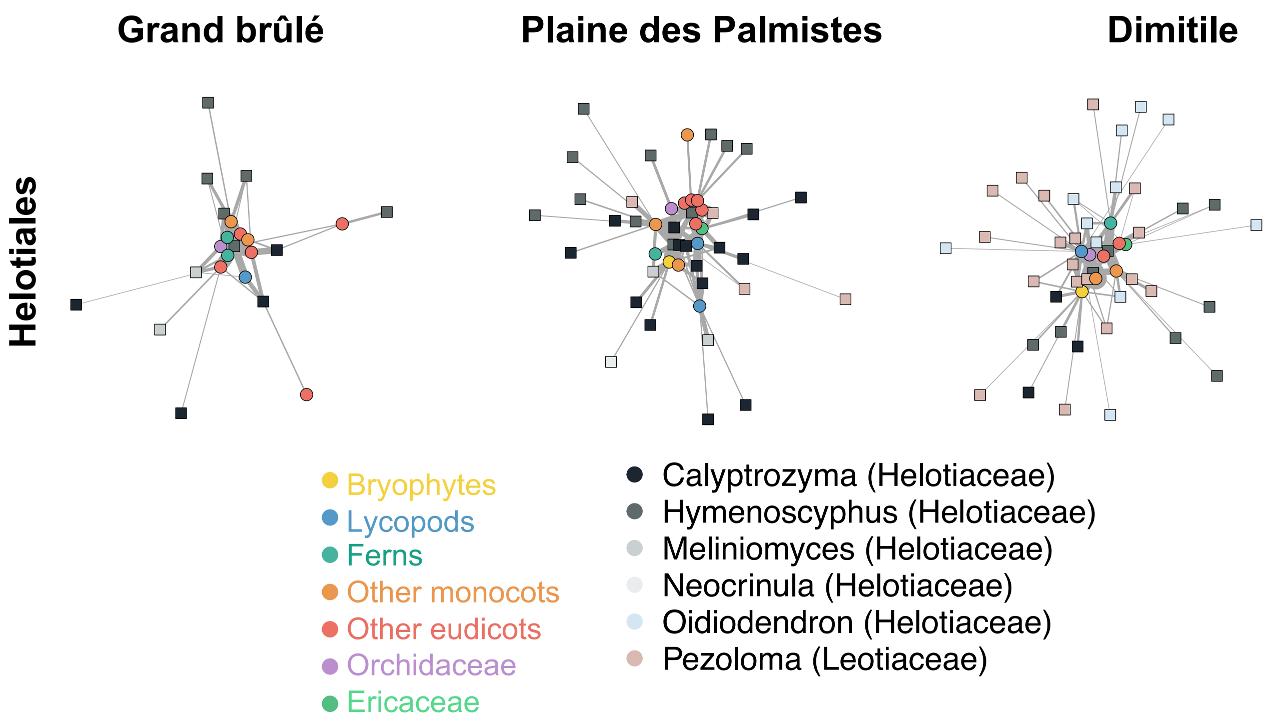


*(e) Cantharellales*

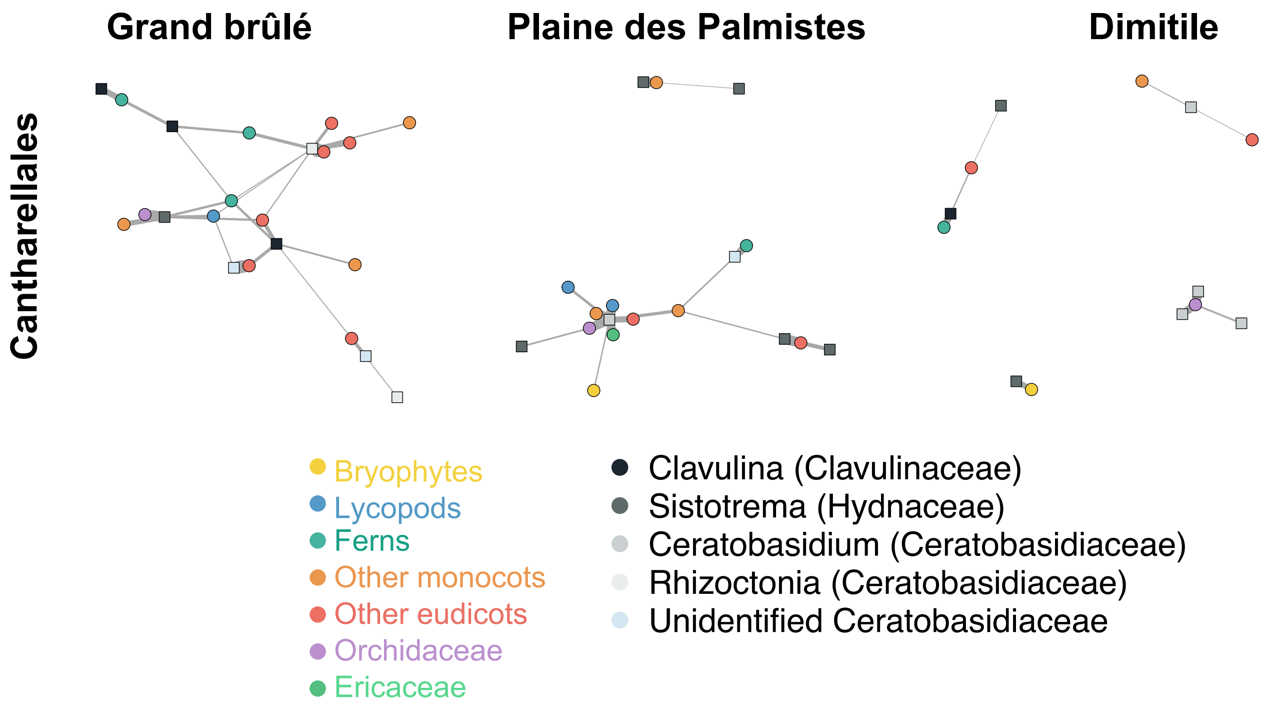
